# SEACells: Inference of transcriptional and epigenomic cellular states from single-cell genomics data

**DOI:** 10.1101/2022.04.02.486748

**Authors:** Sitara Persad, Zi-Ning Choo, Christine Dien, Ignas Masilionis, Ronan Chaligné, Tal Nawy, Chrysothemis C Brown, Itsik Pe’er, Manu Setty, Dana Pe’er

## Abstract

Metacells are cell groupings derived from single-cell sequencing data that represent highly granular, distinct cell states. Here, we present single-cell aggregation of cell-states (SEACells), an algorithm for identifying metacells; overcoming the sparsity of single-cell data, while retaining heterogeneity obscured by traditional cell clustering. SEACells outperforms existing algorithms in identifying accurate, compact, and well-separated metacells in both RNA and ATAC modalities across datasets with discrete cell types and continuous trajectories. We demonstrate the use of SEACells to improve gene-peak associations, compute ATAC gene scores and measure gene accessibility in each metacell. Metacell-level analysis scales to large datasets and are particularly well suited for patient cohorts, including facilitation of data integration. We use our metacells to reveal expression dynamics and gradual reconfiguration of the chromatin landscape during hematopoietic differentiation, and to uniquely identify CD4 T cell differentiation and activation states associated with disease onset and severity in a COVID-19 patient cohort.

## Introduction

A fundamental disconnect currently exists between the cellular resolution of single-cell genomics data and the cluster-level resolution of analysis, which has dramatically limited these technologies in fulfilling their potential for biomedical research. A dataset that harbors tens of thousands of cells is typically summarized as a handful of clusters in order to overcome the noise and sparsity inherent to single-cell data. Sparsity is particularly acute in single-cell assay for transposase-accessible chromatin sequencing (scATAC-seq) data, which only captures the trinary zygosity states at a few thousand of the hundreds of thousands of open chromatin regions in a cell, making it impossible to infer regulation at the single-cell level (**Supplementary Fig. 1**). While single-cell RNA sequencing (scRNA-seq) data is not as sparse, projects such as the Human Cell Atlas^1^ and Human Tumor Atlas Network^2^ are scaling to millions of cells, causing even routine dimensionality reduction and visualization tasks to struggle with computational complexity and confounding by sample-level batch effects. As a result, large scRNA-seq datasets are also best analyzed at the cluster level.

Cluster-level analysis has led to important biological discoveries. However, a typical cluster is not homogenous; structured variability in gene programs within clusters suggests underlying cell-state heterogeneity (**Fig. 1A,B**). For example, cells within T-cell clusters can exhibit different levels of activation and metabolic activity^3^. Moreover, single-cell data has been shown to reside on a continuum^4–7^. For instance, binning the expression of *GATA2*, a driver of erythroid fate, in one cluster of erythroid precursor cells^8^ by developmental progression demonstrates gradual cell-state changes within each bin during human hematopoiesis that is accompanied by epigenomic variation (**Fig. 1C,D**). The accessibility landscape of the *GATA2* locus suggests that its expression dynamics are enabled by gradual opening of regulatory elements (**Fig. 1D** and **Supplementary Fig. 1B**). Such dynamics are lost in any discrete cluster-level analysis.

**Fig. 1:**
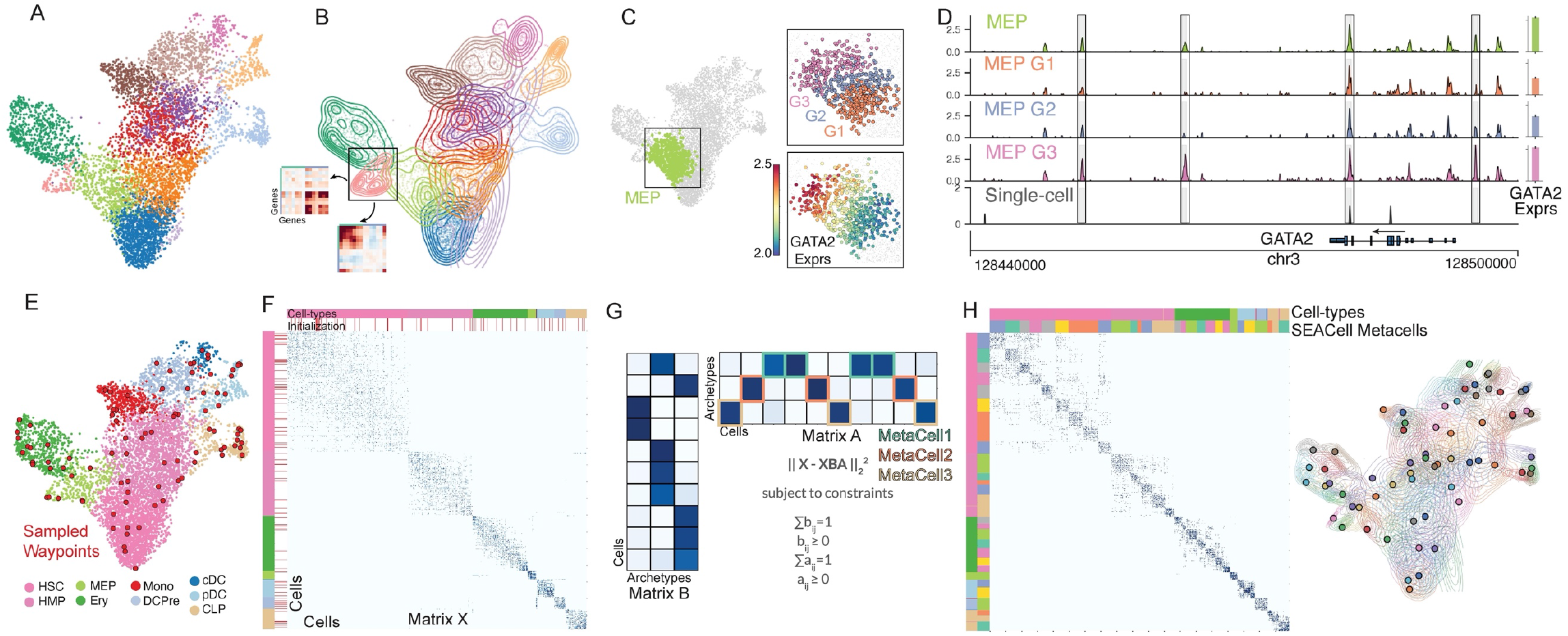
Overview of the SEACells algorithm for cell-state identification from single-cell data. A. scRNA-seq UMAP (uniform manifold approximation and projection) of 6800 CD34+ hematopoietic stem and progenitor stem cells. Cells colored by cluster. B. Contour plots of each cluster highlight density and indicate the presence of multiple cell-states within each cluster. Inset, gene-gene covariance matrices reveal that each state is accompanied by distinct gene expression programs. C. Left, UMAP with megakaryocyte-erythroid progenitor (MEP) cluster highlighted. Right, MEP cluster is divided into three equal-sized bins based on developmental progression (top), reflecting imputed expression of *GATA2* (known driver of MEP lineage) (bottom). D. Coverage plots showing *GATA2* accessibility in all MEPs (top), a single MEP cell (bottom) and in the three bins in (C). Right, expression of *GATA2* in corresponding cells. Highlighted peaks demonstrate how accessibility dynamics track with expression dynamics. Information about dynamics is masked at cluster level, whereas peak identification in single cells is too noisy. E. UMAP as in (A), colored by cell-type. The SEACells algorithm for metacell identification is initialized by waypoints (large red circles), a subset of cells sampled to uniformly cover the phenotypic landscape. F. Heatmap showing the cell-to-cell affinity matrix computed using an adaptive Gaussian kernel. Cells are sorted by cell-types (top annotation row). Second annotation row shows the SEACells initialization. G. Schematic of archetypal analysis. The data matrix is decomposed into two the archetype matrix B and embedding matrix A. Metacell membership is identified based on column-wise maximal values across the matrix A. H. Left, cell-cell affinity matrix from (F), but ordered by metacell assignment. Right, contour plot overlying UMAP from (E), highlighting the distribution of metacells; cells and contours colored by metacell assignment.

The concept of metacells^9^—groups of cells that represents distinct, highly granular cell states, whereby within-metacell variation is due to technical rather than biological sources—was proposed as a way to maintain statistical utility while maximizing effective data resolution^9^. Metacells are far more granular than clusters, and are optimized for homogeneity within cell groups, rather than for separation between clusters. However, existing approaches^9, 10^ fail on scATAC-seq data, aggressively cull outliers (particularly inappropriate for disease studies, which are often driven by rare cell populations), and are poorly distributed across the phenotypic space. Consequently, metacells have not been routinely used in single-cell analysis, and scATAC-seq data has remained heavily underutilized.

Here, we present SEACells, a graph-based algorithm that uses iterative archetypal analysis to compute metacells. We evaluate our approach on peripheral blood data with discrete, well-separated cell-types, and on CD34^+^ hematopoietic stem and progenitor cell (HSPC) data from human bone marrow with continuous gradients underlying early decisions in hematopoiesis. SEACells metacells provide robust, comprehensive characterizations of scRNA-seq cell states, including gene-gene relationships representative of each state^11^; and they successfully describe chromatin cell states at resolutions that enable the inference of regulatory elements underlying gene expression. Our metacells achieve a sweet spot between signal aggregation and cellular resolution, and they capture cell-states across the phenotypic spectrum, including rare states. They are also computationally tractable, enabling powerful downstream analysis of large-scale datasets. We show that our metacells overcome technical batch effects to allow for superior data integration when matching metacells across samples in a cohort. We use SEACells to learn dynamics of expression and accessibility during hematopoietic differentiation and temporal dynamics of T-cell response during COVID-19 infection, biological insights that are missed by single-cell and cluster-level analysis. SEACells provides a powerful toolkit for gene regulatory inference from scATAC-seq data and a tractable solution for integration of large cohort based single-cell data.

## Results

### The SEACells algorithm identifies metacells across the phenotypic manifold

SEACells seeks to aggregate single cells into metacells that represent distinct cellular states, in a manner agnostic to data modality. Using a count matrix as input, it provides per-cell weights for each metacell, per-cell hard assignments to each metacell, and the aggregated counts for each metacell as output. Moreover, an explicit design goal of our approach is to capture the full spectrum of cell states in the data, including rarer states. We base SEACells on a few key assumptions: 1) single-cell profiling data can be approximated by a lower-dimensional manifold (phenotypic manifold), 2) much of the observed variability across cells is due to incomplete sampling; molecular profiles only represent a small fraction of transcripts in each cell, and 3) most cells can be assigned to a finite set of cell states, each characterized by a distinct combination of active gene programs. Biology is modular—each cell needs to perform a distinct set of tasks and each task requires the activity of a relevant gene program, creating constraints and structure. Moreover, many gene programs interact through feedback and feedforward regulation, further constraining the system.

SEACells takes advantage of graph-based algorithms for manifold learning that have been proven to capture the cell state landscape in single-cell genomics data faithfully and robustly^4, 6, 7, 12–15^. The algorithm first constructs a nearest-neighbor graph to represent the phenotypic manifold. It then applies archetype analysis^16, 17^ to iteratively refine metacells, and finally aggregates counts into a set of output metacells. Manifold construction is tailored to each data modality, at which point the algorithm can proceed in data-type agnostic fashion (**Supplementary Fig. 2**). We use CD34+ cells from early human hematopoiesis to demonstrate our method (**Fig. 1**). For initializing the metacell search, we utilize our max-min sampling approach^5^. Max-min sampling identifies a set of representative cell states that are distributed uniformly across the phenotypic manifold (**Fig. 1E**), and it is particularly adept at dealing with density differences, thus ensuring the capture of rare states. These sampled cell states are waypoints (multiple per cell type) that define clear structure in the neighbor graph; however, the cell-states themselves remain somewhat diffuse (**Fig. 1F**).

To refine metacells, we employ archetypal analysis^16^, a robust, linear matrix decomposition approach shown to optimally capture manifold structure and to identify cell states representing characteristic biological processes and tasks (**Fig. 1G**, Methods). Although archetype analysis is linear in nature, applying it to the neighbor-graph-defined adjacency matrix enables it to capture the non-linear structure of the manifold. Archetypal analysis finds the set of archetypes that optimally reconstruct the data matrix, while constraining them to reside within the phenotypic manifold, focusing the computation on the strongest axes of variation. The archetype procedure partitions the data in such a way that the cell-cell similarity matrix has tight block structure along the diagonal, which best represents distinct cell states in the data (**Fig. 1H**). While early human hematopoiesis is largely defined by differentiation trajectories, we note that the cells underlying these relatively continuous processes are well-represented by a set of distinct metacells.

### SEACells metacells represent accurate and robust cell states in diverse data

We first evaluated SEACells performance on a public multiome (simultaneous single-cell RNA-seq and ATAC-seq) dataset of peripheral blood mononuclear cells (PBMCs), as a well-studied system with distinct cell populations. We found that SEACells metacells are comprehensive and well-distributed among cell types in both RNA and ATAC data (**Fig. 2A,B**). Metacells of both modalities also exhibit a high degree of cell-type purity, which agrees well with the fact that PBMCs are made up of discrete, mature cell types (**Fig. 2A,B**, Methods). Further, reciprocal projections of RNA and ATAC metacells onto each other demonstrate that metacells of different modalities are highly concordant (**Supplementary Fig. 3A,B**, Methods).

**Fig. 2:**
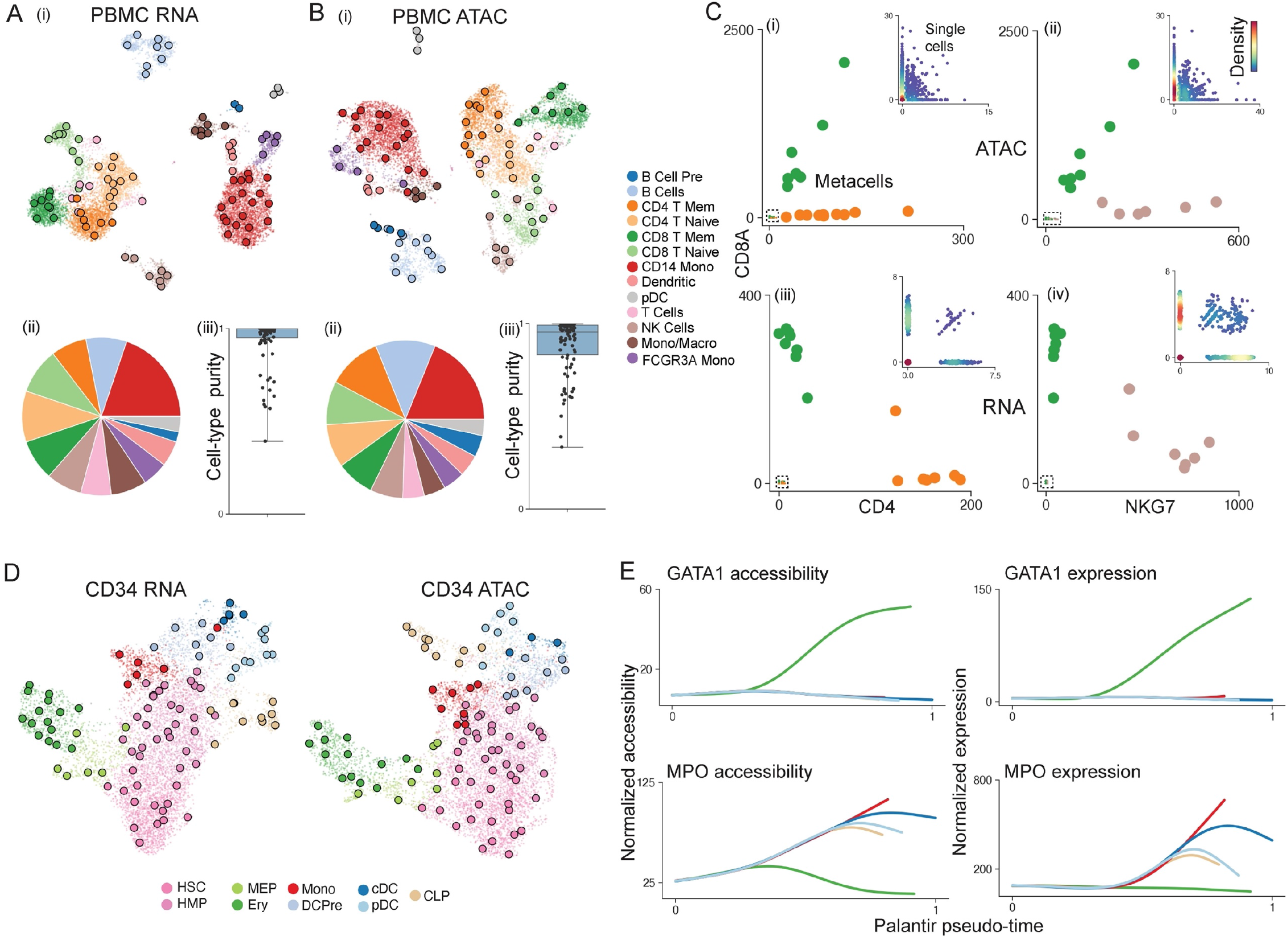
SEACells metacells accurately identify cell states and outperform competing approaches. A. (i) UMAP of human PBMCs derived from RNA data of a multiome dataset, highlighting cell types and SEACell metacells. (ii) Distribution of metacells per cell type for the RNA modality. (iii) box plot of distribution of cell-type purity (frequency of the most represented cell-type in each metacell). High purity represents a more accurate metacell. B. As in (A), using ATAC data from the PBMC multiome dataset. C. Metacell accessibility (i) and expression (iii) of *CD4* and *CD8A* accurately distinguishes CD8 (green) and CD4 (orange) T-cell compartments. Metacell accessibility (ii) and expression (iv) of *NKG7* and *CD8A* distinguish NK (pink) and CD8 (green) T-cells. Insets, corresponding single-cell accessibility is too sparse to achieve the same distinction. D. UMAPs of CD34+ hematopoietic stem and progenitor stem cells highlighting cell types and the SEACell metacells independently constructed from RNA (left) and ATAC (right) data. E. Accessibility (left) and expression (right) of *GATA1* (erythroid factor) and *MPO* (myeloid factor) along the Palantir pseudotime axis representing hematopoietic differentiation. Palantir was run on RNA aggregates using ATAC metacells and accurately recapitulates dynamics.

The key advantage of metacells is that they help to overcome data sparsity, which is extreme in scATAC-seq. We found that each SEACells metacell provides a more complete molecular characterization than individual cells—for example, by revealing accessibility at known marker genes for major cell types (**Supplementary Fig. 3C,D**). Accessibility and expression of *CD4* and *CD8* from metacells, but not most individual cells, can accurately distinguish the two T-cell subsets, and *NKG7* and *CD8A* are sufficient to distinguish NK and CD8 T-cell populations (**Fig. 2C**). Metacells thus comprise pure cell types expressing expected markers in this data; they are granular enough to distinguish states within cell types; and they can be queried with classical immune markers.

We next tested whether SEACells can accurately determine metacells in continuous differentiation trajectories during the earliest decisions in human hematopoiesis, when cells are not yet well-separated. We collected a single-cell multiome dataset of 6,800 hematopoietic stem and progenitor cells (HSPCs) from healthy bone marrow sorted for pan-HSPC marker CD34 (Methods). Similar to PBMCs, we found that metacells are well-distributed across all bone marrow cell types and span the RNA and ATAC phenotypic manifolds (**Fig. 2D**). To determine whether metacell resolution is sufficient to accurately recover gene expression dynamics that are lost in clustering, we applied the Palantir trajectory algorithm^5^ directly to metacells. Palantir could indeed recover the known expression and accessibility dynamics of key hematopoietic genes (**Supplementary Fig. 4**). As a further challenge, we ran Palantir on aggregated RNA from metacells computed on the ATAC modality, since the sparsity of scATAC-seq data renders cell-state identification much more difficult (**Fig. 2E** and **Supplementary Fig. 4**). The fidelity of captured gene trends reinforces that SEACells metacells overcome sparsity but retain cell-state heterogeneity and dynamics in systems with continuous state transitions.

We also used this dataset to assess the robustness of SEACells (Methods). First, we verified that SEACells is robust to different initializations by observing consistency in the metacells identified in different runs (**Supplementary Fig. 5A**). Specifically, we jointly embedded metacells from two initializations using diffusion components and tested the percentage of mutually nearest neighboring cell-states between the initializations that connect states of the same cell type (Methods). SEACells metacells are extremely robust to different initializations in both RNA and ATAC modalities (**Supplementary Fig. 5B**). We then demonstrated that specifying different numbers of metacells still recovers consistent states (**Supplementary Fig. 5C,D**). Our results confirm that SEACells calls metacells robustly in both RNA and ATAC modalities.

Another key performance metric is the ability to capture rare cell states. SEACells were able to accurately recover rare cell-types such as pDCs and B-cell precursors in the PBMC RNA and ATAC modalities (**Fig. 2A,B**). To further test the ability of SEACells to identify rare intermediate cell states in continuous trajectories, we generated a second multiome dataset of T-cell depleted bone marrow cells representing the full span of human hematopoiesis (**Supplementary Fig. 6A,F**). As expected, we observed an extraordinary diversity of densities across the phenotypic manifold, with low-density regions representing rare intermediate cell types (**Supplementary Fig. 6B,C,G,H**). SEACells accurately identified metacells in these low-density regions (**Supplementary Fig. 6D,I**), otherwise masked by clustering (**Supplementary Fig. 6E,J**), demonstrating that the algorithm can recover rare cell types and cell states in both discrete and continuous datasets across RNA and ATAC data modalities.

### SEACells empowers gene regulatory inference

Peaks of ATAC-seq read counts represent open chromatin regions, and gene regulation can be inferred by identifying putative transcription factor (TF) binding motifs within these accessible regions. Single-cell ATAC-seq provides many observations (cells) with the potential to infer more complex gene regulatory models—using trajectory inference, for example—at fine resolution^18–20^. However, the sparsity of scATAC-seq data has severely restricted its utility, as analysis typically occurs at the resolution of clusters. We surmised that SEACells metacells provide an ideal trade-off between fine resolution and sufficient coverage to overcome sparsity for diverse gene regulatory inference tasks.

A typical SEACells metacell contains 1.2 million reads, a large improvement over the 25,000 reads in an individual cell, but still far fewer than the 50 million reads processed in a typical bulk sample. The first step towards building a SEACells regulatory toolbox is thus to improve the signal-to-noise ratio in ATAC peak calling. We take advantage of the characteristic ATAC-seq fragment length distribution (**Supplementary Fig. 7A**)^21^, in which the first mode represents nucleosome-free (NFR) fragments likely enriched for TF binding events, and the second mode represents nucleosomes. Since nucleosomes occupy a broader region of the genome compared to TFs, we observed that they cause many false positive motif calls; peaks called using all fragments thus result in poorly resolved regulatory elements (**Supplementary Fig. 7B**). Using only NFR fragments identifies fewer peaks; however, we found that these peaks are enriched for TF-bound open chromatin, including a large number of additional peaks that were obscured when considering all fragments (**Supplementary Fig. 7B,C**). Regulatory element identification thus benefits from using NFR fragments rather than all fragments (**Supplementary Fig. 7C**).

The next task in gene regulatory inference is to associate each gene with the specific elements that regulate it. Only a subset of local open-chromatin peaks is relevant to the expression of a gene, since ATAC-seq profiles reflect regulatory elements as well as structural factors such as CTCF^22^. The correlation between accessibility peaks and expression across cells has been used to predict the set of cis elements that regulate each gene, using either multiome^18^ or separate scRNA-seq and scATAC-seq data that is integrated^23^. Data sparsity precludes robust linkage of regulatory elements with genes at the single-cell level. Using SEACell metacells from the CD34^+^ bone marrow dataset, we computed correlations between gene expression and NFR peak accessibility for each peak within +/-100 kb of each gene in a core hematopoietic gene set^5^. For all core genes, accessibility of the most correlated peak using ATAC metacells faithfully tracks with gene expression, representing a substantial improvement over correlations identified from the same data at the single-cell level (**Fig. 3A** and **Supplementary Fig. 8**). For example, the correlation between peak accessibility and expression in metacells for key erythroid lineage regulator *TAL1* is 0.82, and cells on the erythroid trajectory exhibit the highest values, whereas the correlation is 0.03 at single-cell level, with no distinction among erythroid cells (**Fig. 3A**).

**Fig. 3:**
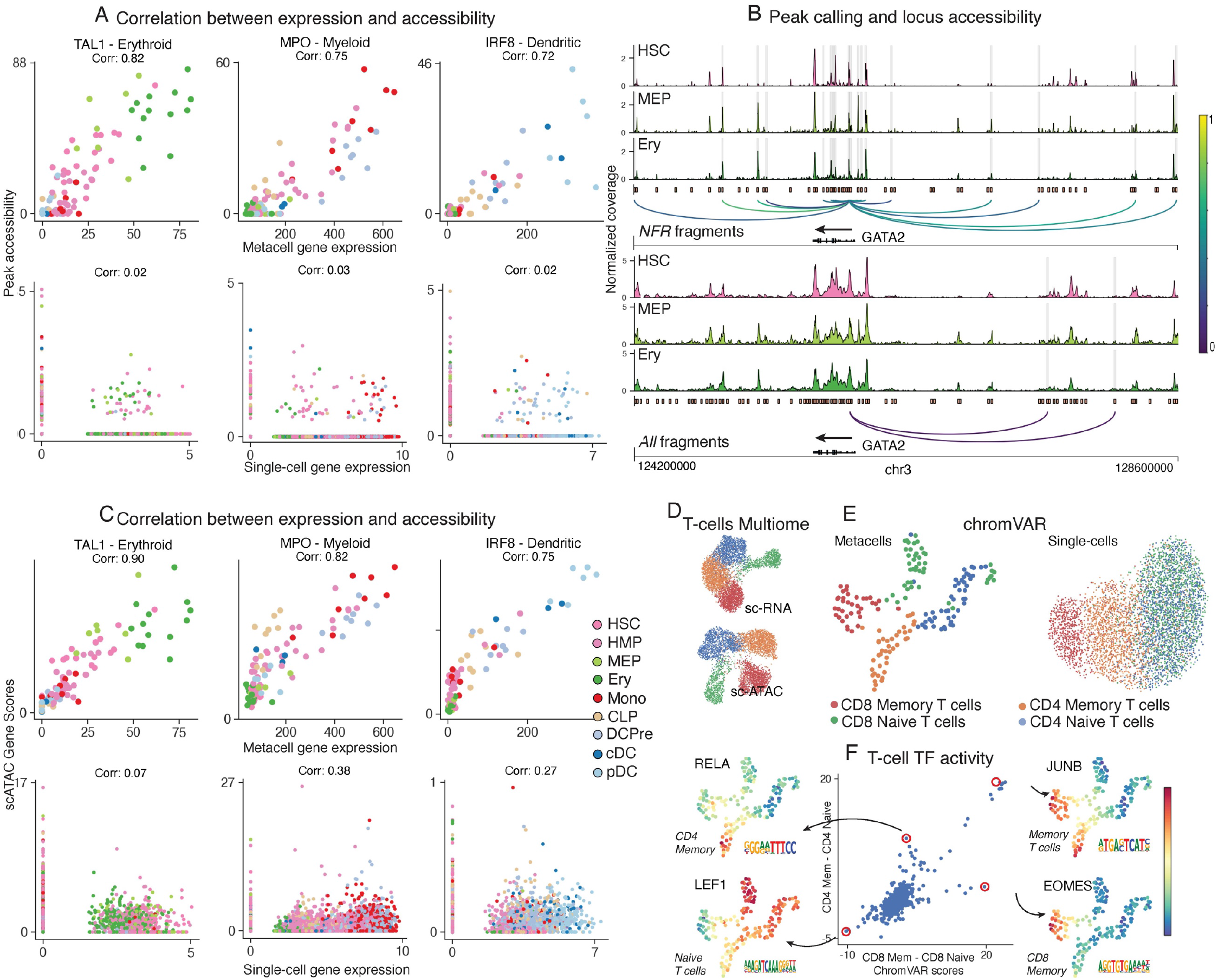
SEACells empowers a gene regulatory toolkit. A. Spearman correlation between ATAC metacell-aggregated (top) or single-cell (bottom) gene expression and accessibility of the most correlated peak in *TAL1* (erythroid), *MPO* (myeloid) and *IRF* (dendritic) marker genes, computed on CD34+ multiome data. Each metacell and single cell is colored based on cell type. B. Accessibility landscape of erythroid factor *GATA2* in hematopoietic stem cells (HSC), myeloid-erythroid progenitors (MEP) and erythroid cells (Ery) using NFR (top) or all ATAC (bottom) fragments. Restricting chromatin accessibility analysis to NFR fragments improves peak resolution and the association of regulatory elements with genes. Arcs are colored by peak-gene Spearman correlation (color values between 0 and 1 at right), determined using SEACells ATAC metacells. Highlighted peaks correlate significantly with *GATA2* expression. C. Relationships between metacell-aggregated (top) and single-cell (bottom) gene expression and ATAC gene scores for *TAL1*, *MPO* and *IRF*. Spearman correlations (Corr) computed using the CD34+ multiome data. Metacell gene scores were computed by aggregating peaks that correlate significantly with expression (e.g. Fig. 3B). Gene scores for single-cell data were computed using ArchR. D. RNA and ATAC UMAPs of the T-cell subset from the PBMC multiome dataset. E UMAPs derived from chromVAR scores computed using single cells or metacell aggregates. All peaks were used for chromVAR analysis. Metacell chromVAR scores accurately recapitulate differences between T-cell subsets, whereas single-cell chromVAR scores fail to distinguish CD4 and CD8 T-cells. F. chromVAR score distributions can be used to identify key TFs that define different T-cell compartments. Each dot represents a TF. X-axis shows the difference between SEACells metacell chromVAR scores between the two CD8 compartments. Y-axis shows the difference between SEACells metacell chromVAR scores between the two CD4 compartments

To build a comprehensive map of regulatory elements, we identified all peaks significantly correlated with a gene compared to GC-content-matched peaks sampled from the data as an empirical background set^18^ (Methods). For the key erythroid factor *GATA2*, single-cell data only recovers 2 of 11 associations detected using metacells (**Fig. 3B**). To systematically explore the accuracy of predicted peak-gene associations, we computed gene scores^23^ by aggregating the accessibility of all significantly correlated peaks and comparing them to gene expression (Methods). SEACells gene scores are substantially better correlated and outperform aggregating single cells across all correlated peaks (correlation of 0.05, compared to 0.88 using SEACells) (**Fig. 3C** and **Supplementary Fig. 9**). This improvement was consistent across the core hematopoietic genes, as well as all genes with expression in at least 10 cells (**Fig. 3C** and **Supplementary Fig. 9C)**. SEACells metacells thus clearly identify cis elements that are significantly correlated with gene expression and likely regulate the corresponding gene, enabling complex gene regulatory modeling.

To overcome data sparsity in scATAC-seq, genome-wide information is often aggregated for all peaks associated with a particular TF and summarized as a TF activity score. chromVAR^24^ is a widely used tool for predicting transcription factor activity from scATAC-seq data. It provides a per-cell deviation score for a TF by computing whether the peaks predicted to contain its binding motif have greater accessibility compared to a GC-matched background peak set^24^. To demonstrate that metacell resolution can substantially improve TF activity inference, especially in more complex regulatory landscapes, we used T-cell subsets, which reside on a relatively continuous landscape^11, 25, 26^ driven by competing feedback loops. We determined chromVAR scores for all T-cell subsets (CD4 naive and memory, CD8 naive and memory) using the PBMC multiome dataset (**Fig. 3D**). chromVAR scores provide an alternate representation of the ATAC data, useful for all downstream analyses including clustering and visualization. Indeed, chromVAR scores using metacells accurately recovered the distinction between different T cell subsets, whereas single-cell chromVAR scores failed to distinguish CD8 and CD4 (**Fig. 3E**). We identified several known compartment-specific TFs that likely drive these T-cell states, including *JUNB*, a factor active in CD4 and CD8 memory T-cells^27^; *LEF1*, active in CD4 and CD8 naive T-cells^28^; *EOMES*, a key regulator of CD8 memory cells^29^ and *RELA*, a factor necessary for CD4 memory cell function^30^ (**Fig. 3F**) Single-cell chromVAR scores of these factors failed to distinguish the same populations (**Supplementary Fig. 10**). In summary, SEACells substantially improves the regulatory toolkit for analyzing and interpreting scATAC-seq data, including widely used tools such as chromVAR.

### SEACells outperforms metacell approaches for RNA and is the only approach amenable to ATAC data

Baran and colleagues^9^ introduced and effectively articulated the metacell concept. Their MetaCell algorithm was demonstrated on healthy systems and designed around MARS-seq data, which has a high instance of extreme values^31^, so it culls outliers aggressively. However, rare cell populations often drive biology, especially in contexts such as cancer and regeneration. We found that on lung adenocarcinoma scRNA-seq data^32^, MetaCell throws out more than one-third of all cells (**Supplementary Fig. 11A,B**). Another approach, Super-cells^10^, is effectively a very fine clustering strategy that adapts widely-used community detection algorithms to generate a large number of small clusters. We determined that SEACells is superior to both approaches on RNA, and it is the only method that works on scATAC-seq data (**Fig. 4**).

**Fig. 4:**
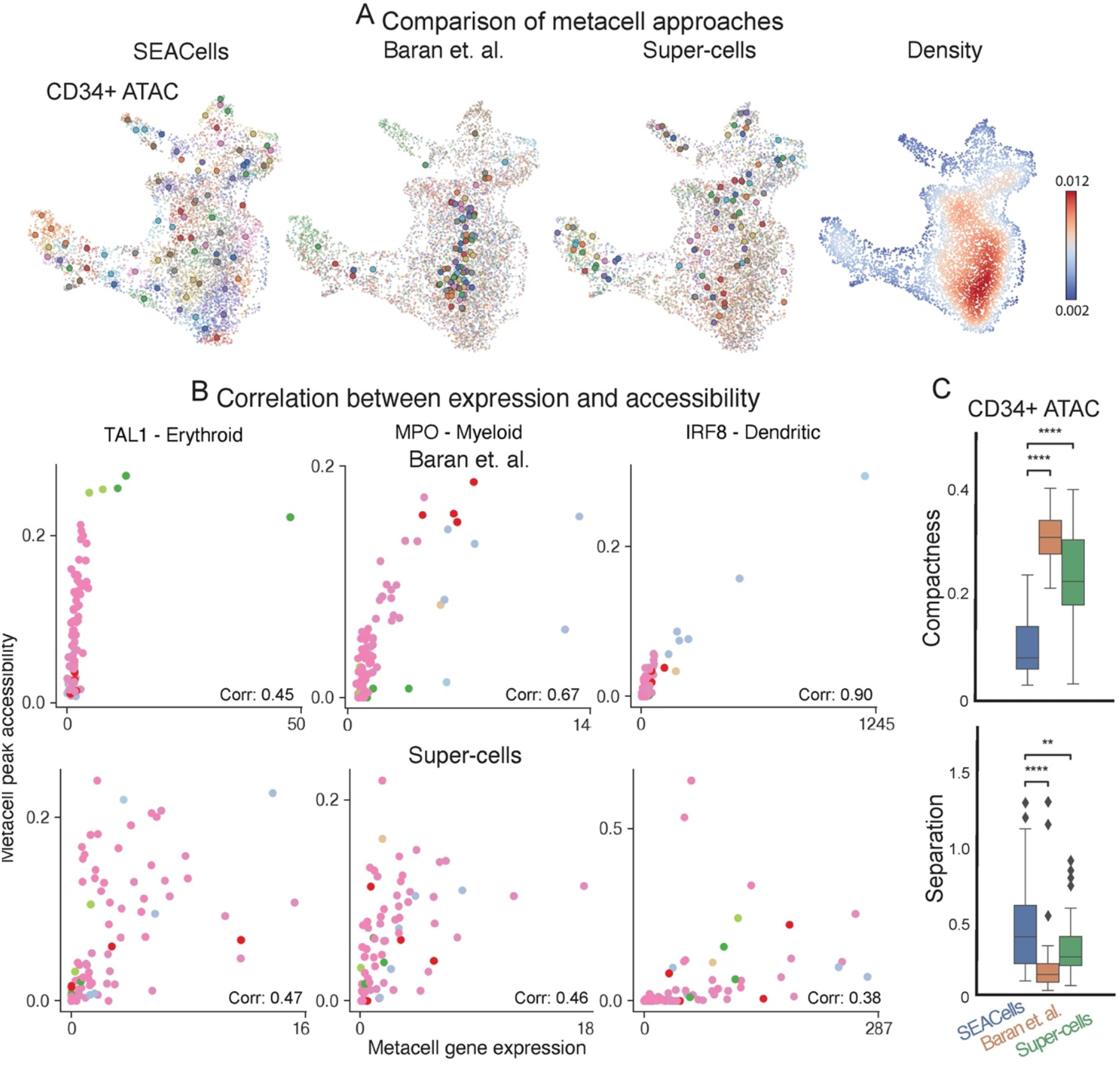
SEACells outperforms existing methods in cell-state representation and correlation of expression and accessibility. A. ATAC modality UMAPs of CD34^+^ bone marrow (as in Fig. 2D), colored by metacell aggregates identified by the specified method or colored by cell density. Dots, cells; circles, metacells. B. Pearson correlation between metacell-aggregated gene expression and accessibility of the most correlated peak in *TAL1* (erythroid gene), *MPO* (myeloid gene) and *IRF* (dendritic gene) using the CD34+ bone marrow ATAC metacells called by MetaCell (top) or Super-cells (bottom). C. Top: Metacell compactness (average diffusion component standard deviation; Methods) measured in the ATAC modality of CD34^+^ bone marrow multiome data. A lower score indicates more compact metacells. Bottom: Metacell separation (distance between nearest metacell neighbor in diffusion space; Methods) measured in the ATAC modality of CD34^+^ bone marrow multiome data. Greater separation indicates better performance. Comparisons were carried out on all metacells, or metacells in low-density or high-density regions. Wilcoxon rank-sum test; ns: *P* > 0.05, * 0.01 < *P* < 0.05, ** 0.001 < *P* < 0.01, *** 0.0001 < *P* < 0.0001, **** *P* < 0.0001.

We compared the three algorithms using ATAC and RNA modalities from CD34^+^ bone marrow and PBMC datasets. Since both MetaCell and Super-cells require a gene count matrix, we aggregated peaks in the gene body to derive a count matrix representation for ATAC data. Unlike these methods, SEACells explicitly samples the entire manifold, optimizing the inclusion of cell states distinct from those already detected. Ours was the only algorithm to cover the entire phenotypic landscape (**Fig. 4A** and **Supplementary Fig. 12**). For ATAC in particular, MetaCell and Super-cells neglected the majority of cell states by focusing calls on cell-dense regions. In bone marrow, they failed to represent important populations such as common lymphoid progenitor (CLP) cells, monocytes, and DC subpopulations, and in PBMCs, they failed to identify coherent cell states (**Fig. 4A**). Super-cells severely under sampled metacells in low density regions (**Supplementary Fig. 12A**), failing to accurately recover the distinction between different T-cell states.

By definition, a metacell represents a single biological cell state, meaning that its constituent cells should share the same cell-type label. We evaluated cell type purity in PBMC data, which contains well-separated cell types, and found that SEACells metacells of both modalities show significantly greater purity than metacells from other methods (**Supplementary Fig. 11C**). These differences in performance greatly impact the downstream analysis and interpretability of scATAC-seq data. Peak accessibility and gene expression are also much better correlated in metacells from SEACells (**Fig. 3A** and **Supplementary Fig. 8A**), than MetaCell or Super-cells (**Fig. 4B** and **Supplementary Fig. 13**).

To quantify performance at higher resolution, we defined metrics for metacell compactness and separation. An ideal metacell is compact (exhibits low variance amongst constituent cells) and well-separated (remains distant from cells of a neighboring metacell). We measured compactness by the diffusion component variability of cells within a metacell and separation by the diffusion distance between a metacell and its nearest neighbor (Methods). In both bone marrow and peripheral blood ATAC datasets, we found that SEACells metacells are significantly more compact and better separated than MetaCell and Super-cells metacells (**Fig. 4C** and **Supplementary Fig. 14A,B**), especially in low cell-density regions, which are biologically relevant (**Supplementary Fig. 14A,B**). Super-cells does show marginally better separation in high density regions, however, since it creates a few very large partitions containing hundreds of cells in these regions (**Supplementary Fig. 11D**).

While the different approaches are qualitatively similar using the RNA modality (**Supplementary Fig. 12B,C**), SEACells metacells are significantly more compact than Super-cells (*P* < 1e-5, Wilcoxon rank-sum test) and marginally more compact than Baran et al. metacells (**Supplementary Fig. 14C**). Conversely, SEACells RNA metacells are significantly better separated than Baran et al. metacells (*P* < 1e-2, Wilcoxon rank-sum test), whereas the separation of Super-cells is artificially boosted by the large partition size in high density regions (**Supplementary Fig. 14D**). Similar to ATAC, SEACells metacells have greater cell-type purity in the PBMC RNA data, which comprises distinct cell types (**Supplementary Fig. 11C**).

Collectively, our results show that metacells generated by SEACells better represent the catalog of cell-states present in the data and are more homogenous, compact and well-separated than alternative methods across both RNA and ATAC modalities.

### SEACells reveals gene accessibility dynamics during hematopoietic differentiation

Hematopoietic differentiation is characterized by the upregulation of lineage-defining genes and the downregulation of stem-cell identity genes, driven by precise changes in enhancer accessibility that enable or impede transcription factor binding at these loci (**Fig. 5A**). Both bulk and single-cell ATAC-seq data reveal extensive poising of regulatory elements in stem cells, whereby most enhancers regulating lineage genes are accessible and primed for lineage-specific expression^18, 33, 34^. To demonstrate the potential of SEACells metacells for advanced scATAC-seq analysis, we set out to examine how the primed and permissive epigenomic landscape of hematopoietic stem cells dynamically reconfigures to a landscape with sharply reduced plasticity and developmental potential in differentiated cells.

**Fig 5:**
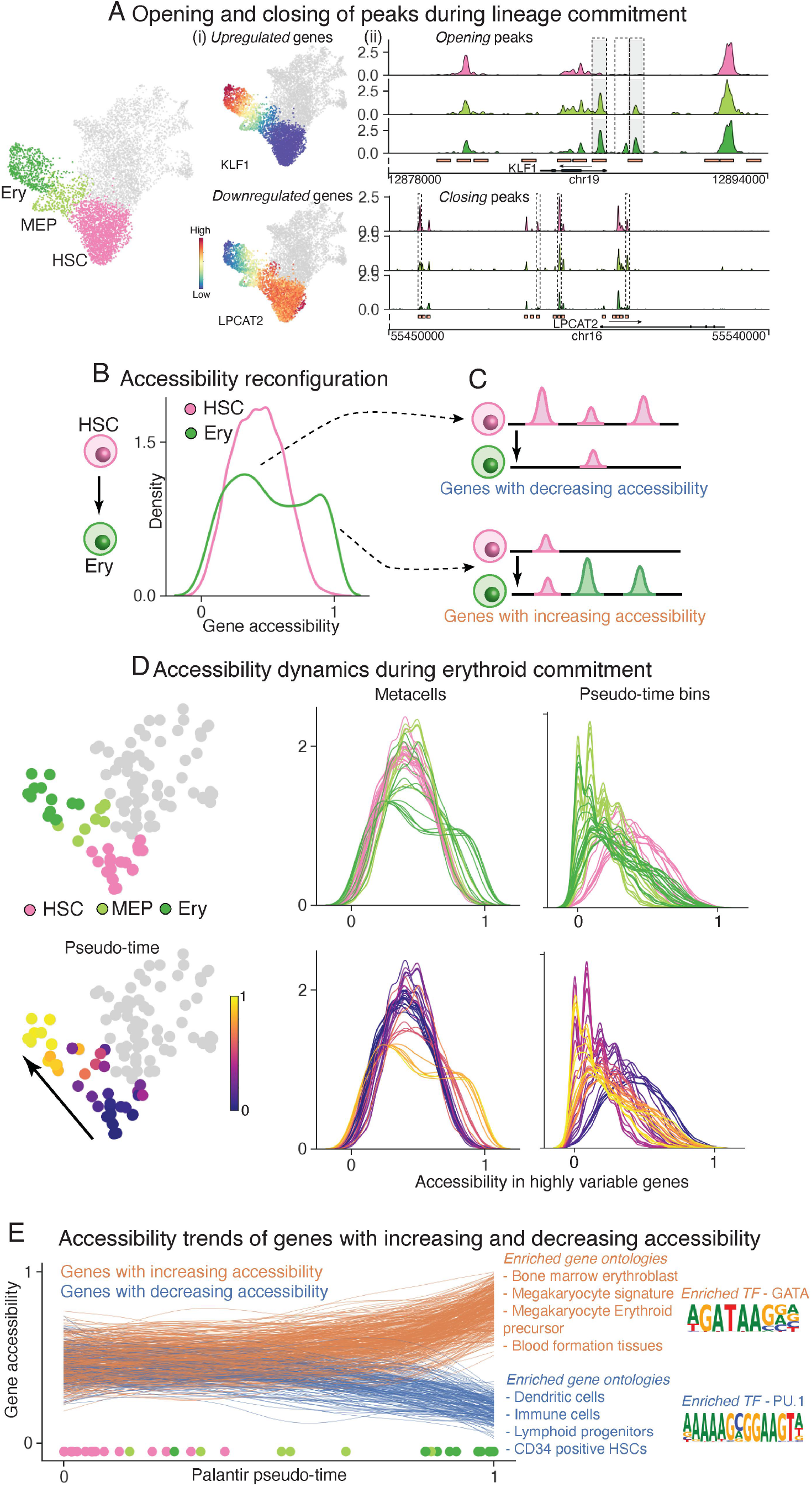
Charting chromatin accessibility of hematopoietic differentiation using SEACells metacells. A. Differentiation along a particular lineage involves upregulation of lineage-defining genes and downregulation of stem genes or genes that define other lineages. Left, RNA modality UMAP of CD34^+^ bone marrow, with erythroid lineage cells highlighted. Middle, UMAPs colored by expression of erythroid gene *KLF1* and stem gene *LPCAT2*, which are upregulated and downregulated, respectively, during erythroid differentiation. Right, accessibility landscapes of *KLF1* (top) and *LPCAT2* (bottom), aggregated by cell type, during erythroid differentiation. B. Distribution of gene accessibility for all highly regulated genes, for hematopoietic stem cells (HSC) and erythroid cells (Ery). Unimodal gene accessibility in HSCs is reconfigured to a bimodal distribution during erythroid differentiation. C. Cartoon representing observed peak dynamics: Bimodal distribution results from a subset of genes losing open peaks (top) and another subset gaining open peaks (bottom). D. Chromatin accessibility distribution of highly regulated genes in all metacells along the erythroid lineage (left, middle). Each line represents a meta-cell, colored by its stage (top) and pseudotime (bottom). The emergence of bimodality is gradual and continuous. Right, signal is poorly defined when using pseudotime bins rather than metacells. E. Accessibility dynamics of genes that gain (orange) and lose (blue) open peaks during differentiation from HSCs to erythroid cells. Trajectory computed using Palantir and each line represents a fit gene trend. Pseudotime of each respective meta-cell plotted on the bottom. Middle, results of gene ontology analysis using immune cell gene signatures. Opening peaks are enriched for GATA motifs, and closed peaks are enriched for PU.1, master regulators of erythroid and myeloid fates, respectively.

Tracking accessibility dynamics requires overcoming sparsity to identify which regulatory elements are open and accessible. We identified open elements in each metacell (**Supplementary Fig. 15**, Methods), then defined a metric of gene accessibility as the fraction of gene-associated peaks (**Fig. 3C**, Methods) that are open in a given metacell, ranging from 0 (all peaks closed) to 1 (all peaks open). Our accessibility scores track with gene expression for key lineage-specific genes, indicating that they accurately represent underlying biology (**Supplementary Fig. 16A**). For example, *GATA1* and *MPO* scores undergo specific increases in erythroid and myeloid lineages, respectively, consistent with their characterized roles^5^ (**Supplementary Fig. 16B**).

We next examined gene accessibility across cell compartments for all highly regulated genes, and observed that the earliest cell type, hematopoietic stem cell (HSC), follows a unimodal distribution centered at 0.5 (**Fig. 5B**). In contrast, expression of these genes in HSCs follows a long-tailed distribution, indicating that only a subset is expressed, and suggesting an epigenomic landscape in stem cells that is poised for hematopoietic gene expression (**Supplementary Fig. 17A**), as previously observed^18, 33, 34^. As cells differentiate along a lineage, genes that define the lineage gain accessibility peaks, while genes that define alternative lineages lose peaks (**Fig. 5B,C**). The resulting bimodality of differentiated cells is most clearly observed in the erythroid lineage (**Fig. 5B**). All other lineages show the emergence of long-tailed distributions (**Supplementary Fig. 17B**). Previous studies have shown that the erythroid lineage is established first^5, 35^, and posited that the lack of clear bimodality in other lineages could be due to the capture of CD34^+^ sorted cells that have not yet expressed lineage programs. We therefore performed a similar analysis on an scATAC-seq dataset of unsorted bone marrow mononuclear cells^19^ and observed more pronounced bimodality across hematopoietic lineages (**Supplementary Fig. 17C, D**).

We focused on gene accessibility dynamics in the erythroid lineage. We first applied Palantir^5^ to SEACells metacells using the RNA modality of multiome data to determine a pseudotime ordering of metacells along this lineage (**Fig. 5D**, Methods). We then examined the gene accessibility dynamics of highly regulated genes in each metacell along the pseudotemporal order and observed that epigenomic reconfiguration is itself gradual and continuous (**Fig. 5D**). Using pseudotime bins instead of metacells does not reveal any bimodality or dynamics, demonstrating that the resolution of SEACells metacells is uniquely suited for capturing dynamics (**Fig. 5D**). We next fit a generalized additive model to examine gene accessibility as a function of pseudotime (**Fig. 5E**, Methods). The absence of step-like behavior in any accessibility trends reinforces the continuous nature of epigenomic reconfiguration. Moreover, the opening and closing of regulatory elements at diverging lineage-specific loci mirrors each other (**Fig. 5E**), suggesting that similar mechanisms drive these two processes through gradual changes in plasticity and developmental potential, as observed in studies that combine lineage tracing with scRNA-seq profiling^36^.

Gene ontology enrichment analysis revealed that genes with increasing accessibility in the erythroid lineage establish erythroid cell identity and function, whereas those with decreasing accessibility are enriched for HSC and diverse other lineage identity genes, in further support of epigenomic priming in HSCs (**Fig. 5E**, Methods). Finally, the enrichment of TF motifs in peaks gained and lost in erythroid differentiation predicts a role for *GATA2* and *PU.1*, respectively (**Fig. 5E**, Methods), consistent with the known mutual antagonism of these factors in the decision between erythroid and myeloid lineages^37^. Together, our results show that SEACells metacells make it possible to model the dynamics of gene accessibility during differentiation. We find that a unimodal landscape of open chromatin in HSCs is reconfigured to a bimodal distribution in differentiated cells that involves the gradual and continuous opening and closing of peaks.

### SEACells enables the integration of large-scale single-cell datasets

Advances in scRNA-seq technology and atlas projects such as the Human Cell Atlas^1^ are prompting the generation of single-cell datasets spanning millions of cells^38–43^ and hundreds of individuals, rendering even the most fundamental analyses such as dimensionality reduction and clustering computationally infeasible. SEACells identifies robust, well-defined metacells from any sample, and thus can be used to integrate large-scale single-cell datasets in a computationally efficient manner. Moreover, by enumerating meta-cells on each sample, we provide per sample summary statistics that is less susceptible to batch effects, facilitating data integration that is better able to resolve biological (rather than technical) differences between individuals. We demonstrated the utility of SEACells using a recently published dataset of PBMCs containing over 175,000 cells from 23 healthy donors and 17 critical COVID-19 patients^44^.

We first applied SEACells to identify metacells in each sample (**Fig. 6A** and **Supplementary Fig. 18**) and verified that the metacell states are consistent across healthy donors and across COVID-19 patients (**Supplementary Fig. 19**, Methods). Encouraged by this high level of reproducibility, we used metacell gene expression counts for downstream tasks such as data integration^45^, clustering^46^ and visualization using UMAPs (**Fig. 6B**). Batch effects prior to integration are severe (**Supplementary Fig. 20A,B**), but are significantly lower in metacells compared to single cells (**Supplementary Fig. 20C**), enabling reasonable downstream analysis even without integration. While data integration eliminated batch effects in both single cells and metacells (**Supplementary Fig. 20C**), metacells required orders of magnitude less compute time compared to data integration, visualization, and clustering at the single-cell level (**Supplementary Fig. 21**). This ability to scale is particularly important when new data accumulates, and existing analyses need to be rerun. A single upfront investment in metacell assignment avoids compounding the near-exponential increases in runtime associated with adding cells, for each single-cell-level analysis (**Supplementary Fig. 21**).

**Fig 6:**
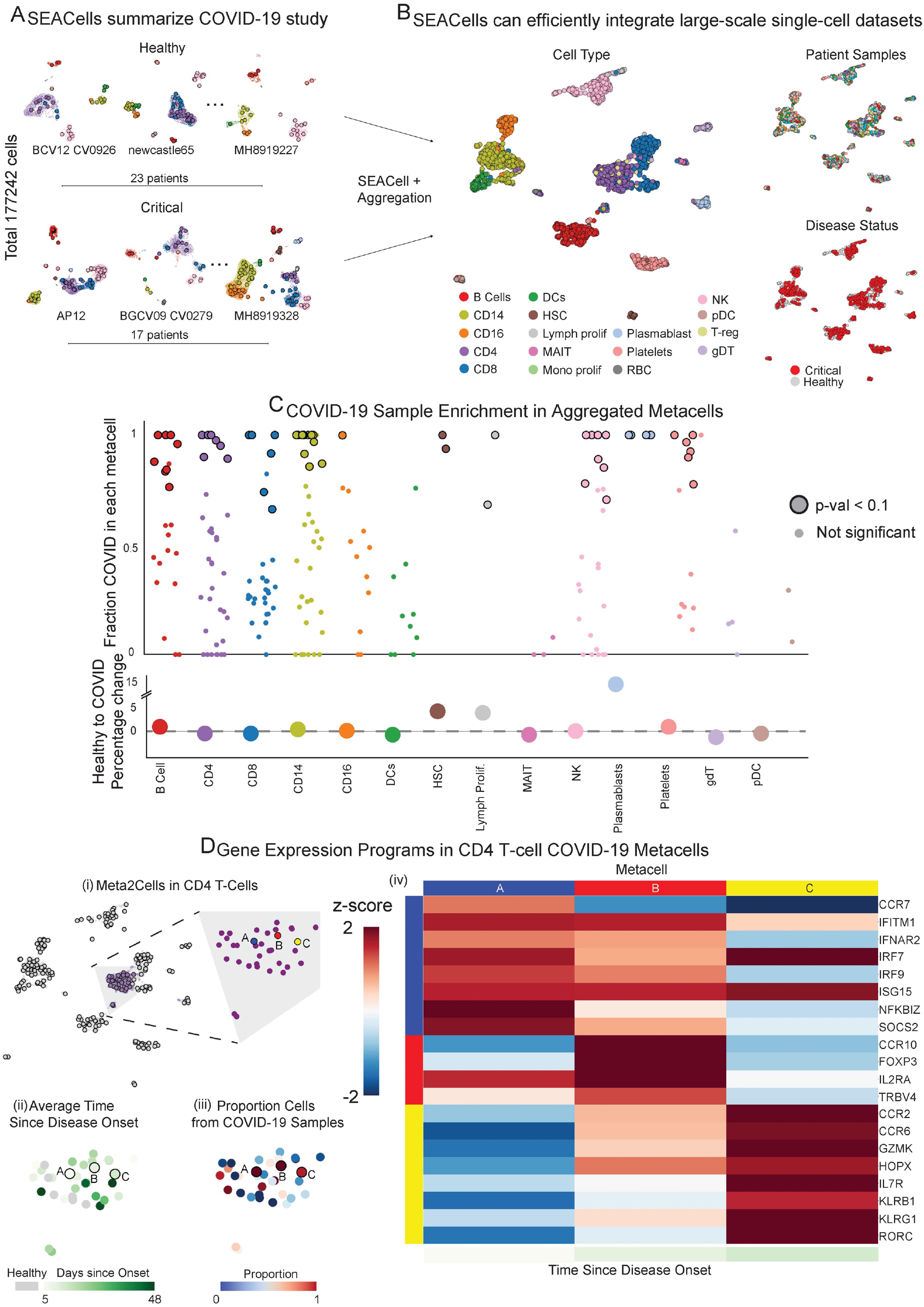
SEACells metacells identify dysregulated states in COVID-19 patients. A. UMAPs showing PBMC profiles and respective metacells for a subset of healthy patients and critical COVID-19 patients^44^. Dots, cells; circles, metacells. Cells and metacells are colored by cell type. B. UMAPs showing metacells from different patients integrated using Harmony^45^. Metacells are colored by cell type (left), sample (top right) or disease status (bottom right). C. Top: Differential abundance of SEACells metacell states in COVID-19 patients compared to healthy individuals, computed using a permutation test (Methods). Significantly differential metacells are plotted as enlarged circles. Bottom: difference in proportion of cells derived from COVID-19 patients compared to healthy individuals, analyzed at the cell-type level. D. (i) UMAP of metacell aggregates (meta2cells). Right: zoom-in on CD4 T-cell metacells. Three metacells enriched in COVID-19 patients compared to healthy donors are highlighted. (ii) Same as (i), with metacells colored by time since disease onset. (iii) Same as (i), with metacells colored by proportion of COVID-19 cells. (iv) Expression patterns of T-cell activation and differentiation enriched in highlighted CD4 T-cell metacells.

### SEACells identifies the temporal dynamics of T-cell response during COVID-19 infection

We next examined whether SEACells metacells can be used to identify state changes from healthy to severe COVID-19 patients. We pooled metacells from all donors and re-applied SEACells to derive metacell aggregates representing states across all samples (**Supplementary Fig. 22A**). Each aggregated metacell is a combination of healthy and COVID-19 metacells, such that the fraction of COVID-19 cells can be visualized for each state (**Supplementary Fig. 22B**). Our results reveal a broad spectrum of metacell states, from those specific to healthy donors to those exclusive to COVID-19 (**Supplementary Fig. 22B**), prompting us to develop a permutation test to identify cell-states that differ significantly between the two (**Fig. 6C**, Methods). By contrast, analysis at the cell-type level completely masks the extensive heterogeneity present in individual states within the cell-type (**Fig. 6C**).

We focused our analysis on CD4^+^ T cells, which are known to differentiate into distinct subsets upon activation and differentiation^25, 26^, using differential gene expression analysis at the metacell level to identify cell-state defining genes. Within CD4^+^ T-cell metacells, this analysis revealed a fine-grained trajectory of phenotypes enriched in patients with COVID-19, with meaningful correspondence between T cell phenotypes and temporal stage of disease (**Fig. 6D**). For example, a metacell enriched in COVID-19 patients soon after infection (metacell A) contains cells in an early activation state distinguished by the expression of NF-*κ*B response genes, IFN-*α* receptor subunit *IFNAR2,* and downstream interferon-stimulated genes (*IRF7, IRF9, ISG15, IFITM1*), reflecting T cell responsiveness to type I IFN, a cytokine associated with viral infections and SARS-COV2 pathology^47^ (**Fig. 6D**). A metacell enriched in COVID-19 patients approximately 10 days after symptom onset (metacell B) comprises Foxp3^+^ Treg cells expressing the chemokine receptor gene *CCR10*, suggesting recruitment to the inflamed lung or mucosal epithelium and a potential role in regulation of inflammation^48^ (**Fig. 6D**). Finally, a metacell enriched in patients with persistent severe COVID-19 at day 13 (metacell C) contains cells that express hallmark T_H_17 genes (*RORC* and *CCR6*), reflecting a shift towards type III inflammation.

We postulate that while data integration methods aim to make samples more similar without distinguishing between batch and biological signal, aggregating data into metacells on the per-sample level provides robust capture of true biological variation between the samples. Indeed, our results show that SEACells can capture biologically meaningful CD4 T-cell subsets and states, highlighting the full spectrum of activation and differentiation during an evolving viral infection, in which cells transition between active and quiescent states over the course of hours and days. Differential abundance testing^15^ at the single-cell level failed to recover these dynamics (**Supplementary Fig. 22C,D**).

## Discussion

SEACells identifies robust, reproducible metacells from single-cell data that overcome sparsity while retaining the rich heterogeneity of the data. SEACells metacells are more compact, better separated, and more evenly distributed across the full cell-state landscape than metacells generated by existing methods. We have shown that they faithfully represent both discrete and continuously varying cell states, and that they provide enormous benefits for scaling to large cohort-based datasets, including carrying out data integration across samples and modalities. Critically, only SEACells is currently able to derive cell states from scATAC-seq data in an accurate and comprehensive manner, greatly empowering gene regulatory inference.

The performance of SEACells is due to its (i) representation of single-cell phenotypes using an adaptive Gaussian kernel to accurately capture the major sources of variation in the data, (ii) use of max-min cell sampling for initialization to ensure even representation of cell-states across phenotypic space, regardless of cell densities, and (iii) application of archetypal analysis for identifying highly interpretable metacells. The adaptive kernel and max-min sampling make SEACells particularly adept at robustly identifying rare cell-states, which often represent critical populations that drive biology or disease.

Kernel representation also eliminates the need for specific data representations such as gene scores, allowing SEACells to generalize to multiple modalities. We show that metacells are particularly effective on scATAC-seq data, which is currently analyzed at the cluster level due to extreme sparsity, and thus remains underutilized. Whereas gene scores, open regulatory elements, and correlations between gene expression and chromatin accessibility cannot be determined robustly at the single-cell level, they can be computed for individual metacells. Such improvements in fundamental ATAC analysis enable more sophisticated regulatory network inference, promising wide utility for SEACells in studies with single-cell chromatin profiling data.

Our procedures for computing peak-to-gene associations, gene scores and gene accessibility assume the availability of either multimodal data or integrated RNA and ATAC modalities. Several approaches have been developed for data integration across modalities^13, 49^ and are likely to exhibit improved performance when applied at the metacell level. Given the kernel representation, SEACells should generalize to other modalities such as CUT&Tag^50, 51^ or other single-cell chromatin modification measurements^52^ with appropriate preprocessing.

We also anticipate that SEACells will be used extensively as a scalable solution for integrating large single-cell datasets from cohorts. Metacells can be computed separately for each sample, rendering the integration of additional cohort members extremely resource-efficient, while retaining heterogeneity in the data. Computing metacells at the sample level also provides a more robust representation of sample-specific biology than data integration approaches that struggle to distinguish biological and technical differences between samples. Aggregating dozens of cells per metacell provides a more comprehensive expression or chromatin accessibility profile, and also generates a distribution over these features, facilitating comparison between metacells across samples. As demonstrated in COVID-19 data, sample-level sufficient statistics provided by SEACells are particularly well suited to compare disease states between healthy and normal, as well as more nuanced disease states such as progression.

An important consideration when running SEACells is how to specify the number of metacells. There should be enough metacells to capture cell states at high resolution, while maintaining sufficient cells per metacell to ensure robustness. The optimal number thus depends on biological heterogeneity in the dataset and the total number of cells profiled. For example, cells from a homogeneous cell line will have less biological structure compared to a similar sized tissue sample. To choose the number of metacells, we recommend examining the initialization to ensure that cell-states span the full phenotypic manifold (Methods).

Metacell analysis allowed us to determine a metric for gene accessibility and to demonstrate that chromatin landscape reconfiguration is continuous and gradual during hematopoietic differentiation. Further, we utilized the scalability and robustness of SEACells to integrate a large-scale COVID-19 scRNA-seq dataset and identify a disease progression of COVID-19-enriched CD4^+^ T-cell states relating to differentiation and activation. These critical states are not detected by differential abundance testing at the single-cell level. In addition to enabling cohort-scale analysis, SEACells metacells serve as more robust cell-state inputs, which facilitates the distinction of biological signal from batch effect—features that enabled our discovery of the T-cell state continuum. SEACells is a powerful discovery tool for emerging single-cell cohorts.

## Data Availability

The newly generated CD34+ bone marrow and T-cell depleted bone marrow multiome datasets will be deposited to GEO. Filtered and processed count matrices including cell-type annotations and ATAC fragment files are available on Zenodo at **`10.5281/zenodo.6383269`**.

## Code Availability

SEACells is available as a Python module at https://github.com/dpeerlab/SEACells. Jupyter notebooks detailing the usage of SEACells include metacell identification, aggregation and the ATAC preprocessing, and gene regulatory toolkit are available at https://github.com/dpeerlab/SEACells/tree/main/notebooks. Modified ArchR pipeline for peak calling using NFR fragments is available at https://github.com/dpeerlabArchR

## Author Contributions

M.S and D.P conceived and designed the study. S.P, Z-N.C, M.S and D.P developed the SEACells algorithm. S.P, I.P, M.S and D.P developed additional analysis methods, statistical tests and analyzed the data. S.P and M.S implemented SEACells and other analysis methods. I.M and R.C designed, optimized, and executed all single-cell multiome experiments. C.D, M.S and D.P performed analysis of hematopoietic dynamics. S.P, C.C, M.S and D.P performed COVID-19 analysis. S.P, T.N, I.P, M.S and D.P wrote the manuscript.

## Acknowledgements

We thank Cassandra Burdziak, Joe Chan and Ricard Argelaguet for valuable conversations related to this manuscript. This study was supported by NCI grant U54 CA209975 and NCI Human Tumor Atlas Network U2C CA233284 (D.P), Alan and Sandra Gerry Metastasis and Tumor Ecosystems Center at MSKCC (I,M and R.C) and Functional Genomics Initiative at MSKCC (D.P).

## Methods

### SEACells algorithm

SEACells - Single-cEll Aggregation of High-Resolution Cell-states - is an algorithm to determine metacells, groups of cells that represent singular cell-states from single-cell data. The SEACells algorithm assumes that biological systems consist of well-defined and finite sets of cell-states defined by co-varying patterns of gene expression. Observed single-cell data are assumed to be noisy measurements of these cell-states with current state of the art single-cell measurement technologies able to capture <10% of transcripts or <5% open chromatin regions. Despite the high degree of noise, cells sampled from the same states are assumed to be closely related in their phenotypes as a result of gene expression patterns and regulatory mechanisms that define the cell-states. Thus, SEACells algorithm aims to aggregate closely related single-cells and identify metacells that represent cell-states. As a result of aggregation, metacells overcome the sparsity issues that plague single-cell data, with single-cell ATAC-seq data particularly limited in its utility due to its sparsity. SEACells metacells also provide a scalable representation to efficiently handle large-scale single-cell data. While clustering is widely used to overcome issues of sparsity, clustering masks the substantial heterogeneity present in the data (**Fig. 1A-D**). SEACells metacells achieve a resolution that retains the heterogeneity while overcoming the sparsity issues of single-cell data.

The major inputs to SEACells algorithm are: (i) raw count matrices (E.g.: gene expression for RNA, peak or bin counts for ATAC), (ii) low dimensional representation of the data such as PCA derived using an appropriate preprocessing procedure dependent on data modality and biological system and (iii) the number of metacells to be identified. Using this information, SEACells produces as output groupings of cells that represent metacells and aggregated metacells X feature raw counts matrices. SEACells algorithm is available as a Github repository as https://github.com/dpeerlab/SEACells. In addition to documentation and tutorials for computing metacells, the repository also includes tutorials for computation of gene expression - peak accessibility correlations, ATAC gene scores, open peaks in metacells and gene accessibility scores using multiome or integrated RNA & ATAC data.

SEACells comprises five main steps:

1. Construct a k-nearest neighbors graph using Euclidean distances between cells computed in the lower dimensional embedded space. This KNN graph provides a representation of the phenotypic manifold to identify tightly connected groups of cells, to be aggregated into metacells.
2. An affinity matrix of cell-to-cell similarities is derived using the nearest neighbor graph. The distances in the graph are transformed to similarities using an adaptive Gaussian kernel to account for the dramatic differences in densities in the phenotypic manifold spanned by single-cell data. The affinity or kernel matrix (**Fig. 1F**) also encodes the non-linear relationships between cells .
3. The kernel matrix then serves as input to kernel archetypal analysis, for linear decomposition of the input single-cell data. The linear nature of this procedure maximizes interpretability. Archetypal analysis decomposes the data into an archetype matrix, linear combinations of cells that are representative of the cell-states on the phenotypic manifold and a membership matrix that reconstructs the single-cells as linear combinations of archetypes (**Fig. 1G**). This procedure partitions the data in such a way that the cell-cell similarity matrix has tight block structure along the diagonal, which best represents distinct cell states in the data (**Fig. 1H**). Each distinct partition is a group of cells and represents a metacell. The specified number of metacells are used as input to archetypal analysis.
4. The groupings identified through archetypal analysis are SEACells metacells. Single-cell raw counts are aggregated using these groupings to derive a metacell X feature count matrix.
5. Normalized metacell count matrices can be used for all downstream tasks including clustering, visualization, data integration, trajectory inference, ATAC-seq based regulatory inference and other analysis performed with single-cell data.

These design principles ensure that metacells identified by SEACells algorithm are robust, compact, well-separated, provide sufficient meta-cells, and span the entire phenotypic manifold. SEACells can be applied to single-cell datasets with discrete cell-types and continuous trajectories. SEACells has been tested and benchmarked using scRNA-seq and scATAC-seq datasets and in principle can be applied to other single-cell modalities as well. An appropriate preprocessing procedure that generates a faithful low-dimensional representation of the single-cell data is a critical element for success of SEACells algorithm to generalize to additional data modalities. We have outlined a procedure to analyze each data modality separately, but a graph or representation derived using multiple modalities^13, 49^ can also be used as input to SEACells.

#### Low Dimensional Embedding

A central input to SEACells algorithm is a low dimensional representation of single-cell data. This representation is used as input to construct the k-nearest neighbor graph using Euclidean distance between cells. Single-cell data is extremely noisy due to low capture rates, and as a result measuring distances between cells in the measured expression or accessibility is unreliable. A low dimensional representation such as PCA, overcomes the noise in the data with the top components encoding information about biological variability. Low-dimensional embedding can be derived by using appropriate pre-processing and normalization steps for the data modality of interest (**Supplementary Fig. 2**). This allows us to be both flexible to data type, and robust to the extensive degree of sparsity and noise in data types such as scRNA-seq and scATAC-seq. We utilized the following preprocessing steps adapted to the characteristics of each technology.

#### PCA for scRNA-seq

Following standard practices, we perform three main pre-processing steps using the scanpy^53^ package: 1) library size normalization by dividing raw counts by total molecules per cell, 2) log-transformation with a pseudo-count of 0.1 and 3) selection of highly variable genes. Based on our prior observations for PBMCs and CD34 bone marrow datasets , we chose 2500 highly variable genes for analysis. This number should be adapted to ensure all the heterogeneity is captured in the dataset of interest. Principal components are computed from these highly variable genes, with the number of principal components being selected based on proportion of variance explained (Typically 50).

#### SVD for scATAC-seq

We used the ArchR package^23^ for preprocessing of sc-ATAC data. Fragment counts for each cell were computed in 500 base genome bins. The counts were normalized using TF-IDF^54^. The normalized counts were used as input to derive a low-dimensional embedding using SVD. Similar to PCA, the number of principal components were selected based on the proportion of variance explained (Typically 30). As has been observed previously, despite normalization the first SVD component shows high correlation with number of fragments per cell (correlation > 0.97) and is excluded from downstream analysis.

#### Nearest Neighbor graph

A k-nearest neighbor graph is constructed using Euclidean distance in the low-dimensional embedding (PCA or SVD). The graph is a representation of the phenotypic manifold, with nodes as single cells, each connected to their most similar neighbors. The nearest graph can be represented as a matrix *D* ∈ *R^nXn^*, where *n* is the number of cells. *D_ij_* represents distance between cells *I* and *j* if they are neighbors and *D*∈*R^nXn^*, otherwise. The graph serves as input for the construction of the cell-cell kernel matrix. As default, 50 neighbors are used for nearest neighbor graph construction, and we have previously demonstrated that the kernel matrix construction is robust to a reasonable range of number of nearest neighbors^5^.

#### Adaptive Gaussian kernel

The goal of SEACells algorithm is to identify metacells that are tightly related groups of cells to represent cell-states (**Fig. 1H,G**). Therefore, we need to transform the *distances* in the neighbor graph to *similarities* between neighboring cells. Gaussian kernels provide an approach for this transformation but assume that densities in underlying data are approximately uniform. Single-cell data, however, show remarkable variability in data densities (**Supp. Fig. 6**) with low-density regions or rare cell-types often the most meaningful in describing the biology of the system. We have previously demonstrated that an adaptive kernel that uses neighbor distance as the scaling factor for each cell, rather than a fixed parameter, is highly effective in faithfully representing the phenotypic similarities^5, 55^. Therefore, SEACells uses an adaptive (width) Gaussian Kernel to determine similarities between cells and more faithfully represent the underlying phenotypic manifold (**Fig. 1F**). The adaptive kernel corrects for densities using the distance to the *l^th^* (*l*<*k*) nearest neighbor as a scaling factor i.e, the scaling factor of cell *i* is given by σ = distance to *l^th^* nearest neighbor.

The adaptive Gaussian kernel is then given by

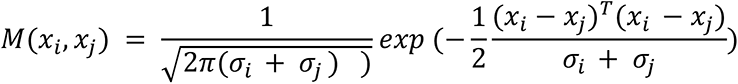

where *x_i_* is the low-dimensional embedding corresponding to cell *i*.

*M*∈*R^nXn^* is the affinity matrix. *M_ij_* represents the similarity between cells *I* and *j* if they are neighbors and *M_ij_* = 0 otherwise.

#### Kernel Archetypal Analysis

The adaptive kernel serves as input to a kernel archetypal analysis procedure . Archetypal analysis identifies a linear decomposition of the data matrix where the goal is to identify a specified number of archetypes that are each a linear combination of the data points represented by the archetype matrix. The data points themselves are represented as a linear combination of the archetypes in a membership matrix to reconstruct the original data matrix (**Fig. 1G**). Therefore, archetypal analysis functions in a manner similar to an autoencoder with the lower dimensionality of the archetype and membership matrices creating an information bottleneck that ensures an optimal linear decomposition of the data^17^. Membership matrices can be used to derive cell partitions that are aggregated to metacells (**Fig. 1H**). The linear nature of archetypal analysis ensures maximal interpretability and identification of metacells. Further, the iterative procedure of archetypal analysis supports the use of cell-cell similarity kernels which already encode the non-linear relationships between cells.

Formally, the data matrix *X*∈*R^d×n^* consists of *d*-dimensional vectors corresponding to *n* cells. Here *d* is the dimensionality of low-dimensional embedding such as PCA or SVD. The goal of archetypal analysis is to decompose data as *X*≈*ZA* i.e, the data matrix *X* is represented as a convex combination of a latent archetype matrix *Z*∈*R^d×s^* and cell membership matrix *A*∈*R^s×n^*, where *s*<<*n* is the number of archetypes. As these latent archetypes are *a priori* unknown, they are themselves defined as convex combinations *Z* = *XB* of the data, *X*, and archetype weight matrix *B*∈*R^n×s^*. To ensure that data points are convex combinations of archetypes, and vice versa, weight matrices *A* and *B* must be row- and column-stochastic, respectively, such that their entries are non-zero and rows/columns sum to one.

Formally, for entries *a_ji_* ∈ *A* and *b_ji_* ∈ *B*,

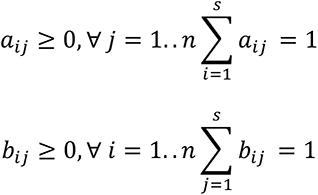

Taken together, the objective of archetypal analysis is to find matrices *A*, *B* such that product *XBA* forms a faithful reconstruction of the original data matrix *X*.

The objective of kernel archetype analysis is to minimize squared reconstruction error (SRE) as follows:

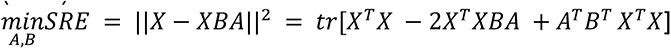

Note that this optimization problem depends only on the dot product i.e., linear kernel of the data matrix *X*. Therefore, non-linear kernels can be substituted to model systems where the data is generated from non-linear combinations of archetypes using the kernel trick. We therefore replace the linear kernel with the adaptive gaussian kernel *M* and the optimization problem can then be formulated as.

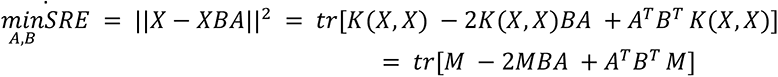

The number of archetypes, *s*, representing the number of metacells, is a parameter.

#### Optimizing Archetypes and Cell Assignments

The objective function for kernel archetype analysis involves optimizing the non-convex product *AB*, and thus has many local minima. The objective function is, however, convex in *A* given a fixed *B* matrix, and vice versa. Therefore, alternating minimization of weight matrices *A* and *B* is used to make the problem of solving archetypal analysis more tractable. Given this, we use the Frank-Wolfe updates to optimize each weight matrix in turn, as described in ^17^.

#### Initialization

As archetypal analysis is a non-convex problem, solutions depend on the initialization of archetype and cell assignments. Given the density differences in the phenotypic manifold, random sampling of cells will lead to significant overrepresentation of initial points in the high density regions and severe underrepresentation of cells in the biologically critical low density regions. Therefore, we employ max-min sampling of waypoints, described and implemented by ^5^ to initialize archetypal analysis.

Given a set of waypoints, each additional waypoint is chosen to maximize the distance to the current set, i.e. maximize the minimum distance to any of the points in the current set. This ensures that the sampled waypoints are uniformly distributed across the phenotypic manifold irrespective of the density (**Fig. 1E**). We first derive a diffusion map embedding using the adaptive Gaussian kernel *M*. As demonstrated previously^5^, each diffusion component represents an axis of biological variance in the data. Diffusion maps can be used to describe both continuous trajectories as well as to define separation between discrete cell-types. Waypoints are sampled from each component and pooled for initialization. The number of components can be chosen by the eigengap statistic, though in practice we observed that the first 10 diffusion components typically account for all the biological variability in the data.

Waypoint sampling is used to initialize the matrix *B*, following which matrix *A* is updated and the process is repeated until convergence.

#### Metacell identification

Archetypal analysis computes partial assignments of cells to archetypes. However, in order to aid in interpretability and facilitate downstream analysis, metacells are constructed by (1) computing binarized assignments of cells to archetypes (of the *A* matrix) and (2) aggregating single cells assigned to each SEACell by summing over raw counts (**Fig. 1G**). This summarized SEACells data matrix is significantly less sparse and noisy and can then be used for more robust downstream analysis.

#### Metacell normalization

Metacell raw counts can be normalized analogous to single-cell data normalization. Metacell counts are divided by the total counts per metacell and then multiplied by the median of the total metacell counts to avoid numerical issues. The data is then log-transformed using a pseudo count of 0.1

#### Notes about number of metacells

The number of metacells should be specified as a parameter to the SEACells algorithm. We have evaluated the robustness of SEACells to the number of metacells and determined that the results are robust across a wide range of parameters (**Supplementary Fig. 5**). We currently use a heuristic of one metacell per 75 single-cells in the dataset under consideration. However, the number is largely dependent on the biological structure in the data. As an example, a dataset profiling 10k cells from a homogeneous cell-line will be expected to be less heterogeneous and thus encode less biological structure compared to a similar sized single-cell dataset of more complex biological systems such as tumors or differentiation. Therefore, we recommend examination of initialization to ensure that cell-states span the entirety of the phenotypic manifold to choose the number of metacells.

### Toolkit for scATAC analysis

Analysis of bulk ATAC-seq data has provided a broad array of tools that are powerful in interpretation of open chromatin data. Direct application of these tools is prohibitive for single-cell data analysis due to issues of sparsity. SEACells metacells are aggregates of tightly related cells and therefore are substantially less sparse and encode more information per cell-state compared to analysis at the single-cell level. Metacells faithfully retain the heterogeneity and structure of the data, and hence enable the construction of a robust toolkit for scATAC-seq data analysis with adaptations of tools from bulk data analysis.

#### Peak calling

Peak calling was performed using ArchR^23^. ArchR first clusters single-cell data and uses the MACS2 peak caller^56^ to identify peaks separately for each cluster. Each peak is then resized to 500 bases with the peak summit at the center and overlapping peaks across different clusters are merged. The merged peaks are again resized to 500 bases.

ATAC-seq data provides a profile of open chromatin regions spanning transcription factor binding and nucleosomes in non-repressed regions. The fragment size distribution of ATAC-seq data contains characteristic modes which reflect the diversity of information (**Supplementary Fig. 7A**). Since the first mode represents nucleosome free regions (NFR), we implemented a change in the ArchR pipeline to identify peaks using only the NFR fragments (fragment length < 147) rather than use of all fragments as is default in ArchR. This change leads to substantially better sensitivity in identification of regulatory elements (**Supplementary Fig. 7B,C**).

The modified ArchR pipeline is available at https://github.com/dpeerlab/ArchR

#### Peak-gene associations and gene scores

While use of NFR fragments improves the sensitivity of called peaks, not all identified peaks represent TF binding events that directly regulate the expression of a gene. Examples include structural factors such as CTCF which also show a signal in ATAC-seq data but do not directly regulate the expression of a gene. Therefore, studies have proposed the use of correlation of peak accessibility and gene expression using multiome or integrated ATAC & RNA data to identify the candidate list of peaks i.e., regulatory elements that likely regulate the expression of the gene^18^. SEACells metacells retain the heterogeneity in data but overcome data sparsity and thus provide an ideal resolution to compute these associations which are unreliable when computed using single-cell data due to high degree of noise and sparsity. We use metacells identified using the ATAC modality for building the peak-gene associations t for robust associations.

We adopted the procedure outlined by Ma et. al.^18^ to identify significant peak-gene associations. For each gene, Pearson correlations were computed for each peak across a span of 100kb upstream and 100kb downstream of the gene using the normalized metacell expression and normalized ATAC accessibility. To assess the significance of the peak-gene correlation, an empirical background of 100 peaks were sampled that matched the GC content and accessibility of the peak under consideration. Peaks were binned into 100 bins separately based on GC content and accessibility to sample the empirical background. Any peak with a nominal p-value < 1e-1 was considered a significant peak-gene association. Peaks identified using NFR fragments were used for this analysis. The aggregate accessibility of all peaks associated with a gene was used to determine the metacell gene score.

For single-cell comparisons, normalized single-cell expression and normalized single-cell accessibility were used for determining peak-gene associations. Gene scores for single-cell ATAC were computed using the ArchR^23^ defaults.

#### chromVAR using SEACell metacells and single-cells

chromVAR^24^ was run using default parameters using the chromVAR “human_pwms_v2” motif database. chromVAR scores were computed using aggregated fragment counts for metacells and single cell fragment counts for single-cell data. Similar to the single-cell data analysis, chromVAR scores were first reduced to 50 principal components using knee-point analysis. PCs then served as input to umaps for visualization.

#### Metacell peak calling

Identification of the set of open regulatory elements is practically implausible at single-cell level due to noise and sparsity. SEACells metacells however, provide a sufficient number of fragments per cell-state to enable the identification of open regulatory elements in each state. We observed that *de novo* peak calling in each metacell results in loss of sensitivity (**Supplementary Fig. 7**).. Therefore, we use the peaks identified by ArchR across all cells as an atlas to determine the subset of peaks open in each metacell.

A procedure inspired by MACS2 is used to identify open regulatory elements in metacells since the peaks themselves were called by MACS2. The fragments mapping to peaks are modeled as a Poisson distribution. The mean of the Poisson distribution for a metacell *s* is estimated using^56^:

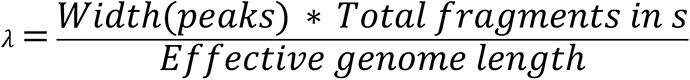

Since all the widths are identical, the first term of the numerator is set to 500. Rather than use the whole genome length as the denominator, *effective genome length* was set to be *num. of peaks * 5000*, a more stringent local estimate of the mean as proposed in MACS2. For a peak *p* in metacell *s* with *n* fragments , λ is used to estimate the *p-value* of observing more than *n* fragments and *p* is considered open in *s* if *p-value < 1e-2*.

We noticed that some of the ATAC metacells had low overall fragment counts - therefore we computed fragments per peak and total fragments from 2 nearest metacells. We apply this procedure for all metacells to avoid any biases.

#### Gene accessibility scores

Gene accessibility scores for a gene and metacell is defined as the fraction of gene associated peaks that are open in the particular(**Fig. 4B**). Gene accessibility scores range from 0, indicating all correlated peaks are closed, to 1, indicating that all correlated peaks are open.

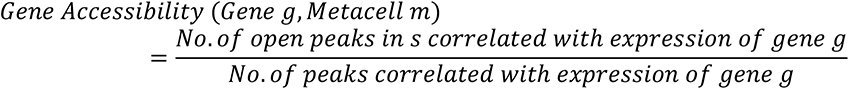

### Multiome data generation

#### CD34+ bone marrow cells

Cryopreserved bone marrow stem/progenitor CD34+ cells from a healthy donor were purchased from AllCells, LLC. (catalog no. ABM022F) and stored in vapor phase nitrogen. Vial was removed from the storage and immediately thawed at 37 °C in a water bath for 2 min while gently shaking. Next, vial content (1 mL) was transferred to a 50-mL conical tube. To prevent osmotic lysis and ensure gradual loss of cryoprotectant, 1 mL of warm medium (IMDM with 10% FBS supplement) was added dropwise after washing the storage vial, while gently shaking the tube. Then, the cell suspension was serially diluted 5 times with 1:1 volume additions of warm complete growth medium with 2-min wait between additions. The final ∼32-mL volume of cell suspension was pelleted at 300g for 5 min. After removing supernatant, cells were washed once with 10-mL of warm media and twice in ice-cold 1× PBS with 0.04% (wt/vol) BSA supplement to remove traces of medium. Cell concentration and viability were determined with a Countess II automatic cell counter using 0.4% trypan blue staining method.

Single Cell Multiome ATAC + Gene Expression was performed with a 10X genomics system using Chromium Next GEM Single Cell Multiome Reagent Kit A (catalog no. 1000282) and ATAC Kit A (catalog no. 1000280) following Chromium Next GEM Single Cell Multiome ATAC + Gene Expression Reagent Kits User Guide and demonstrated protocol - Nuclei Isolation for Single Cell Multiome ATAC + Gene Expression Sequencing. Briefly, 200,000 cells (viability 95%) were lysed for 4min and resuspended in Diluted Nuclei Buffer (10x Genomics, PN-2000207). Lysis efficiency and nuclei concentration was evaluated on Countess II automatic cell counter by trypan blue staining. 9,660 nuclei were loaded per transposition reaction, targeting recovery of 6,000 nuclei after encapsulation. After transposition reaction nuclei were encapsulated and barcoded. Next-generation sequencing libraries were constructed following the User Guide, which were sequenced on an Illumina NovaSeq 6000 system.

#### T-cell depleted bone marrow cells

Cryopreserved bone marrow cells from healthy donor were purchased from AllCells, LLC. (ABM007F) and stored in vapor phase nitrogen. Vial was removed from the storage and immediately thawed at 37 °C in a water bath for 2 min while gently shaking. Next, vial content (1 mL) was transferred to a 15 mL conical tube. To prevent osmotic lysis and ensure gradual loss of cryoprotectant, 1 mL of warm medium (IMDM with 10% FBS supplement) was added dropwise after washing the storage vial, while gently shaking the tube. Then, the cell suspension was dropwise diluted to 15 mL by addition of warm complete growth medium. The final 15 mL volume of cell suspension was pelleted at room temperature, 400*g* for 5 min. After removing supernatant, cells were washed once with 1 mL of Cell Staining Buffer (CSB), Biolegend (420201), cells were centrifuged again at 400*g* for 5 minutes 4 °C and resuspended in 100 uL of CSB. Concentration and viability were determined with a Countess II automated cell counter using 0.4% trypan blue staining method. Cells were incubated with Human TruStain FcX™ (Fc Receptor Blocking Solution), Biolegend (422301), for 10min at 4 °C. After blocking, bone marrow cells were stained with CD3 Monoclonal Antibody (UCHT1), PE-Cyanine7, eBioscience™ (25-0038-42) 1:100 for 20 minutes at 4 °C. Cells were washed 2 times with CSB before Fluorescence-activated cell sorting (FACSymphony S6, BD Biosciences) where CD3 negative cells were collected. Sorted cells were concentrated, count and viability was determined with a Countess II automated cell counter using trypan blue staining.

Single Cell Multiome ATAC + Gene Expression was performed with a 10X genomics system using Chromium Next GEM Single Cell Multiome Reagent Kit A (1000282) and ATAC Kit A (1000280) following Chromium Next GEM Single Cell Multiome ATAC + Gene Expression Reagent Kits User Guide and demonstrated protocol - Nuclei Isolation for Single Cell Multiome ATAC + Gene Expression Sequencing. Briefly, 300,000 cells (viability 95%) were lysed for 4 min and resuspended in Diluted Nuclei Buffer, 10x Genomics (2000207). Lysis efficiency and nuclei concentration was evaluated on Countess II automated cell counter by trypan blue and DAPI staining. 16,100 nuclei were loaded per transposition reaction, targeting recovery of 10,000 nuclei after encapsulation. After transposition reaction nuclei were encapsulated and barcoded. Next-generation sequencing libraries were constructed following the 10X Genomics User Guide and were sequenced on an Illumina NovaSeq 6000 system.

### Application of SEACells to PBMCs and bone marrow datasets

#### Data preprocessing

##### CD34+ Bone marrow Multiome data - RNA modality

Count matrices for the two samples were generated using CellRanger ARC^57^. Starting with the filtered barcode matrices from CellRanger ARC, barcodes from the bottom and top 2.5th percentile in molecule counts were excluded. Further cells with less than 0.4 fraction of reads in peaks in ATAC modality and greater than 20% of reads from mitochondria from the RNA modality were excluded from downstream analysis. The specified cutoffs were also chosen based on the respective empirical distributions to remove outliers.

Data was generated in two lanes. For each sample, scrublet^58^ was used to compute doublet scores using default parameters and a cluster of cells with high doublet scores were removed from downstream analysis (note, CD34+ has a continuous nature that can lead to false doublet calls). Following the filtering steps, the two samples were concatenated, normalized for molecule counts by dividing the raw data by the total counts per cell. The normalized data was multiplied by the median of total counts across cells to avoid numerical issues and log transformed with a pseudo-count of 0.1. Feature selection was then performed to select the top 2500 most highly variable genes (using scanpy.pp.highly_variable_genes) which was then used as input for principal component analysis with 50 components. The parameters were chosen based on prior analysis on a CD34+ scRNA-seq dataset^5^.

The PCs were used as input for generating umaps and clustering using phenograph^14^. The preprocessing and analysis was undertaken using scanpy^53^. Diffusion components were generated using the adaptive kernel following the functions in the Palantir package^5^ and imputation of gene expression was performed using MAGIC^55^. Each cluster was annotated as specific cell-types using the markers defined in^5^, following which mature B-cells were excluded from the analysis. Highly variable gene selection, PCA, clustering, visualizations, diffusion maps and imputation were repeated following B-cell exclusion. A total of 6881 cells were retained after all the filtering steps.

##### CD34+ Bone marrow Multiome data - ATAC modality

Analysis of the ATAC modality was undertaken using the ArchR pipeline^23^ using the subset of cells post-filtering from the RNA modality. 100k features were used instead of the default 25k features for ArchR processing. Using ArchR, data was normalized using IterativeLSI and SVD used to determine a lower dimensional representation of the sparse data. The first SVD component showed a greater than 0.97 correlation with log library size and was excluded from downstream analysis. SVD was used as input to cluster the data using phenograph and visualize using umaps. Peak calling was performed using the modified ArchR pipeline described in “Peak calling”.

##### T-cell depleted Multiome data

The preprocessing and analysis of RNA and ATAC modalities were performed following the steps outlined for the CD34+ bone marrow data. NK cells, mature monocytes and B-cells were excluded from the analysis in **Supplementary Fig. 6**. A total of 7439 cells were retained after all the filtering steps.

##### PBMC Multiome data

Counts for the PBMC Multiome data were downloaded from 10X Genomics. The preprocessing and analysis of RNA and ATAC modalities were performed following the steps outlined for the CD34+ bone marrow data. Cell-type annotation was performed using the marker genes in ^59^ and no cell-types were excluded from the analysis. A total of 11543 cells were retained after all the filtering steps.

##### Lung adenocarcinoma

Fully annotated count matrices for single-cell profiling of lung adenocarcinoma in patient samples were downloaded from ^32^. All non-immune cells contained in the data-set were used in the analyses, comprising a total of 4770 cells. Each patient sample was individually processed by performing normalization on raw counts, followed by log-transformation. Following the procedure outlined in the manuscript, 1500 most highly variable genes were identified and principal components were computed from the expression of these genes.

##### Bone marrow mononuclear cells scATAC-seq dataset

Fragment files for single-cell ATAC-seq data of bone marrow mononuclear cells and CD34+ cells (total of 5 samples) and the respective cell-type annotations were downloaded from GEO^19^. All the cells described in the manuscript^19^ were used except for T-cells since they do not differentiate in the bone marrow. The preprocessing and peak-calling followed the same procedure outlined for the ATAC modality of the CD34+ bone marrow Multiome dataset. A total of 19438 cells were retained after all the filtering steps.

#### Metacell identification

SEACells was applied with default parameters to PBMC and CD34+ bone marrow datasets. The number of metacells were chosen outlined in the “Notes about number of metacells” section. The number of metacells for each sample were: (i) PBMC multiome: 100, (ii) CD34+ bone marrow multiome: 85, (iii) T-cell depleted bone marrow multiome: 100, (iv) Single-cell ATAC-seq of bone marrow mononuclear cells: 270. SEACells was applied separately for the RNA and ATAC modalities of the multiome datasets, using the PCA and SVD representations respectively. Metacell raw counts for different datasets were determined as described in the “Metacell identification” section. Metacell counts were normalized as described in “Metacell normalization”.

##### Comparison of metacells from two modalities using PBMC multiome data

We used the paired nature of multiome data to compare consistency of metacells identified between the two modalities. Due to the clear separation between cell types, PBMC multiome dataset was used for this analysis to verify whether relationships between metacells within and across cell-types were consistent between the two data modalities. We checked whether single-cell groups derived using ATAC modality could be applied to the RNA modality and retain cell-type consistency.

We first computed the aggregated RNA metacell matrix. We then computed a second aggregated gene expression using the single-cell groups from ATAC modality instead of the RNA modality. We jointly normalized the two aggregated matrices, identified highly variable genes, computed principal components and visualized data using UMAPs (**Supplementary Fig. 3A**). No batch correction was used for this analysis. We repeated the same procedure using aggregated peak counts from ATAC and RNA metacells (**Supplementary Fig. 3B**).

##### Peak-calling, gene scores and gene accessibility in CD34+ bone marrow dataset

Peak calling, peak gene associations, gene score computation and gene accessibility scores were determined as described in the “Toolkit for scATAC analysis” section.

Since only scATAC is available for the BMMC dataset, peak-gene associations identified using the CD34+ multiome dataset were used for the gene accessibility analysis.

### Robustness of SEACells algorithm

Due to its more challenging continuous nature, we used the CD34+ bone marrow data for assessing the robustness of SEACells algorithm. With a series of cell states spanning continuous trajectories, this dataset provides a greater challenge for test of robustness compared to the well-separated PBMC multiome dataset.

#### Robustness to different initializations

Since the max-min sampling procedure relies on a random seed, we first tested the robustness of SEACells algorithm to different initializations. We consider the procedure robust if cells are consistently assigned to the same cell type across different runs. Normalized metacell RNA matrices were determined separately for each initialization. To compare a pair of initializations, we first concatenated the normalized matrices and then computed diffusion components using both groups of metacells. Briefly, PCA was used to derive a low dimensional embedding of the concatenated metacell matrix using the single-cell data determined highly variable genes. Diffusion components were determined using PCs as the input and a permissive 10 diffusion components were used for downstream analysis. For each metacell in an initialization, nearest metacell neighbors from the alternative initialization were computed. Two metacells from different initializations were considered equivalent if they were mutually in each others’ top two nearest neighbors (**Supplementary Fig. 5A**). Neighborhood computation was performed using diffusion components and diffusion distance^5^. We quantified the comparison for each pair of initializations by computing the proportion of mapped metacells from alternate initializations with matching cell-types (**Supplementary Fig. 5B**).

A similar procedure was used to test the robustness of the ATAC modality using aggregated ArchR gene scores instead of gene expression as inputs.

#### Robustness of different numbers of metacells

Robustness to different numbers of metacells (**Supplementary Fig. 5C,D**) were determined using the same procedure outlined above, using the CD34+ RNA modality.

### Metacells methods comparison

#### Baran et. al. MetaCell

MetaCell^9^ approach uses a non-parametric graph algorithm to partition scRNA-seq data into distinct metacells. This algorithm constructs a balanced kNN graph, which is subsampled multiple times into dense subgraphs in order to determine metacell partitions. Outlier cells are identified, and the final output is the assignment of cells to metacells. MetaCell was run using the default processing steps outlined in https://tanaylab.github.io/metacell/articles/a-basic_pbmc8k.html with raw count data.

We ran Baran et. al. metacell on three scRNA-seq datasets using default parameters - CD34+ bone marrow, PBMC and lung adenocarcinoma in order to evaluate the performance in the contexts of continuous differentiation, discrete cell states and a cancer dataset, respectively. For each dataset, Baran et. al. metacell automatically infers the number of metacells and discards a subset of the data as outliers. To compare faithfully across methods, we used the same number of partitions as input to SEACells and Super-cells on the same subset of data.

To apply Baran et. al. metacells to scATAC data, the peak count matrices were modified to be used as input by mapping peaks to the nearest gene and aggregating all peaks within each gene to create a pseudo cell-by-gene count matrix as input. Following this representation, we ran Baran et. al. metacell on the CD34+ bone marrow and PBMC scATAC-seq datasets with default parameters.

#### Super-cells

Super-cells use the walktrap algorithm to partition nodes in a single-cell graph into a predefined number of super-cells^10^. Therefore, similar to SEACells, the number of metacells is a parameter to the Super-cells algorithm. Super-cells constructs a single-cell graph, placing edges between cells with similar transcriptomic profiles, and merges nodes which are highly connected. Effectively, Super-cells can be viewed as fine resolution community-detection based clustering

Super-cells was run using the default parameters specified in https://github.com/GfellerLab/SuperCell, with the graining level chosen to obtain the same number of partitions as those obtained by Baran et. al metacells, in order to compare methods across similar levels of granularity. We ran Super-cells on the CD34+ bone marrow and PBMC scRNA-seq datasets using default parameters.

We applied Super-cells to CD34+ bone marrow and PBMC scATAC-seq data using the aggregation approach we used for running Baran et. al. approach.

### Metrics for metacell benchmarking

We developed a number of metrics to evaluate the quality of identified metacells and quantify the differences between different metacell approaches. Given that metacells represent distinct cell-states of the biological system under consideration, inferred metacells should be (i) compact i.e., low variability amongst cells that are aggregated together with most of the variability a result of measurement noise and (ii) well separated from neighboring metacells since distinct metacells should include distinct gene-gene covariation matrices, even if these distinctions are subtle.

We used diffusion components to quantify both the compactness and separation of metacells. Diffusion maps have been used extensively to robustly and faithfully represent the phenotypic manifold using single-cell data^5^. Each diffusion component represents a key axis of biological variance in both continuous trajectories and discrete states and thus provides an ideal platform to quantify metacell qualities.

#### Compactness

Compactness provides a measure of how homogeneous cells within a metacell are. We first compute diffusion components using single-cell data. For each metacell, the variance in each diffusion component dimension is computed across the cells that constitute the metacell. The average variance across components is reported as the compactness. Since diffusion components are by definition orthonormal, we can compute the variance of each component separately. The average variance ensures that the homogeneity of cells that constitute the metacell are measured across all axes of biological variance.

For a metacell, *s*, the compactness, *Compactness*(*s*) is formally defined as follows.

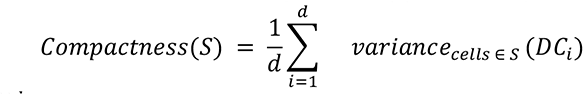

where *DC*∈*R^n×d^* where is the matrix of diffusion components computed using single-cell data.

A high quality metacell should have a low compactness score indicating low variability or equivalently high homogeneity amongst the cells that constitute the metacell.

#### Implementation details

For scRNA-seq, diffusion components are computed based on principal components, and for scATAC-seq, based on the singular value decomposition following preprocessing of single-cell data as described in “Data preprocessing”. The number of components can be chosen by the Eigen gap statistic. We noted that across datasets and modalities, the number of diffusion components ranged from 6-8. For consistency and simplicity, we fixed the number of diffusion components as 10 for all evaluations.

#### Separation

To assess whether metacells are distinct from each other, we evaluated the separation between neighboring metacells using diffusion components. Diffusion components are computed at the single-cell level as described in the “Compactness” section. For each metacell, diffusion embedding is determined as the average of the cells that constitute the metacell. Distance between the metacell and its nearest neighbor is reported as the separation of the metacell. Since diffusion components are a faithful representation of the phenotypic manifold, a greater distance between metacells determined in diffusion space indicates a better separation between them.

#### Cell-type Purity

Cell-type purity is a measure of the consistency of cell-types amongst cells that constitute a metacell and was introduced to assess the quality of Super-cells^10^. Cell-type purity is computed as the proportion of cells which belong to the modal cell-type in a metacell. Note that purity metric is applicable and valid when the biological system under consideration comprises distinct cell-types with distinct functions such as PBMCs. Cell-type purity is not a reliable metric for continuous trajectories since the different cell-types or compartments are merely a partitioning of the trajectory and do not necessarily represent well separated cell-types.

#### Comparison of different metacell approaches using benchmarking metrics

Benchmarking metrics were determined for each metacell for all (data modality, dataset, method) combinations. Cell-type purity was used to assess the quality of PBMC metacells. Different methods were compared using the Wilcoxon rank-sum test. Top performing metacell approaches should have a low score on compactness, high score on separation and high score on cell-type purity.

We compared the metacell approaches using all metacells and separately for metacells in low- and high-density regions to verify that all biologically relevant states are uniformly assessed. We once again used diffusion components to quantify the density of cells. Distance to the 150th neighbor in a single-cell nearest neighbor graph has been demonstrated to be a reasonable approximation for the density in the high dimensional space^55^. We computed the distance to the 150th neighbor for each single-cell using diffusion components. Single cells with densities in the upper quartile of distances were designated as ‘low-density cells’, and similarly, those in the lower quartile of distances were designated as ‘high-density cells’. Analogously, metacells containing these low-density cells were designated as low density metacells, and vice versa for high density metacells. The proportion of all metacells designated as either low or high density were each capped at 30% of all metacells, and these were used as low-density and high-density regions respectively for comparisons (**Supplementary Fig. 14**).

### Characterization of hematopoietic dynamics

#### Palantir application for CD34+ bone marrow data

Palantir^5^ was applied using default parameters using the RNA modality of CD34+ bone marrow Multiome data at single-cell level. Briefly, diffusion components were computed using the adaptive kernel and the number of informative diffusion components (n=7) were identified using the Eigen gap statistic. Palantir was run using these diffusion components using a CD34 high hematopoietic stem cells as the start cell. The terminal states for Erythroid, Lymphoid, Megakaryocyte, Monocytes, cDC and pDC lineages were all set manually. The pseudo-temporal ordering of metacells was computed as the average pseudo-time ordering of the constituent single-cells.

For pseudo-time bins in **Fig. 5D**, cells were categorized into one of forty bins based on their Palantir pseudo-time order, which ranged from 0.0-0.82 for the erythroid lineage. We then created 40 equal sized bins with a bin size of 0.02 and assigned each cell to the respective bin. Fragments that belong to all cells in a bin were pooled and open peaks identified using the Poisson procedure.

#### Accessibility trends

Gene accessibility trends were determined using generalized additive models (GAMs)^60^.A GAM was fit for gene accessibility trend as a function of the Palantir pseudo-time for each gene. Gene accessibility of *g* in cell *i*, *y_gi_* is fit as

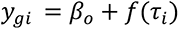

where *i* is a cell along the relevant lineage, *τ_i_* is the Palantir pseudo-temporal ordering of cell *i*. Cubic splines are used as the smoothing functions since they are effective in capturing non-linear relationships. The pseudo-time is then divided into 150 equally sized bins and the smooth trend is derived by using the fit from the Generalized Additive Model to predict the accessibility of the gene at each bin.

#### Gene ontology analysis

Gene ontology analysis (**Fig. 5E**) was performed to identify enriched ontologies in genes with increasing or decreasing accessibility, measuring enrichment using the hypergeometric test. The “c7: immunologic signature” gene sets from Molecular Signature Database (MSigDB) (http://software.broadinstitute.org/gsea/msigdb/index.jsp) was used.

#### Motif enrichments in genes with changing accessibility

Predicted TF binding sites in each peak were determined using FIMO^61^ using default parameters and the cisBP v2 motif database^62^. Hypergeometric tests were used to identify the most enriched motifs in peaks with increasing or decreasing accessibility using all the peaks as the background.

### SEACells application to COVID-19 samples and data integration

#### COVID-19 data preprocessing

Raw counts for single-cell RNA-seq data of peripheral blood mononuclear cells as well as the respective cell-type, disease severity and sample annotations were downloaded from https://covid19cellatlas.org^44^. Cells corresponding to patients annotated as healthy or as critical were used, comprising a total of 96405 cells from 23 healthy patients and 80837 cells from 17 patients. Each sample was individually processed by performing normalization, followed by log-transformation as described in the “Data preprocessing” section. The 1500 most highly variable genes were identified, and principal components were computed from the expression of these genes.

SEACells metacells were also computed separately for each sample using approximately one metacell for every seventy-five single-cells following the procedure described in “Notes about number of metacells”. Following metacell identification, an aggregated metacell X gene expression matrix was computed for each sample.

#### Batch correction

Harmony^45^ was used to perform batch correction across all 40 samples on the metacell aggregated gene expression matrices using default parameters. Harmony (**scanpy.external.pp.harmony_integrate**) was applied to the principal components derived from the top 1500 highly variable genes using default parameters.

Harmony was applied separately at single-cell and metacell levels for comparison (**Supplementary Fig. 20**).

#### Mapping of SEACell metacells between individuals

We mapped metacells across patients to determine consistency. The analysis was performed using the same procedure described in the “Robustness of SEACells algorithm” section. For each pair of patients, Harmony corrected metacell principal components were used for the analysis. Diffusion components were determined using Harmony corrected PCs as the input and a permissive 10 diffusion components were used for downstream analysis. For each metacell in a patient, nearest metacell neighbors from the second patient were computed. Two metacells from different patients were considered equivalent if they were mutually in each others’ top two nearest neighbors (**Supplementary Fig. 19A,B**). We quantified the comparison for each pair of samples by computing the proportion of mapped metacells with matching cell-types (**Supplementary Fig. 19C**).

#### Differential abundance testing of cell-states between healthy individuals and COVID-19 patients

By aggregating single-cells that are most likely a result of technical noise, metacells provide a robust segmentation of the data. Thus, metacells computed per sample thus provide a granular representation for across sample comparison. Metacells are inherently less susceptible to batch effects compared to single-cell data and thus provide a concrete baseline to infer altered cell-state abundances across different conditions (**Supplementary Fig. 20**).

#### Generation of aggregates of metacells in COVID data

While mapping metacells demonstrates consistency between pairs of individuals, the approach does not provide a path to identify *similarities and differences* between healthy individuals and COVID-19 patients. We therefore devised a procedure for seamless comparison across any number of patients and identify enriched and depleted metacells in different conditions.

We re-computed SEACells metacells using the aggregated and batch corrected metacell count matrices for each sample. These second level metacells, or Meta^2^cells, therefore contain metacells across healthy and critical patient samples. To compute Meta^2^cells, we ran the algorithm asking for approximately one Meta^2^cell for every ten metacells, since the dataset was already highly summarized in the first round of aggregation.

To summarize the cell-type annotations of cells in a constituent Meta^2^cell, the modal cell-type of constituent cells was chosen if the purity was greater than 80%, otherwise the cell-type was denoted as ‘Mixed’.

#### Tests for differential abundance of cell-states in COVID-19 patients

The Meta^2^cells computed across healthy individual and critical patients define cell-states, each of which may be more strongly associated with healthy or diseased state. We computed the proportion of COVID-19 metacells in each Meta^2^cell, providing a measure of differential abundance of cell-state in COVID-19 patients. We then devised a permutation test to assess the significance of these differential abundances.

First, the assignment of metacells to Meta^2^cell were randomly permuted. This ensures that the number of metacells assigned to each Meta^2^cell does not change but the constituent metacells and their associated healthy/COVID-19 labels will be permuted providing a representative background distribution. Next, the proportion of metacells derived from COVID-19 samples assigned to each Meta^2^cell was computed. This procedure was repeated for 5000 trials of permutations, and a null distribution on COVID-19 enriched metacell proportions was derived for each Meta^2^cell. The null distribution is then used to compute a *p*-value and a cell-state is nominated to be significantly enriched in COVID-19 is *p*-value < 0.1

#### Gene signatures of enriched cell-states

To assess the biological distinctions between healthy and diseased Meta^2^cell states, we identified the differentially expressed genes for each Meta^2^cell by comparing against other Meta^2^cells of the cell-type using **scanpy.tl.rank_genes_groups**.

#### Single-cell differential abundance testing

We used the extensively used Milo^15^ to perform differential abundance testing at the single-cell level and compared the results to differential abundance testing using metacells. We applied MiloR to all cells from 23 healthy patients and 17 critical patients. MiloR typically accepts a SingleCellExperiment object as input. However, due to memory constraints in passing raw counts for all 177242 cells, we provided MiloR with the pre-computed batch-corrected principal components, annotated with the sample of origin and sample condition. Default parameters as specified in the Milo vignettes were then used to compute neighborhoods as well as their differential abundances. All neighborhoods with at least 80% CD4 cell-type purity were selected for downstream analysis, yielding 276 neighborhoods.

Gene signatures identified in the SEACells metacells of interest were used to compute a gene signature score for each MiloR neighborhood. The gene signature score was computed for each cell by summing across the expression z-scores of the signature genes. Gene signature scores at the neighborhood level were computed by average the scores of single-cells that constitute the neighborhood. To assess whether the cell-states highlighted in **Fig. 6D** could be identified using differential abundance testing at single-cell level, we compared the Milo neighborhood gene signature scores with the gene scores derived using SEACells Meta^2^cell (**Supplementary Fig. 22C**).

## References

1. Regev, A. et al. The Human Cell Atlas. Elife 6, doi:10.7554/eLife.27041 (2017).

2. Rozenblatt-Rosen, O. et al. The Human Tumor Atlas Network: Charting Tumor Transitions across Space and Time at Single-Cell Resolution. Cell 181, 236–249, doi:10.1016/j.cell.2020.03.053 (2020).

3. Wagner, A. et al. Metabolic modeling of single Th17 cells reveals regulators of autoimmunity. Cell 184, 4168–4185 e4121, doi:10.1016/j.cell.2021.05.045 (2021).

4. Bendall, S. C. et al. Single-cell trajectory detection uncovers progression and regulatory coordination in human B cell development. Cell 157, 714–725, doi:10.1016/j.cell.2014.04.005 (2014).

5. Setty, M. et al. Characterization of cell fate probabilities in single-cell data with Palantir. Nat Biotechnol 37, 451–460, doi:10.1038/s41587-019-0068-4 (2019).

6. Haghverdi, L., Buettner, F. & Theis, F. J. Diffusion maps for high-dimensional single-cell analysis of differentiation data. Bioinformatics 31, 2989–2998, doi:10.1093/bioinformatics/btv325 (2015).

7. Cao, J. et al. The single-cell transcriptional landscape of mammalian organogenesis. Nature 566, 496–502, doi:10.1038/s41586-019-0969-x (2019).

8. May, G. et al. Dynamic analysis of gene expression and genome-wide transcription factor binding during lineage specification of multipotent progenitors. Cell Stem Cell 13, 754–768, doi:10.1016/j.stem.2013.09.003 (2013).

9. Baran, Y. et al. MetaCell: analysis of single-cell RNA-seq data using K-nn graph partitions. Genome Biol 20, 206, doi:10.1186/s13059-019-1812-2 (2019).

10. Bilous, M. et al. Super-cells untangle large and complex single-cell transcriptome networks. bioRxiv, 2021.2006.2007.447430, doi:10.1101/2021.06.07.447430 (2021).

11. Azizi, E. et al. Single-Cell Map of Diverse Immune Phenotypes in the Breast Tumor Microenvironment. Cell 174, 1293–1308 e1236, doi:10.1016/j.cell.2018.05.060 (2018).

12. Haghverdi, L., Lun, A. T. L., Morgan, M. D. & Marioni, J. C. Batch effects in single-cell RNA-sequencing data are corrected by matching mutual nearest neighbors. Nat Biotechnol 36, 421–427, doi:10.1038/nbt.4091 (2018).

13. Hao, Y. et al. Integrated analysis of multimodal single-cell data. Cell 184, 3573–3587 e3529, doi:10.1016/j.cell.2021.04.048 (2021).

14. Levine, J. H. et al. Data-Driven Phenotypic Dissection of AML Reveals Progenitor-like Cells that Correlate with Prognosis. Cell 162, 184–197, doi:10.1016/j.cell.2015.05.047 (2015).

15. Dann, E., Henderson, N. C., Teichmann, S. A., Morgan, M. D. & Marioni, J. C. Differential abundance testing on single-cell data using k-nearest neighbor graphs. Nat Biotechnol 40, 245–253, doi:10.1038/s41587-021-01033-z (2022).

16. Hart, Y. et al. Inferring biological tasks using Pareto analysis of high-dimensional data. Nat Methods 12, 233-235, 233 p following 235, doi:10.1038/nmeth.3254 (2015).

17. Bauckage, C., Kersting, K., Hoppe, F. & Thurau, C. in Workshop New Challenges in Neural Computation (2015).

18. Ma, S. et al. Chromatin Potential Identified by Shared Single-Cell Profiling of RNA and Chromatin. Cell 183, 1103–1116 e1120, doi:10.1016/j.cell.2020.09.056 (2020).

19. Granja, J. M. et al. Single-cell multiomic analysis identifies regulatory programs in mixed-phenotype acute leukemia. Nat Biotechnol 37, 1458–1465, doi:10.1038/s41587-019-0332-7 (2019).

20. Trevino, A. E. et al. Chromatin and gene-regulatory dynamics of the developing human cerebral cortex at single-cell resolution. Cell 184, 5053–5069 e5023, doi:10.1016/j.cell.2021.07.039 (2021).

21. Buenrostro, J. D., Giresi, P. G., Zaba, L. C., Chang, H. Y. & Greenleaf, W. J. Transposition of native chromatin for fast and sensitive epigenomic profiling of open chromatin, DNA-binding proteins and nucleosome position. Nat Methods 10, 1213–1218, doi:10.1038/nmeth.2688 (2013).

22. Hashimoto, H. et al. Structural Basis for the Versatile and Methylation-Dependent Binding of CTCF to DNA. Mol Cell 66, 711–720 e713, doi:10.1016/j.molcel.2017.05.004 (2017).

23. Granja, J. M. et al. ArchR is a scalable software package for integrative single-cell chromatin accessibility analysis. Nat Genet 53, 403–411, doi:10.1038/s41588-021-00790-6 (2021).

24. Schep, A. N., Wu, B., Buenrostro, J. D. & Greenleaf, W. J. chromVAR: inferring transcription-factor-associated accessibility from single-cell epigenomic data. Nat Methods 14, 975–978, doi:10.1038/nmeth.4401 (2017).

25. Schnell, A. et al. Stem-like intestinal Th17 cells give rise to pathogenic effector T cells during autoimmunity. Cell 184, 6281–6298 e6223, doi:10.1016/j.cell.2021.11.018 (2021).

26. Gaublomme, J. T. et al. Single-Cell Genomics Unveils Critical Regulators of Th17 Cell Pathogenicity. Cell 163, 1400–1412, doi:10.1016/j.cell.2015.11.009 (2015).

27. Yukawa, M. et al. AP-1 activity induced by co-stimulation is required for chromatin opening during T cell activation. J Exp Med 217, doi:10.1084/jem.20182009 (2020).

28. Laurenti, E. & Gottgens, B. From haematopoietic stem cells to complex differentiation landscapes. Nature 553, 418–426, doi:10.1038/nature25022 (2018).

29. Pearce, E. L. et al. Control of effector CD8+ T cell function by the transcription factor Eomesodermin. Science 302, 1041–1043, doi:10.1126/science.1090148 (2003).

30. Vallabhapurapu, S. & Karin, M. Regulation and function of NF-kappaB transcription factors in the immune system. Annu Rev Immunol 27, 693–733, doi:10.1146/annurev.immunol.021908.132641 (2009).

31. Keren-Shaul, H. et al. MARS-seq2.0: an experimental and analytical pipeline for indexed sorting combined with single-cell RNA sequencing. Nat Protoc 14, 1841–1862, doi:10.1038/s41596-019-0164-4 (2019).

32. Laughney, A. M. et al. Regenerative lineages and immune-mediated pruning in lung cancer metastasis. Nat Med 26, 259–269, doi:10.1038/s41591-019-0750-6 (2020).

33. Gonzalez, A. J., Setty, M. & Leslie, C. S. Early enhancer establishment and regulatory locus complexity shape transcriptional programs in hematopoietic differentiation. Nat Genet 47, 1249–1259, doi:10.1038/ng.3402 (2015).

34. Lara-Astiaso, D. et al. Immunogenetics. Chromatin state dynamics during blood formation. Science 345, 943–949, doi:10.1126/science.1256271 (2014).

35. Velten, L. et al. Human haematopoietic stem cell lineage commitment is a continuous process. Nat Cell Biol 19, 271–281, doi:10.1038/ncb3493 (2017).

36. Weinreb, C., Rodriguez-Fraticelli, A., Camargo, F. D. & Klein, A. M. Lineage tracing on transcriptional landscapes links state to fate during differentiation. Science 367, doi:10.1126/science.aaw3381 (2020).

37. Tusi, B. K. et al. Population snapshots predict early haematopoietic and erythroid hierarchies. Nature 555, 54–60, doi:10.1038/nature25741 (2018).

38. Elmentaite, R., Dominguez Conde, C., Yang, L. & Teichmann, S. A. Single-cell atlases: shared and tissue-specific cell types across human organs. Nat Rev Genet, doi:10.1038/s41576-022-00449-w (2022).

39. Jardine, L. et al. Blood and immune development in human fetal bone marrow and Down syndrome. Nature 598, 327–331, doi:10.1038/s41586-021-03929-x (2021).

40. Elmentaite, R. et al. Cells of the human intestinal tract mapped across space and time. Nature 597, 250–255, doi:10.1038/s41586-021-03852-1 (2021).

41. Sikkema, L. et al. An integrated cell atlas of the human lung in health and disease. bioRxiv, 2022.2003.2010.483747, doi:10.1101/2022.03.10.483747 (2022).

42. Qiu, C. et al. Systematic reconstruction of cellular trajectories across mouse embryogenesis. Nat Genet 54, 328–341, doi:10.1038/s41588-022-01018-x (2022).

43. Srivatsan, S. R. et al. Embryo-scale, single-cell spatial transcriptomics. Science 373, 111–117, doi:10.1126/science.abb9536 (2021).

44. Stephenson, E. et al. Single-cell multi-omics analysis of the immune response in COVID-19. Nat Med 27, 904–916, doi:10.1038/s41591-021-01329-2 (2021).

45. Korsunsky, I. et al. Fast, sensitive and accurate integration of single-cell data with Harmony. Nat Methods 16, 1289–1296, doi:10.1038/s41592-019-0619-0 (2019).

46. Traag, V. A., Waltman, L. & van Eck, N. J. From Louvain to Leiden: guaranteeing well-connected communities. Sci Rep 9, 5233, doi:10.1038/s41598-019-41695-z (2019).

47. Sposito, B. et al. The interferon landscape along the respiratory tract impacts the severity of COVID-19. Cell 184, 4953–4968 e4916, doi:10.1016/j.cell.2021.08.016 (2021).

48. Pan, J. et al. A novel chemokine ligand for CCR10 and CCR3 expressed by epithelial cells in mucosal tissues. J Immunol 165, 2943–2949, doi:10.4049/jimmunol.165.6.2943 (2000).

49. Argelaguet, R. et al. MOFA+: a statistical framework for comprehensive integration of multi-modal single-cell data. Genome Biol 21, 111, doi:10.1186/s13059-020-02015-1 (2020).

50. Wu, S. J. et al. Single-cell CUT&Tag analysis of chromatin modifications in differentiation and tumor progression. Nat Biotechnol 39, 819–824, doi:10.1038/s41587-021-00865-z (2021).

51. Bartosovic, M., Kabbe, M. & Castelo-Branco, G. Single-cell CUT&Tag profiles histone modifications and transcription factors in complex tissues. Nat Biotechnol 39, 825–835, doi:10.1038/s41587-021-00869-9 (2021).

52. Zeller, P. et al. Hierarchical chromatin regulation during blood formation uncovered by single-cell sortChIC. bioRxiv, 2021.2004.2026.440606, doi:10.1101/2021.04.26.440606 (2021).

53. Wolf, F. A., Angerer, P. & Theis, F. J. SCANPY: large-scale single-cell gene expression data analysis. Genome Biol 19, 15, doi:10.1186/s13059-017-1382-0 (2018).

54. Cusanovich, D. A. et al. The cis-regulatory dynamics of embryonic development at single-cell resolution. Nature 555, 538–542, doi:10.1038/nature25981 (2018).

55. van Dijk, D. et al. Recovering Gene Interactions from Single-Cell Data Using Data Diffusion. Cell 174, 716–729 e727, doi:10.1016/j.cell.2018.05.061 (2018).

56. Zhang, Y. et al. Model-based analysis of ChIP-Seq (MACS). Genome Biol 9, R137, doi:10.1186/gb-2008-9-9-r137 (2008).

57. Genomics, X. CellRanger ARC, <https://support.10xgenomics.com/single-cell-multiome-atac-gex/software/pipelines/latest/what-is-cell-ranger-arc?src=social&lss=linkedin&cnm=soc-li-ra_g-program-li-ra_g-program&cid=7011P000000y072> (2021).

58. Wolock, S. L., Lopez, R. & Klein, A. M. Scrublet: Computational Identification of Cell Doublets in Single-Cell Transcriptomic Data. Cell Syst 8, 281–291 e289, doi:10.1016/j.cels.2018.11.005 (2019).

59. Satija, R., Farrell, J. A., Gennert, D., Schier, A. F. & Regev, A. Spatial reconstruction of single-cell gene expression data. Nat Biotechnol 33, 495–502, doi:10.1038/nbt.3192 (2015).

60. Hastie, T. & Tibshirani, R. Generalized Additive-Models - Some Applications. J Am Stat Assoc 82, 371–386, doi:Doi 10.2307/2289439 (1987).

61. Grant, C. E., Bailey, T. L. & Noble, W. S. FIMO: scanning for occurrences of a given motif. Bioinformatics 27, 1017–1018, doi:10.1093/bioinformatics/btr064 (2011).

62. Weirauch, M. T. et al. Determination and inference of eukaryotic transcription factor sequence specificity. Cell 158, 1431–1443, doi:10.1016/j.cell.2014.08.009 (2014).

