## Supplementary Figures for "SEACells: Inference of transcriptional and epigenomic cellular states from single-cell genomics data"

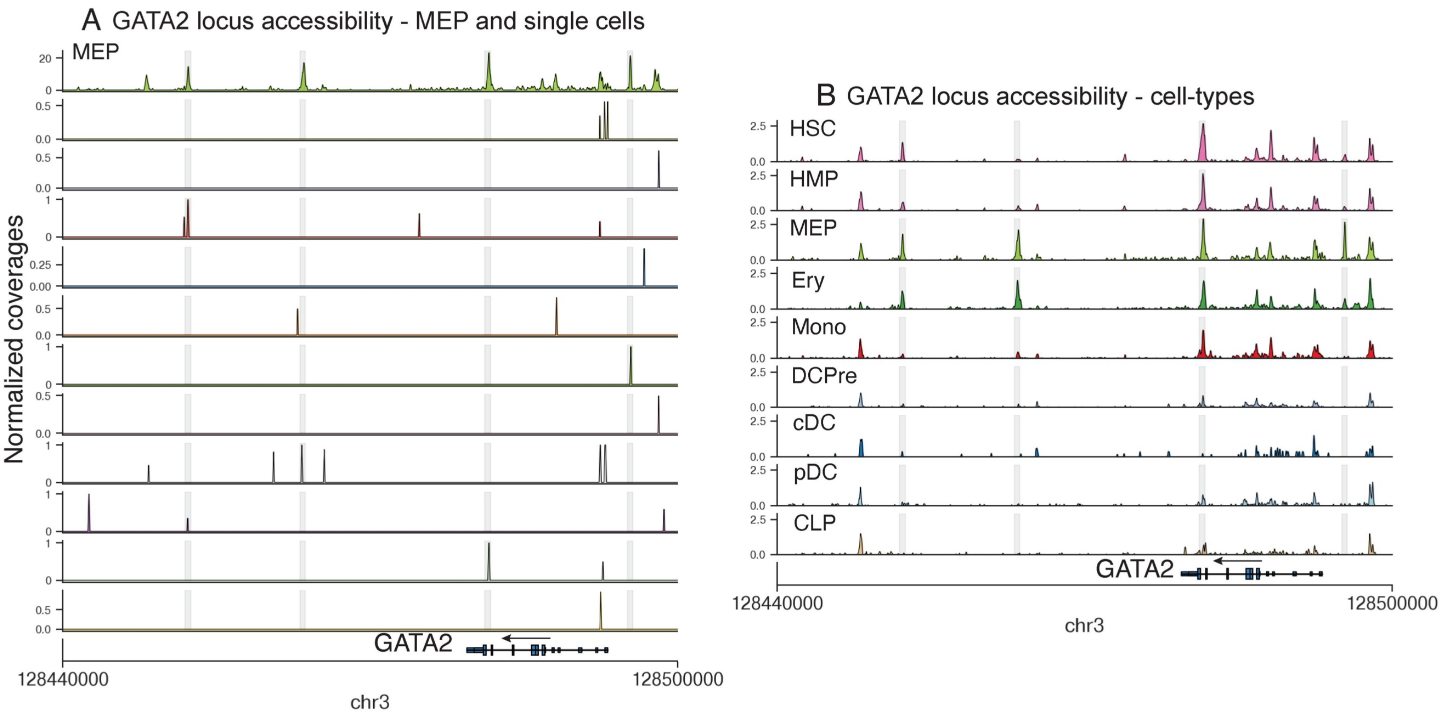


**Supplementary Fig. 1: *GATA2* locus accessibility.**

A. *GATA2* locus accessibility profiles of a random sample of single MEP cells, highlighting the noise and sparsity of single-cell ATAC-seq data. Top row represents an aggregate of 384 MEP cells; each other row represents an individual cell.

B. Accessibility landscape of the *GATA2* locus, across all hematopoietic cell types.

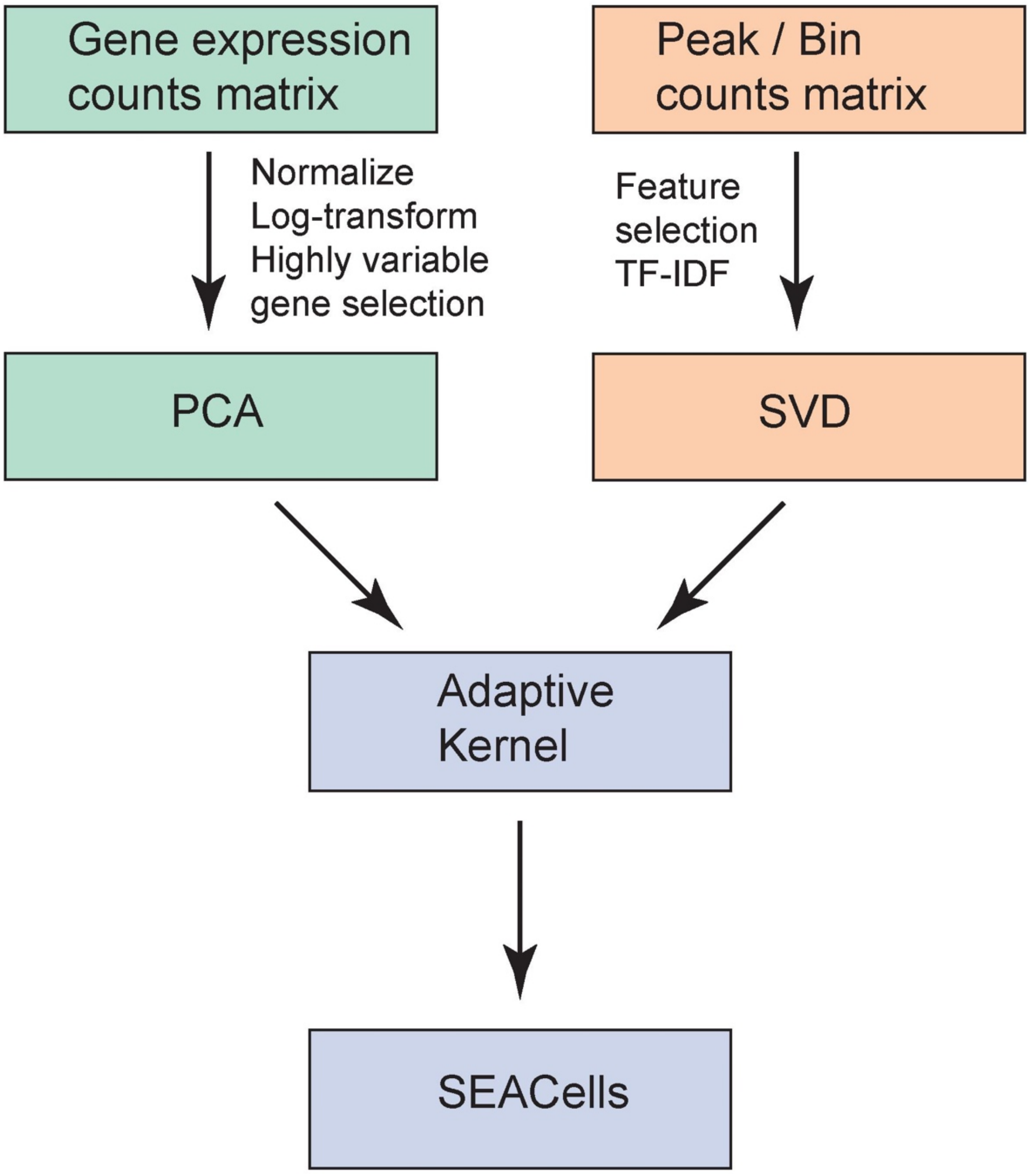


**Supplementary Fig. 2: SEACells workflow for scRNA-seq and scATAC-seq data.**

scRNA-seq and scATAC-seq count matrices are preprocessed differently, using procedures appropriate for the noise and biases of each data type. Principal component analysis (PCA) and singular value decomposition (SVD) are then used to generate a lower-dimensional representation for RNA and ATAC data, respectively. Next, a density adaptive kernel is constructed using the reduced dimensional representations, and input to the SEACells algorithm for metacell construction.

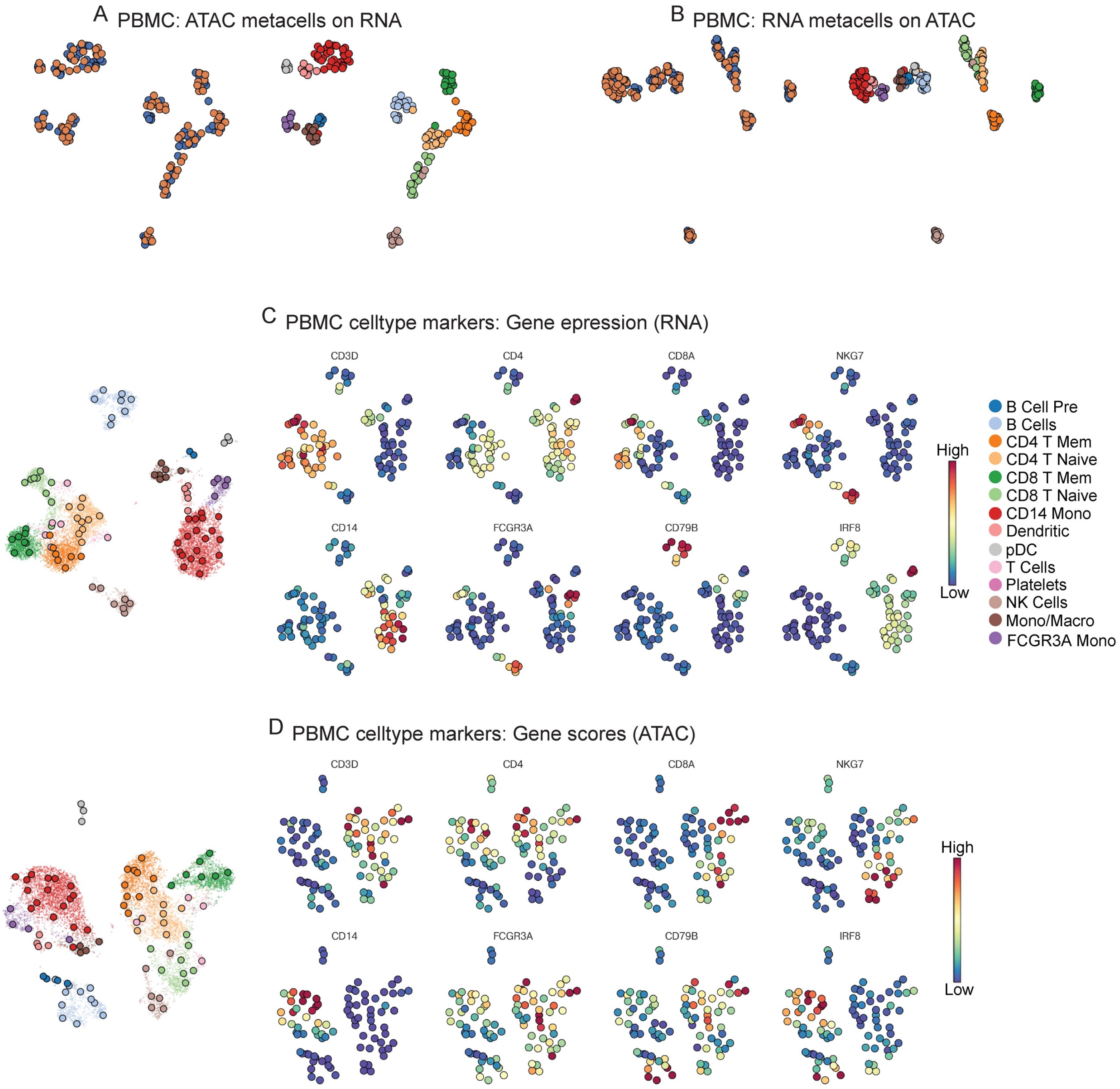


**Supplementary Fig. 3: Metacells are consistent across data modalities and enable cell type identification in PBMC data.**

A,B. UMAPs based on combined ATAC and RNA metacells derived from the PBMC multiome dataset. ATAC metacells were projected on RNA metacells (A) or RNA metacells were projected on ATAC metacells (B) (Methods).

C. Expression of key PBMC cell type markers per RNA metacell.

D. Gene scores of key PBMC cell type markers per ATAC metacell

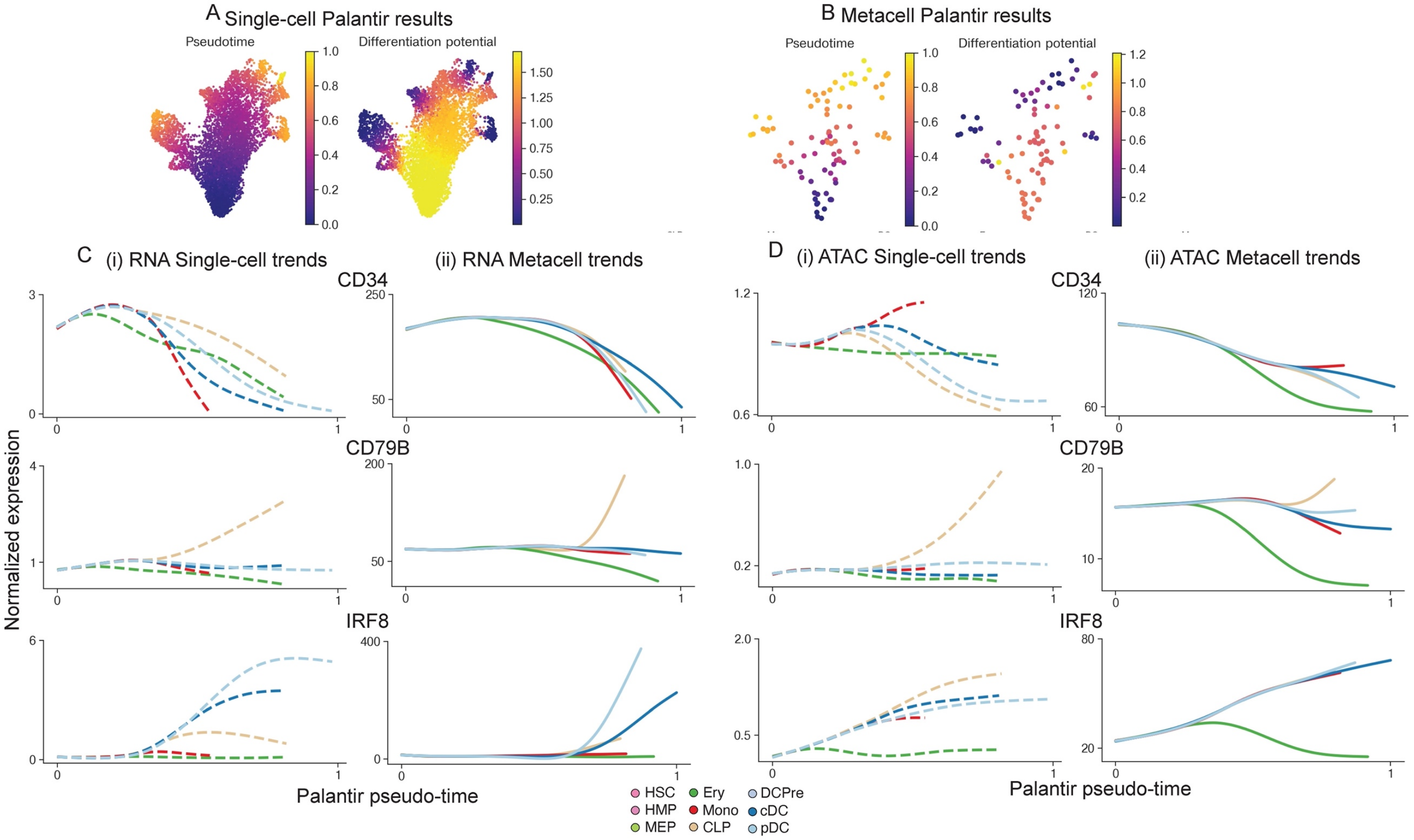


**Supplementary Fig. 4: Gene expression and accessibility trends during human hematopoiesis.**

A. Single-cell RNA UMAP of a CD34+ bone marrow multiome dataset. Cells colored by pseudotime and differentiation potential values, computed by Palantir on single cells using RNA data.

B. Metacell RNA UMAP of dataset in (A). Cells colored by pseudotime and differentiation potential values, computed by Palantir on RNA data corresponding to ATAC-derived metacells.

C. Gene expression trends at single-cell (left) and metacell (right) resolutions. MAGIC-imputed data was used for plotting single-cell trends.

D. Same as (C), for gene accessibility. CD34 accessibility trends from single-cell data do not monotonically decrease across all lineages due to noise in single-cell ATAC measurements.

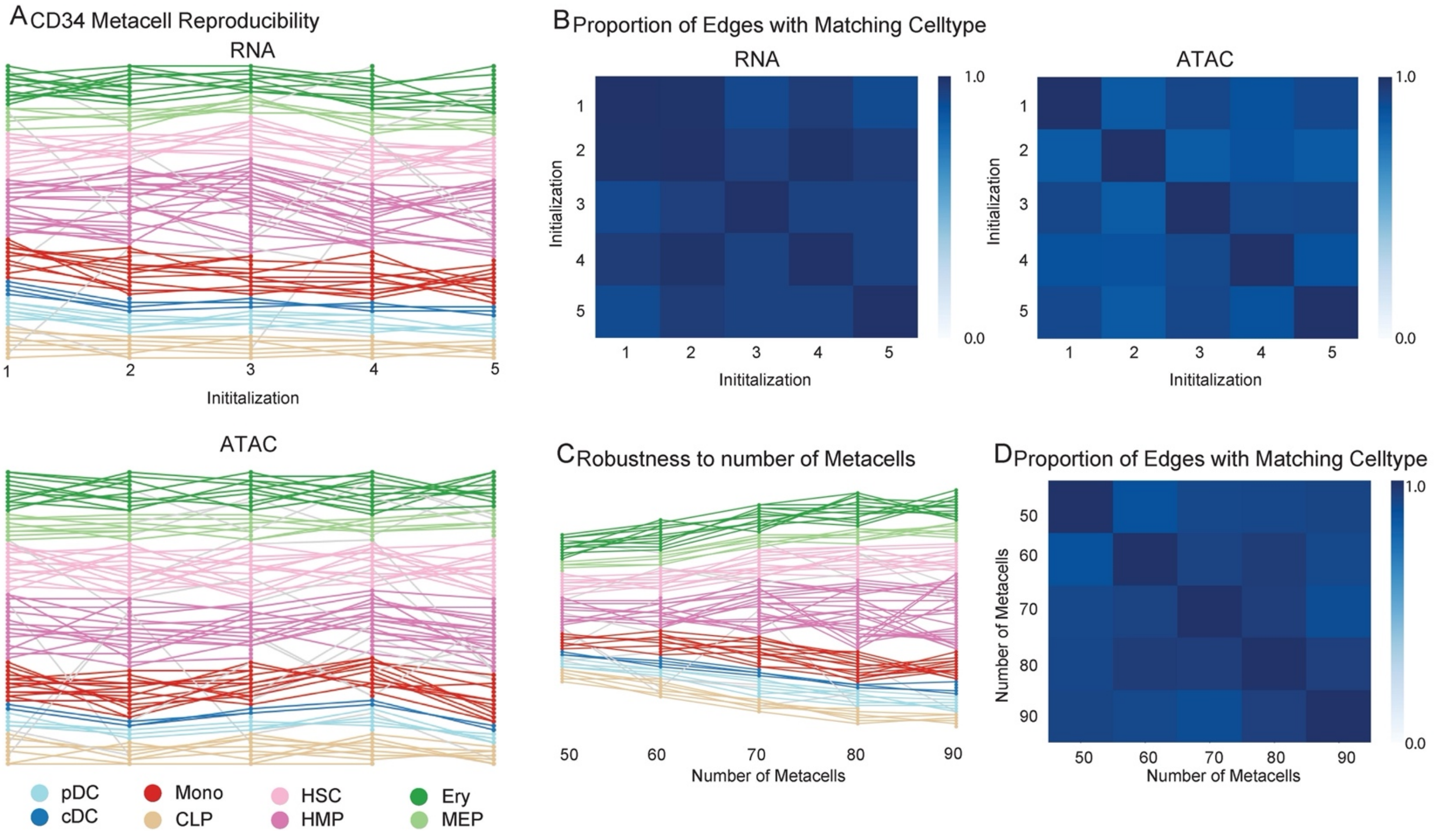


**Supplementary Fig. 5: SEACells metacells are robust and reproducible**

A. Robustness and reproducibility of SEACells metacells to different initializations in CD34 RNA modality (top) and ATAC modality (bottom). Edges connect mutually neighboring metacells between different successive initializations. The edge is colored by cell type when both mutually neighboring metacells agree and grey otherwise.

B. Proportion of mutually nearest neighboring metacells that belong to the same cell type for different waypoint initializations, for the CD34 RNA (left) and ATAC (right) modalities.

C. Robustness to number of metacells. Edges connect mutually neighboring metacells between different SEACells runs using CD34 RNA modality.

D. Proportion of mutually nearest neighboring metacells that belong to the same cell type for CD34 RNA modality, for varying numbers of metacells.

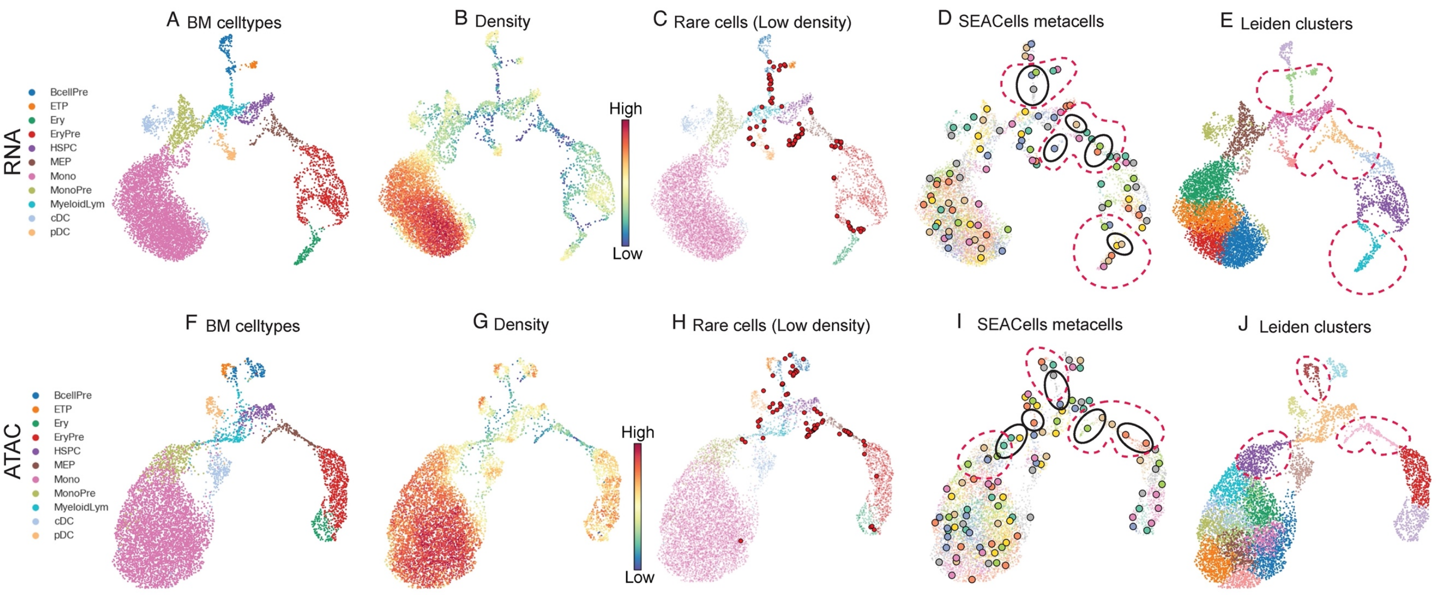


**Supplementary Fig. 6: SEACells identify rare intermediate cell states in continuous trajectories**

A. Single-cell RNA UMAP of T-cell depleted bone marrow cells from a healthy human donor.
B. UMAP colored by cell density.
C. Cells in low-density regions (bottom percentile of density) are highlighted in red
D. SEACells metacells are highlighted in some low-density regions in black. Dotted red lines represent the Leiden clusters that encompass all the underlying metacells.
E. UMAP colored by Leiden clusters
F–I. Same as (A–E), for ATAC modality of T-cell depleted bone marrow cells from a healthy human donor.

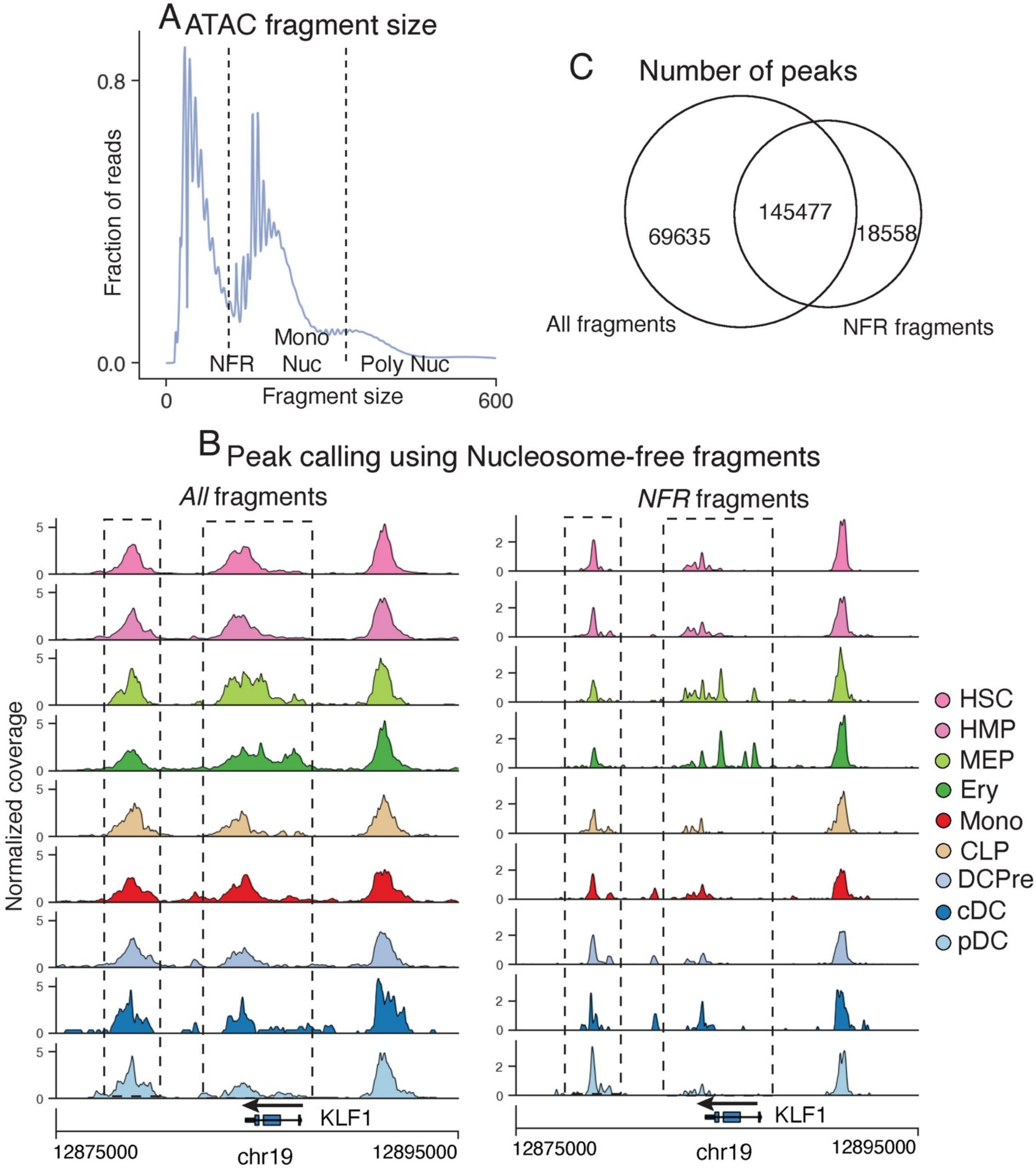


#### Supplementary Fig. 7: scATAC-seq peak calling

A. ATAC-seq fragments follow a characteristic distribution with well-defined modes. The first mode (<147 bp) represents nucleosome-free (NFR) fragments, and subsequent modes represent mono- and poly-nucleosomes.

B. Accessibility landscape of erythroid factor *KLF1* by cell type, using all ATAC fragments (left) or NFR fragments <147 bp (right) to determine coverage. Using NFR fragments substantially improves the resolution of peaks and inferred regulatory elements (dotted boxes).

C. Venn diagram of number of peaks called using NFR or all ATAC fragments in CD34+ multiome data.

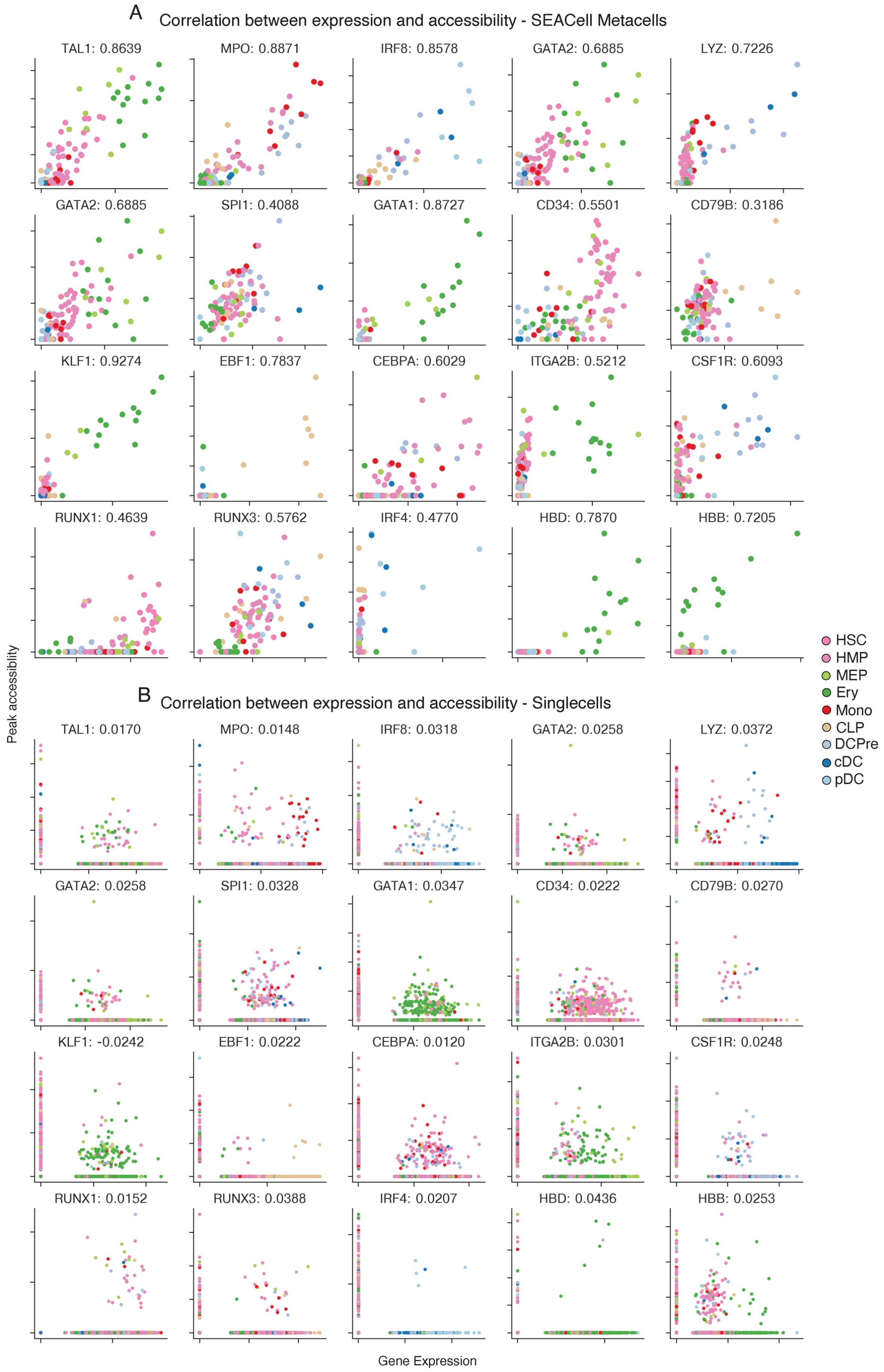

A
Correlation between expression and accessibility - SEACell Metacells
TAL1: 0.8639
MPO: 0.8871
IRF8: 0.8578
GATA2: 0.6885
LYZ: 0.7226
GATA2: 0.6885
SPI1: 0.4088
GATA1: 0.8727
CD34: 0.5501
CD79B: 0.3186
KLF1: 0.9274
EBF1: 0.7837
CEBPA: 0.6029
ITGA2B: 0.5212
CSF1R: 0.6093
RUNX1: 0.4639
RUNX3: 0.5762
IRF4: 0.4770
HBD: 0.7870
HBB: 0.7205
B
Correlation between expression and accessibility - Singlecells
TAL1: 0.0170
MPO: 0.0148
IRF8: 0.0318
GATA2: 0.0258
LYZ: 0.0372
GATA2: 0.0258
SPI1: 0.0328
GATA1: 0.0347
CD34: 0.0222
CD79B: 0.0270
KLF1: -0.0242
EBF1: 0.0222
CEBPA: 0.0120
ITGA2B: 0.0301
CSF1R: 0.0248
RUNX1: 0.0152
RUNX3: 0.0388
IRF4: 0.0207
HBD: 0.0436
HBB: 0.0253
Peak accessibility
Gene Expression
HSC
HMP
MEP
Ery
Mono
CLP
DCPre
cDC
pDC

**Supplementary Fig. 8: Comparison of peak-gene correlations using SEACells metacells and single cells**

A. Relationship between metacell-aggregated gene expression and accessibility of the most correlated peak for a selection of key hematopoietic genes, computed on the CD34+ multiome data. Spearman correlations appear next to the gene symbol.
B. Same as in (A), but at the single-cell level.

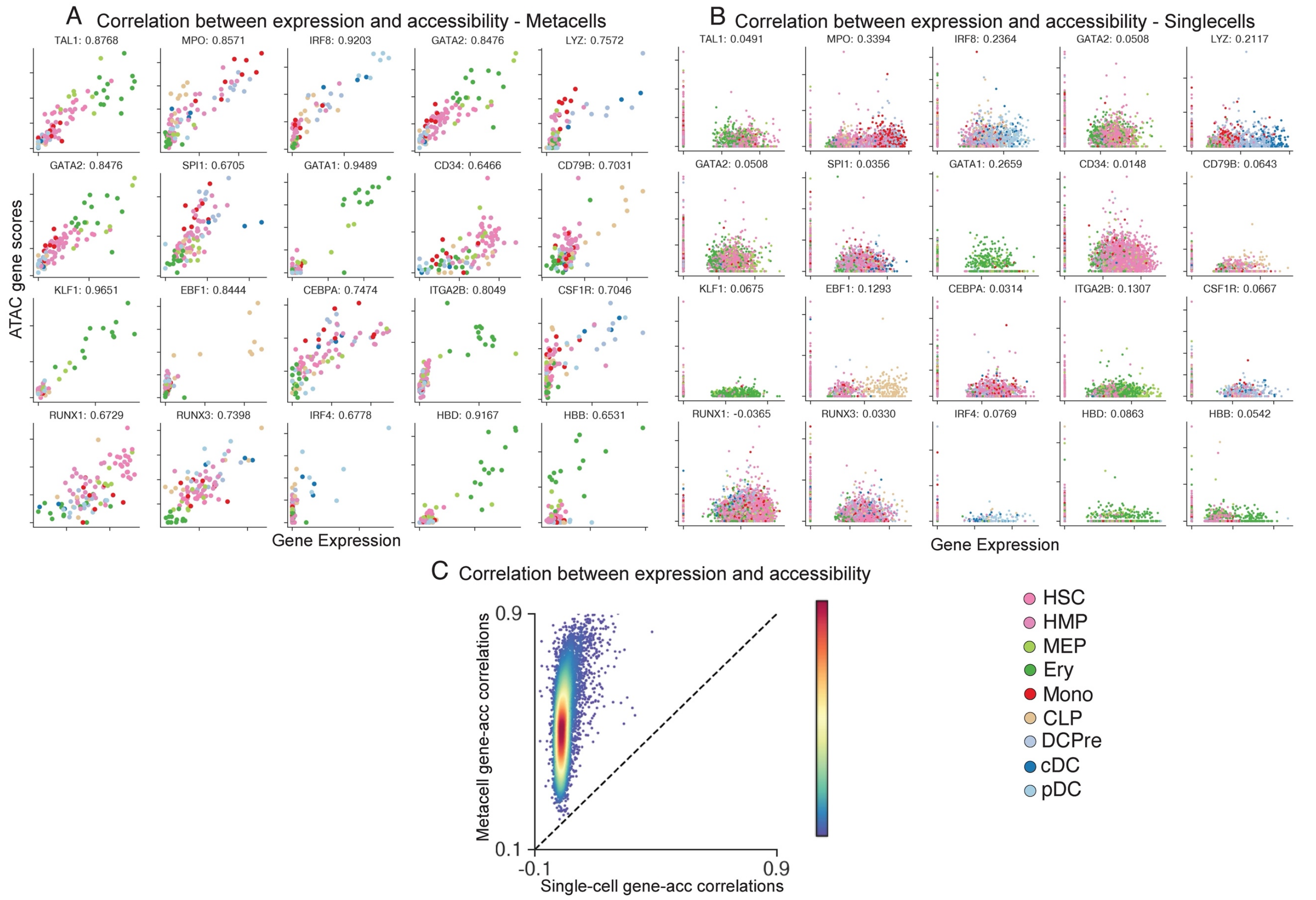


**Supplementary Fig. 9: Comparison of ATAC gene scores using SEACell metacells and single cells**

A. Relationship between metacell-aggregated gene expression and ATAC gene scores for a selection of key hematopoietic genes, computed on the CD34+ multiome data. Gene scores for metacells were computed by aggregating peaks that correlate significantly with expression. Spearman correlations appear next to the gene symbol.

B. Same as in (A), but at single-cell level. Gene scores for single-cell data were computed using ArchR.

C. Spearman correlations between gene expression and ATAC gene scores, plotted for metacells against single cells. Genes are colored by density.

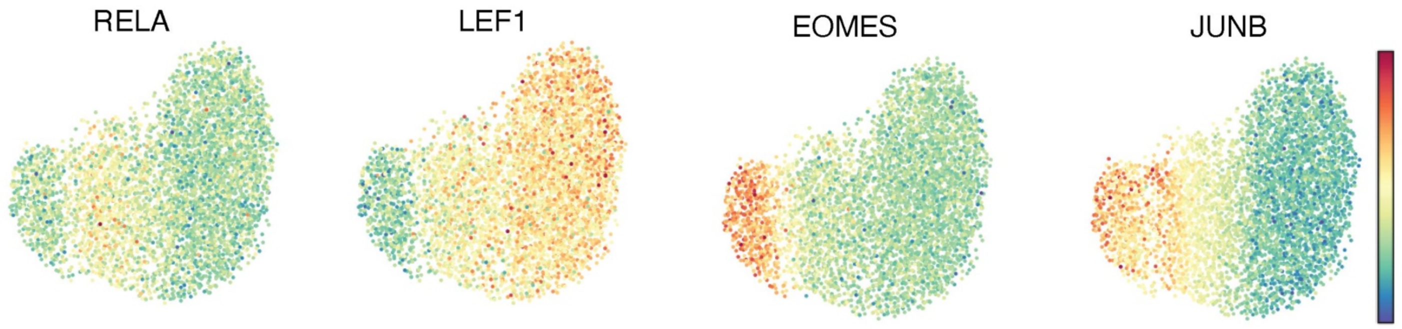


**Supplementary Fig. 10: Single-cell chromVAR scores for T-cell subsets**

UMAPs of T-cell subsets from PBMC multiome data (as in **Fig. 3E**) colored by single-cell chromVAR scores of key T-cell factors.

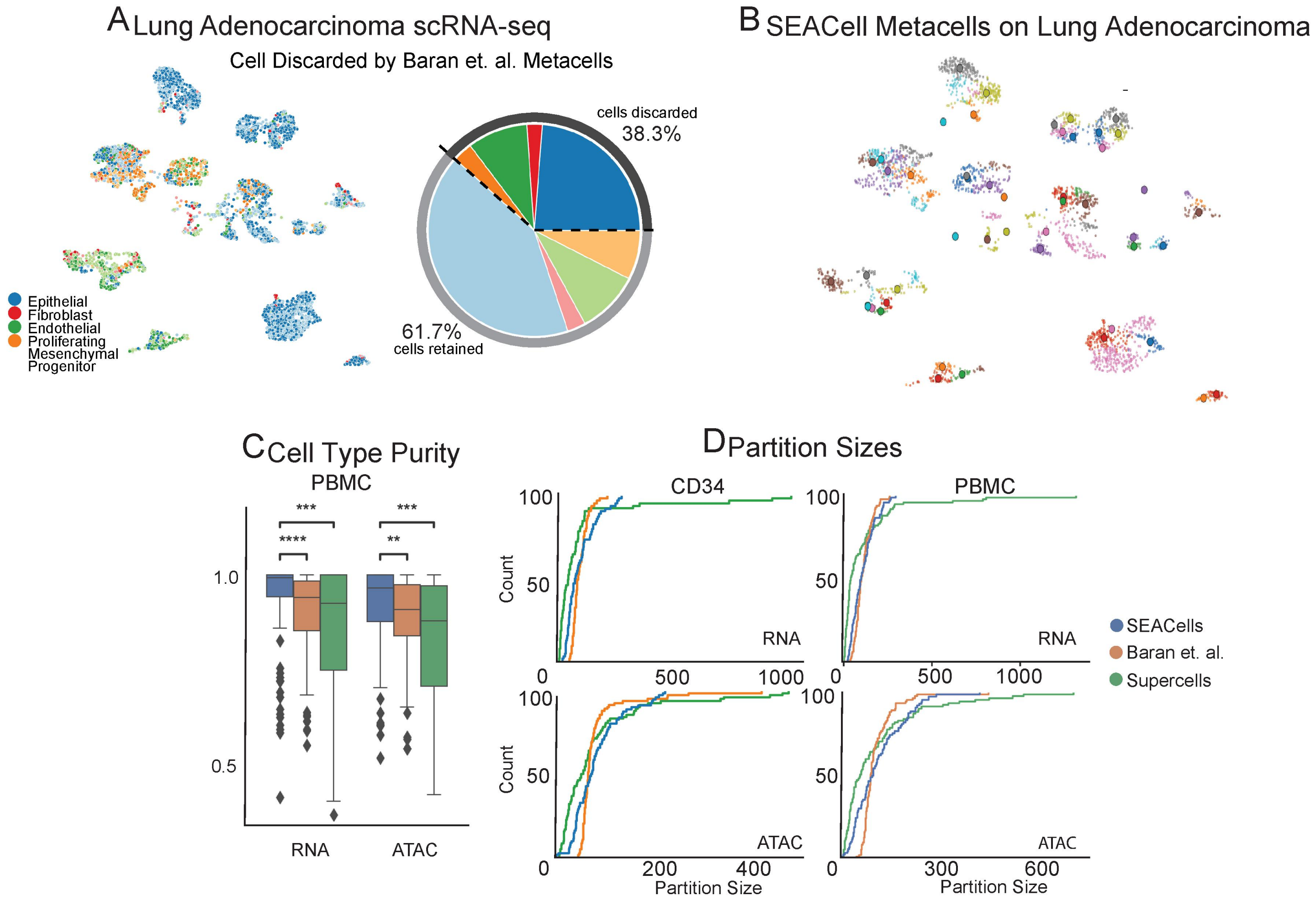


**Supplementary Fig. 11: Key characteristics of different metacell approaches**

A. UMAP of lung adenocarcinoma scRNA-seq data[1]. A large fraction of cells are pre-filtered by the MetaCell method[2] in this perturbed setting. Cells colored by cell type as derived by [ref].

B. SEACells metacells on the same lung adenocarcinoma dataset. No cells were discarded in this analysis.

C. Metacell cell-type purity (fraction of the maximally represented cell-type amongst the cells assigned to a metacell) computed by different methods on PBMC data. Wilcoxon rank-sum test was used to assess the significance of differences (** $0.001 < P < 0.01$, *** $0.0001 < P < 0.0001$, **** $P < 0.0001$).

D. Cumulative distribution plots showing the partition size or number of cells in each metacell. Super-cells produces metacells that are very large and span a large proportion of the high density regions.

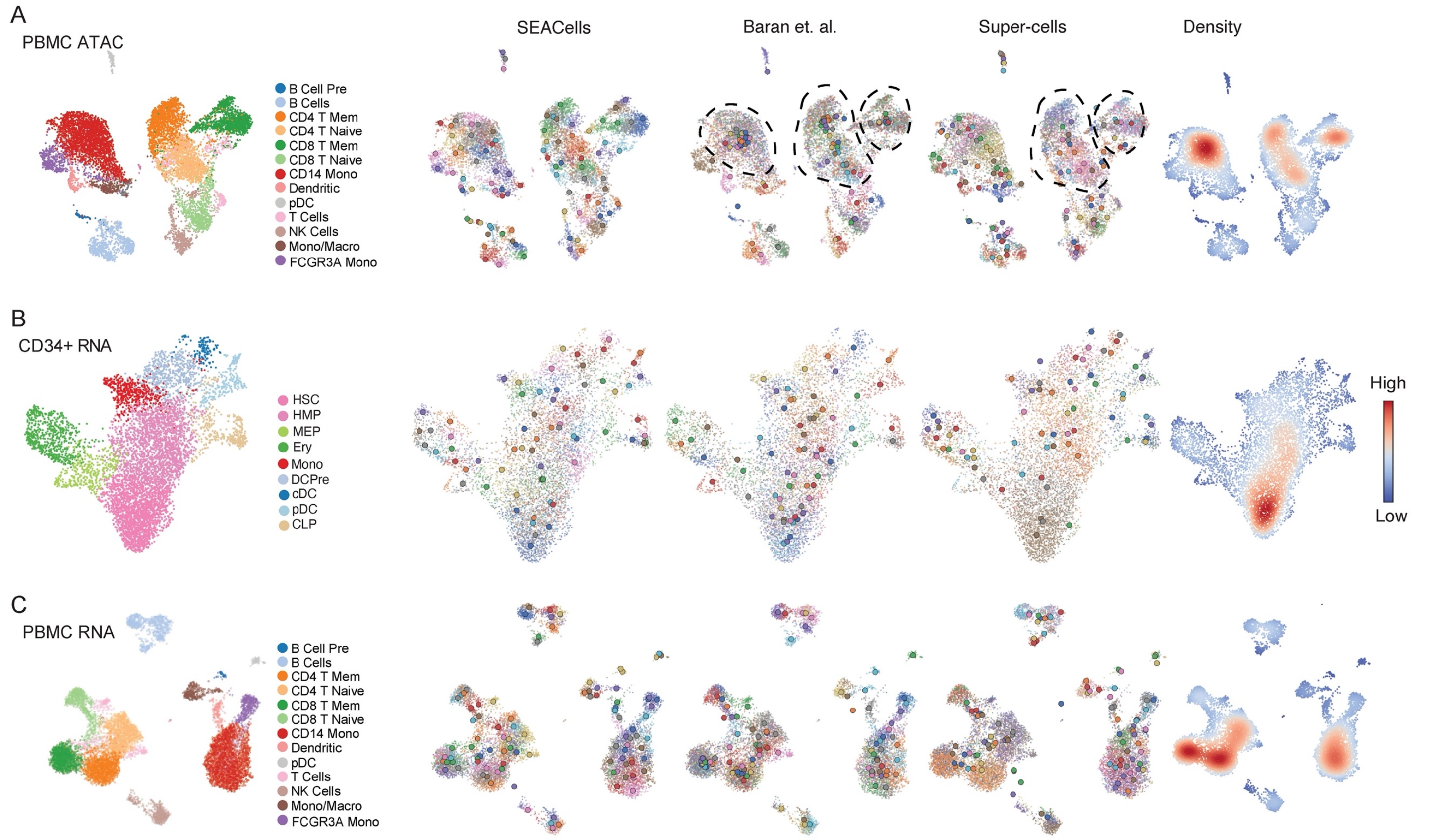


**Supplementary Fig. 12: Performance of different metacell approaches**

A. ATAC modality UMAPs of PBMCs (as in **Fig. 2B**), colored by metacells identified by the specified method or colored by cell density. Dots, cells; circles, metacells. Highlighted regions indicate a lack of well-defined metacells.

B. RNA modality UMAPs of $CD34^{+}$ bone marrow (as in **Fig. 2D**), colored by metacells or cell density.

C. RNA modality UMAPs of PBMCs (as in **Fig. 2A**), colored by metacells or cell density.

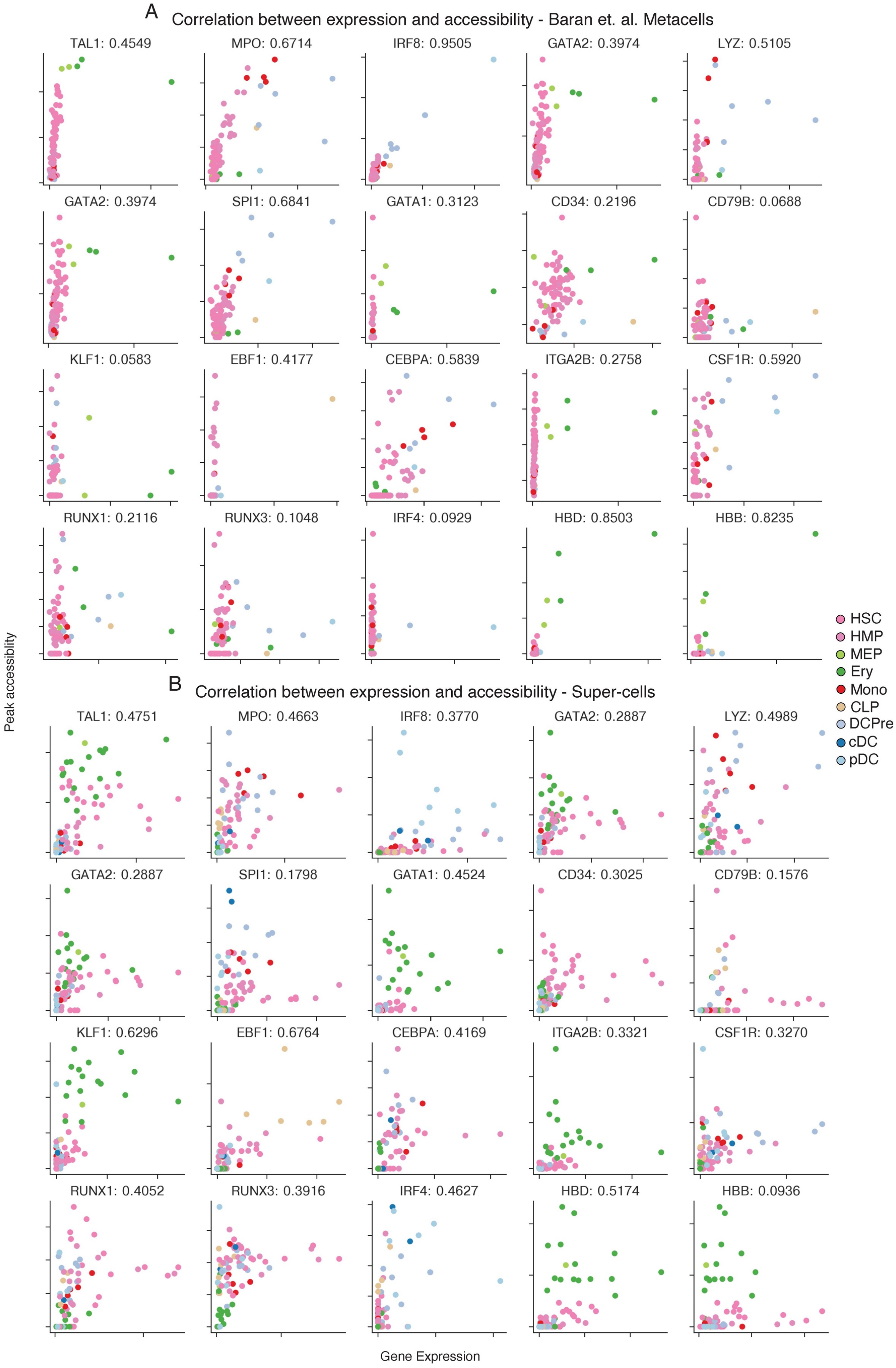
A
Correlation between expression and accessibility - Baran et. al. Metacells
TAL1: 0.4549
MPO: 0.6714
IRF8: 0.9505
GATA2: 0.3974
LYZ: 0.5105
GATA2: 0.3974
SPI1: 0.6841
GATA1: 0.3123
CD34: 0.2196
CD79B: 0.0688
KLF1: 0.0583
EBF1: 0.4177
CEBPA: 0.5839
ITGA2B: 0.2758
CSF1R: 0.5920
RUNX1: 0.2116
RUNX3: 0.1048
IRF4: 0.0929
HBD: 0.8503
HBB: 0.8235
Peak accessibility
HSC
HMP
MEP
Ery
Mono
CLP
DCPre
cDC
pDC
B
Correlation between expression and accessibility - Super-cells
TAL1: 0.4751
MPO: 0.4663
IRF8: 0.3770
GATA2: 0.2887
LYZ: 0.4989
GATA2: 0.2887
SPI1: 0.1798
GATA1: 0.4524
CD34: 0.3025
CD79B: 0.1576
KLF1: 0.6296
EBF1: 0.6764
CEBPA: 0.4169
ITGA2B: 0.3321
CSF1R: 0.3270
RUNX1: 0.4052
RUNX3: 0.3916
IRF4: 0.4627
HBD: 0.5174
HBB: 0.0936
Gene Expression

**Supplementary Fig. 13: Accessibility peaks and gene expression are not well correlated in MetaCell and Super-cells metacells**

A. Relationship between MetaCell[2] aggregated-gene expression and accessibility of the most correlated peak for key hematopoietic genes.

B. Same as (A), computed using Super-cells.

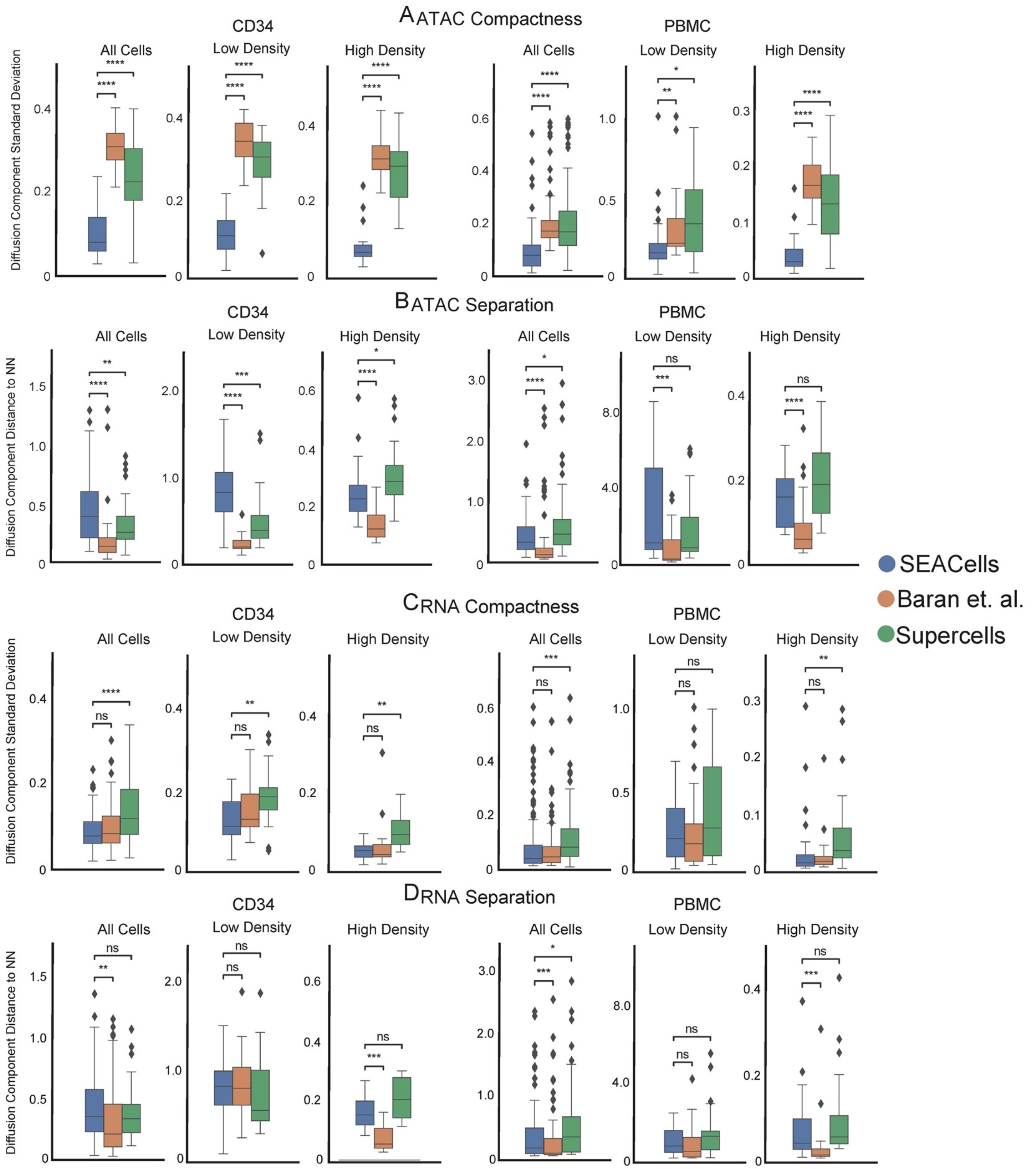
A ATAC Compactness
B ATAC Separation
C RNA Compactness
D RNA Separation
CD34
PBMC
All Cells
Low Density
High Density
Diffusion Component Standard Deviation
Diffusion Component Distance to NN
SEACells
Baran et. al.
Supercells

**Supplementary Fig. 14: Performance of different approaches in achieving metacell compactness and separation.**

A. Metacell compactness (average diffusion component standard deviation; Methods) measured in the ATAC modality of CD34 and PBMC multiome data. A lower score indicates more compact metacells.

B. Metacell separation (distance between nearest metacell neighbor in diffusion space; Methods) measured in the ATAC modality of CD34 and PBMC multiome data. Larger separation indicates better performance.

C. Metacell compactness measured in the RNA modality of CD34 and PBMC multiome data.

D. Metacell separation measured in the RNA modality of CD34 and PBMC multiome data. Larger separation indicates better performance.

Comparisons were carried out on all metacells, or metacells in low-density or high-density regions. Wilcoxon rank-sum test; ns: $P > 0.05$, * $0.01 < P < 0.05$, ** $0.001 < P < 0.01$, *** $0.0001 < P < 0.0001$, **** $P < 0.0001$.

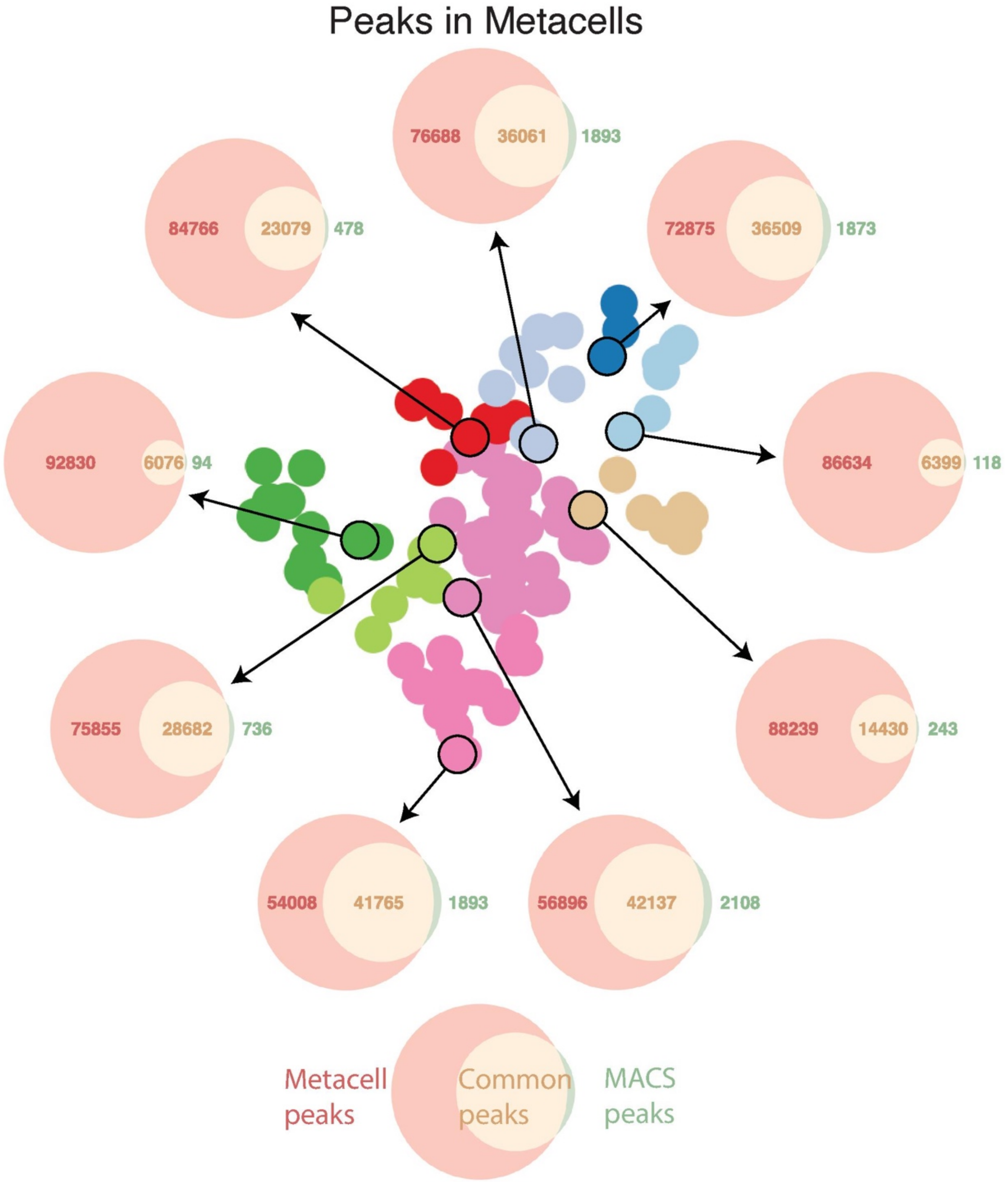


**Supplementary Fig. 15: Metacell peak calling**

Each Venn diagram depicts the number of peaks in a metacell based on *de novo* peak calling using MACS2 on the metacell ATAC fragments (green), open peaks called using Poisson statistics on ATAC fragments from all cells in the sample (red), and the intersection set (pink). *De novo* peak calling always leads to fewer peaks.

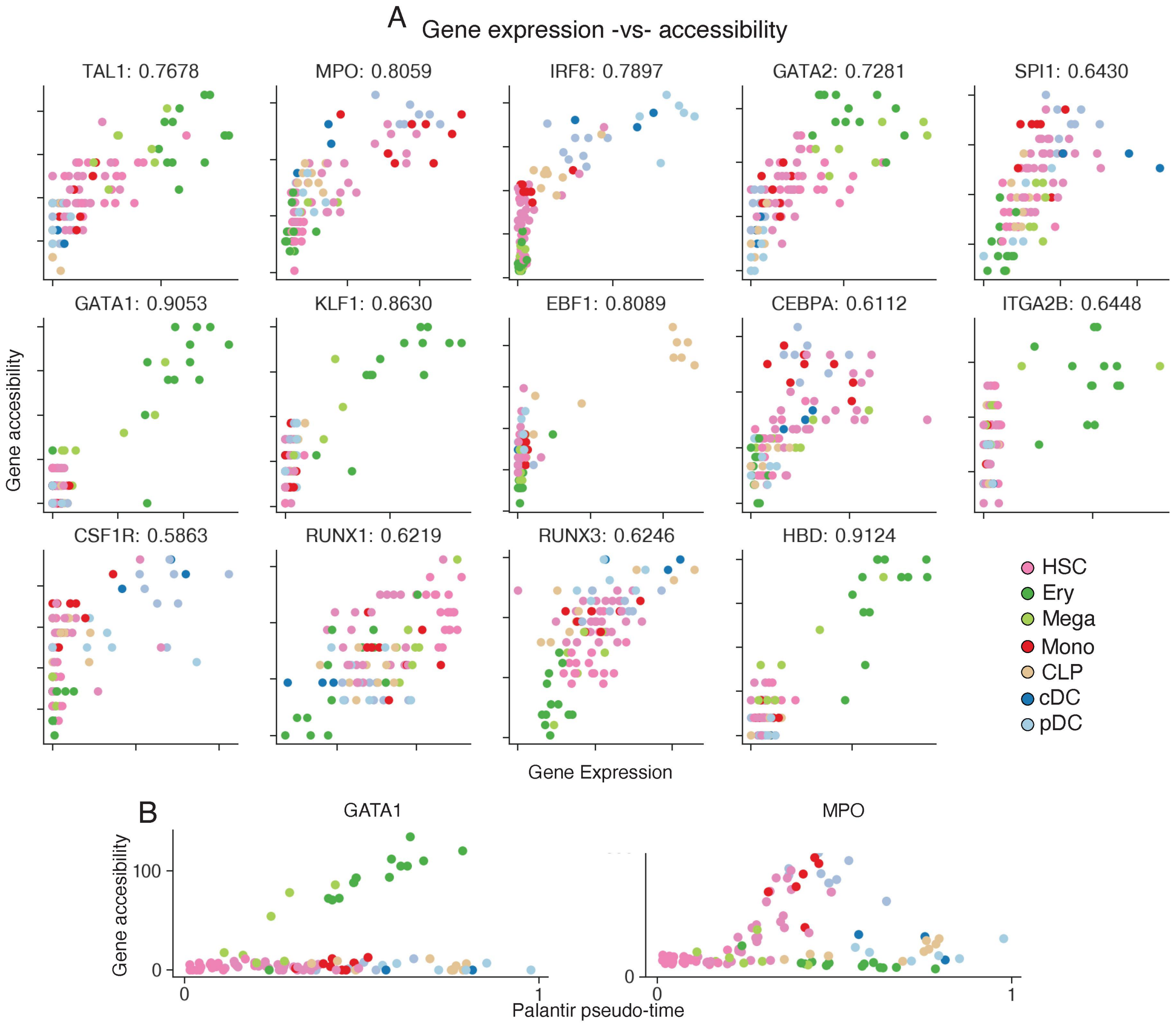


**Supplementary Fig. 16: Gene accessibility in CD34⁺ bone marrow**

A. Correlation between metacell-aggregated gene expression and gene accessibility scores for a selection of key hematopoietic genes. Spearman correlations computed using the CD34⁺ bone marrow multiome data are provided next to gene names.

B. Dynamics of gene accessibility scores for *GATA1* (left) and *MPO* (right). Each dot represents a metacell plotted along pseudotime (x-axis).

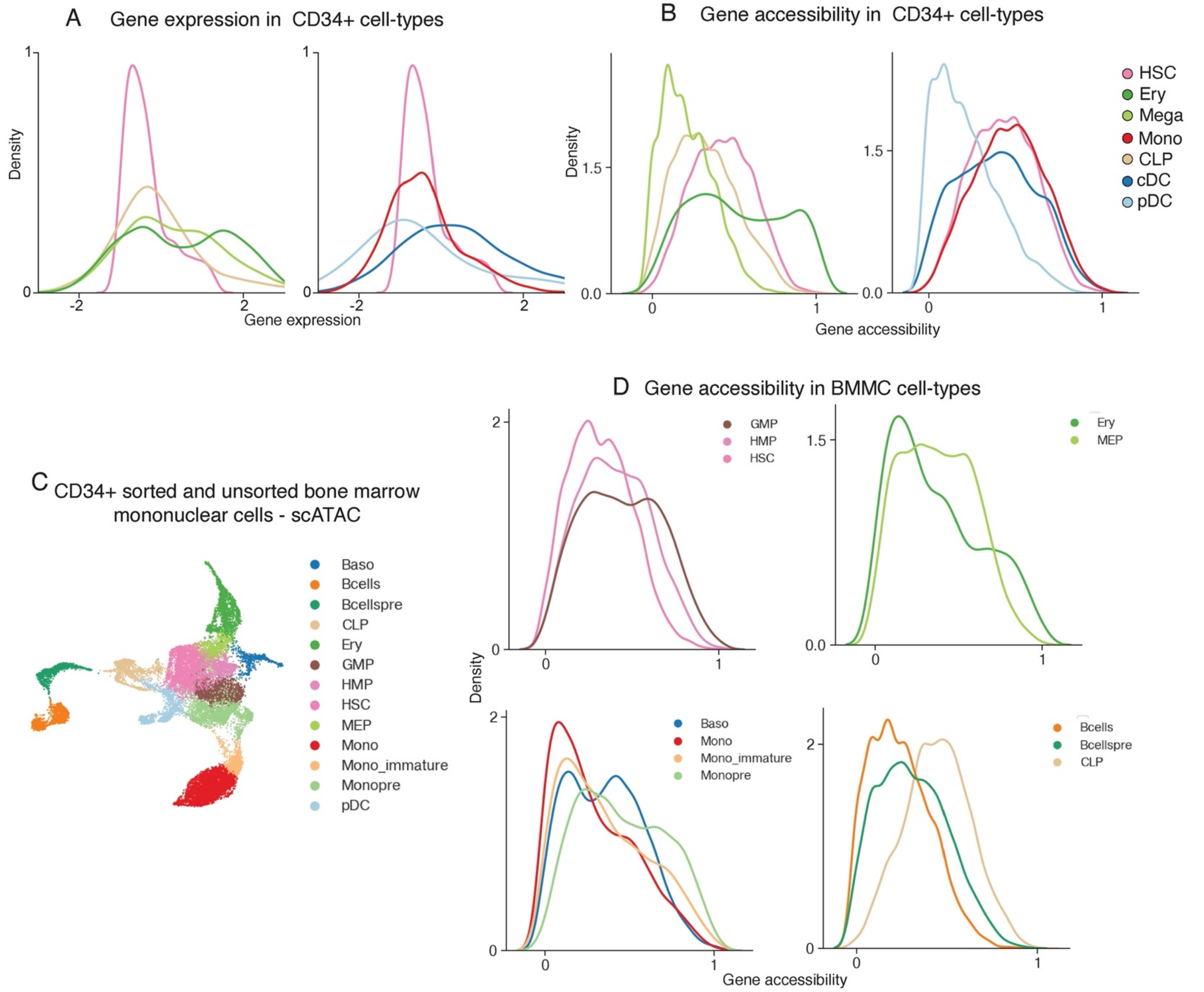


**Supplementary Fig. 17: Accessibility dynamics during hematopoietic differentiation**

A. Distribution of gene expression for all hematopoietic cell types, using CD34$^+$ bone marrow multiome data.

B. Distribution of gene accessibility for all hematopoietic cell types using CD34$^+$ bone marrow multiome data.

C. UMAP of an scATAC-seq dataset of CD34$^+$-sorted and unsorted bone marrow mononuclear cells (BMMCs).

D. Gene accessibility distributions for highly regulated genes in the BMMC dataset. Peak gene correlations were determined using the CD34$^+$ bone marrow multiome data, since only ATAC modality is available for the BMMC dataset.

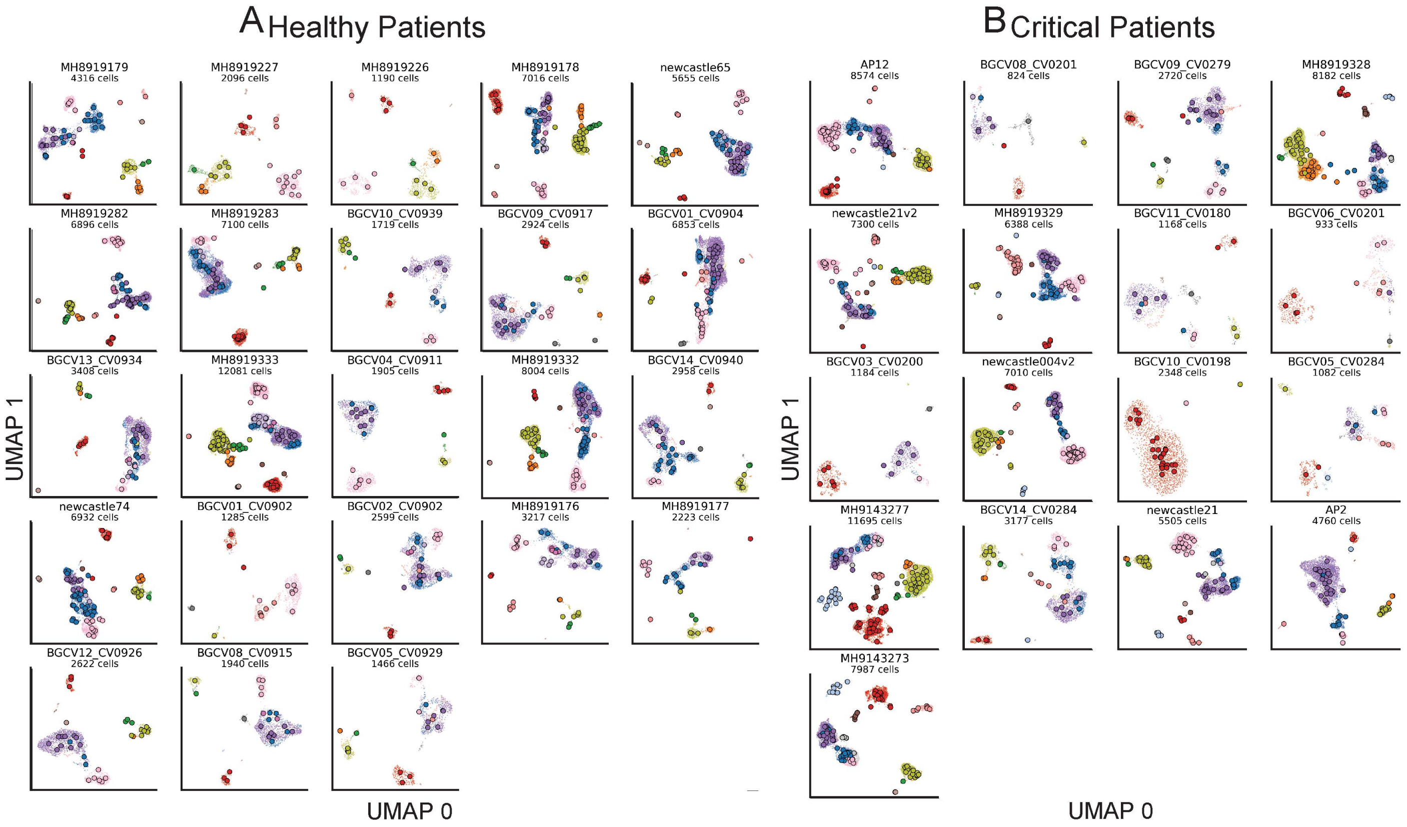


**Supplementary Fig. 18: SEACells metacells results in a COVID-19 cohort**

UMAPs showing SEACells results for healthy donors (A) and COVID-19 patients (B). Each plot represents a single individual.

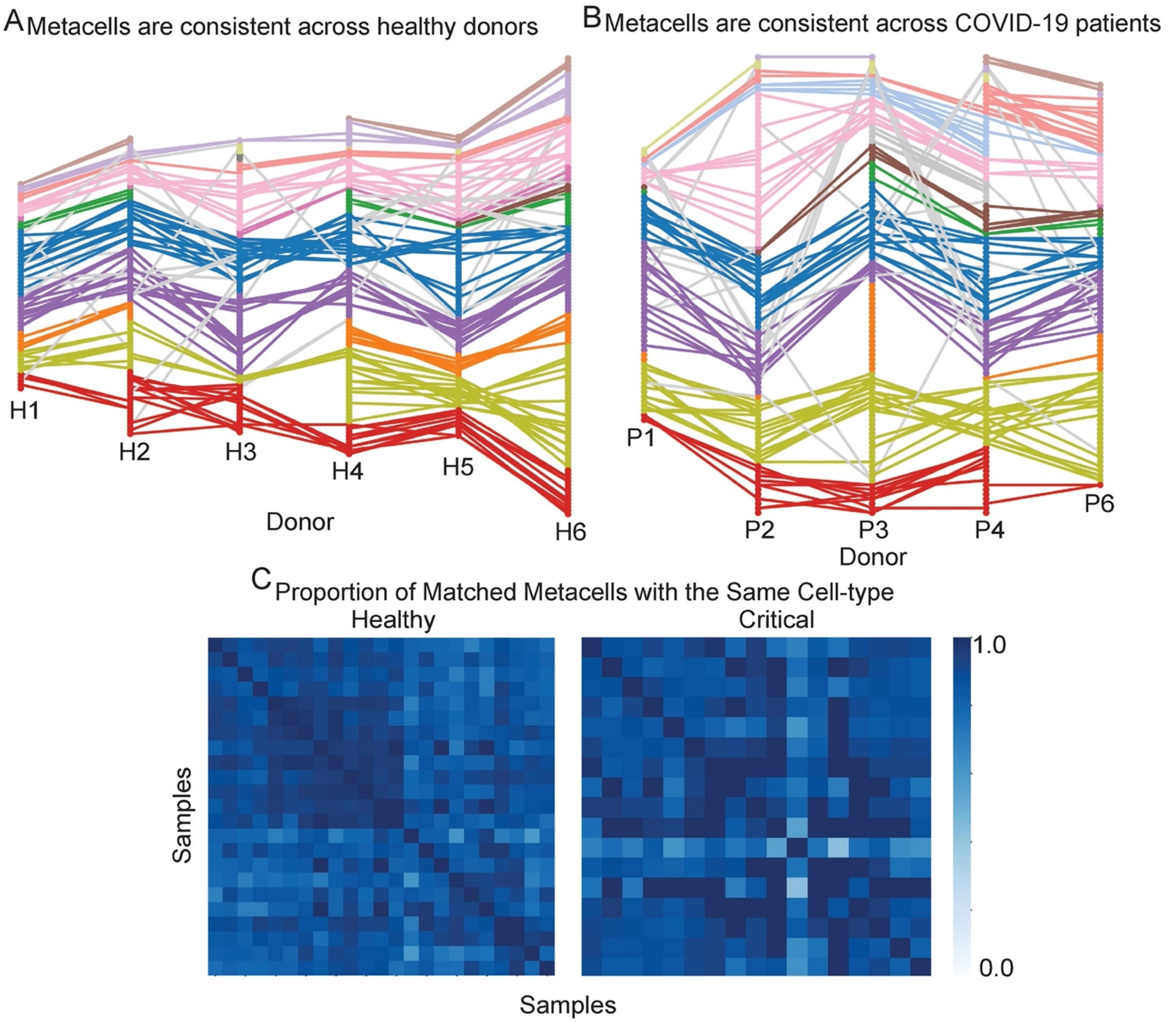


**Supplementary Fig. 19: Consistency of SEACells metacells among healthy and COVID-19 patients**

A. Metacell states are consistent and reproducible between pairs of healthy individuals. Mutually neighboring metacells between different pairs of healthy donors are connected by edges.
B. Same as (A), for COVID-19 patients.
C. Proportion of mutually neighboring edges that match to the same cell-type for healthy donors (left) and COVID-19 patients (right). On average, 88% of edges between healthy donors and 87% of edges between COVID-19 patients are consistent.

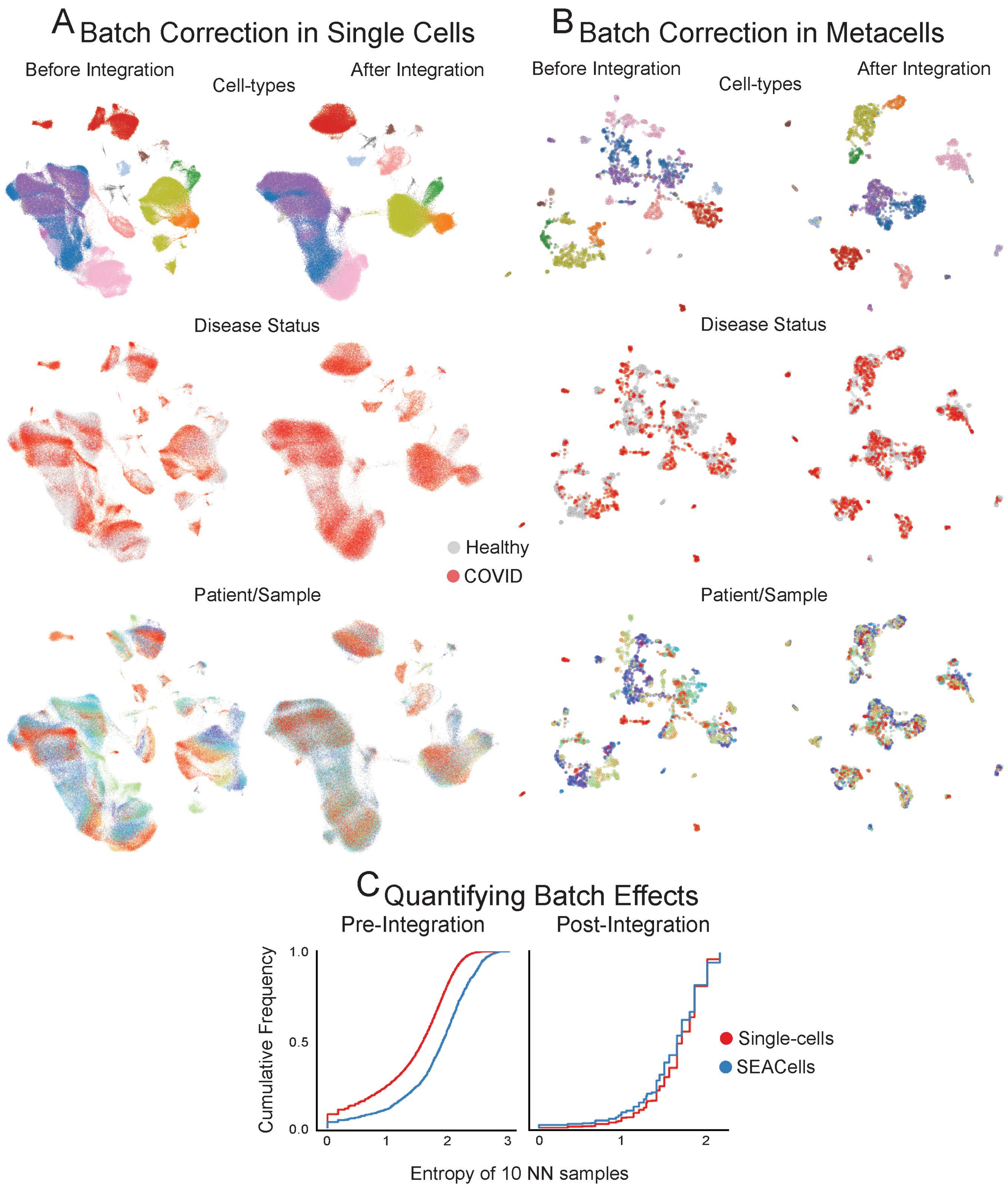


**Supplementary Fig. 20: Large-scale data integration using SEACells**

A. UMAPs showing single-cell data from healthy donors and critical COVID-19 patients before (left) and after (right) batch correction and data integration using Harmony[3].

B. Same as (A) for SEACells metacells instead of single cells.

C. Cumulative distribution showing the entropy of samples among 10 nearest neighbors before integration (left) and after integration (right).

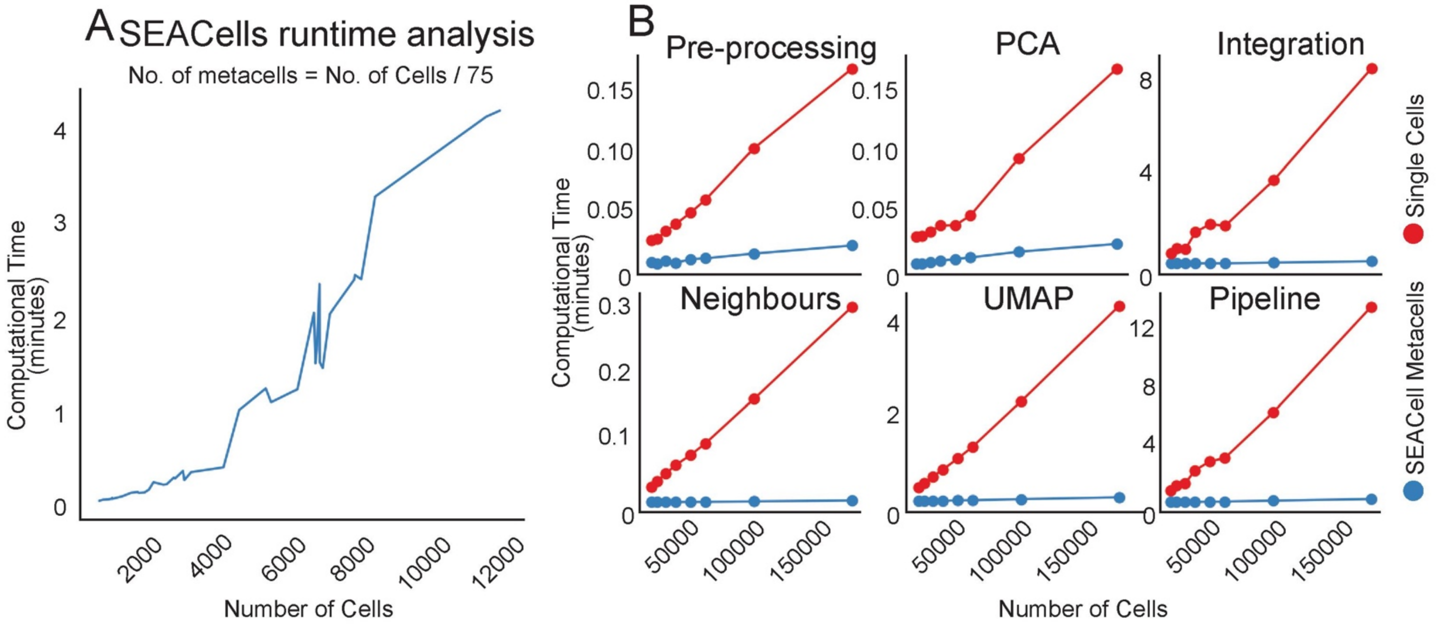


**Supplementary Fig. 21: SEACells runtime analysis**

A. Time to compute SEACells metacells, as a function of single-cell dataset size. The number of metacells was fixed as number of cells / 75.

B. Runtime comparison of SEACells metacells and single cells for different single-cell tasks. Preprocessing involves normalization and batch correction. Pipeline refers to runtime for all tasks shown.

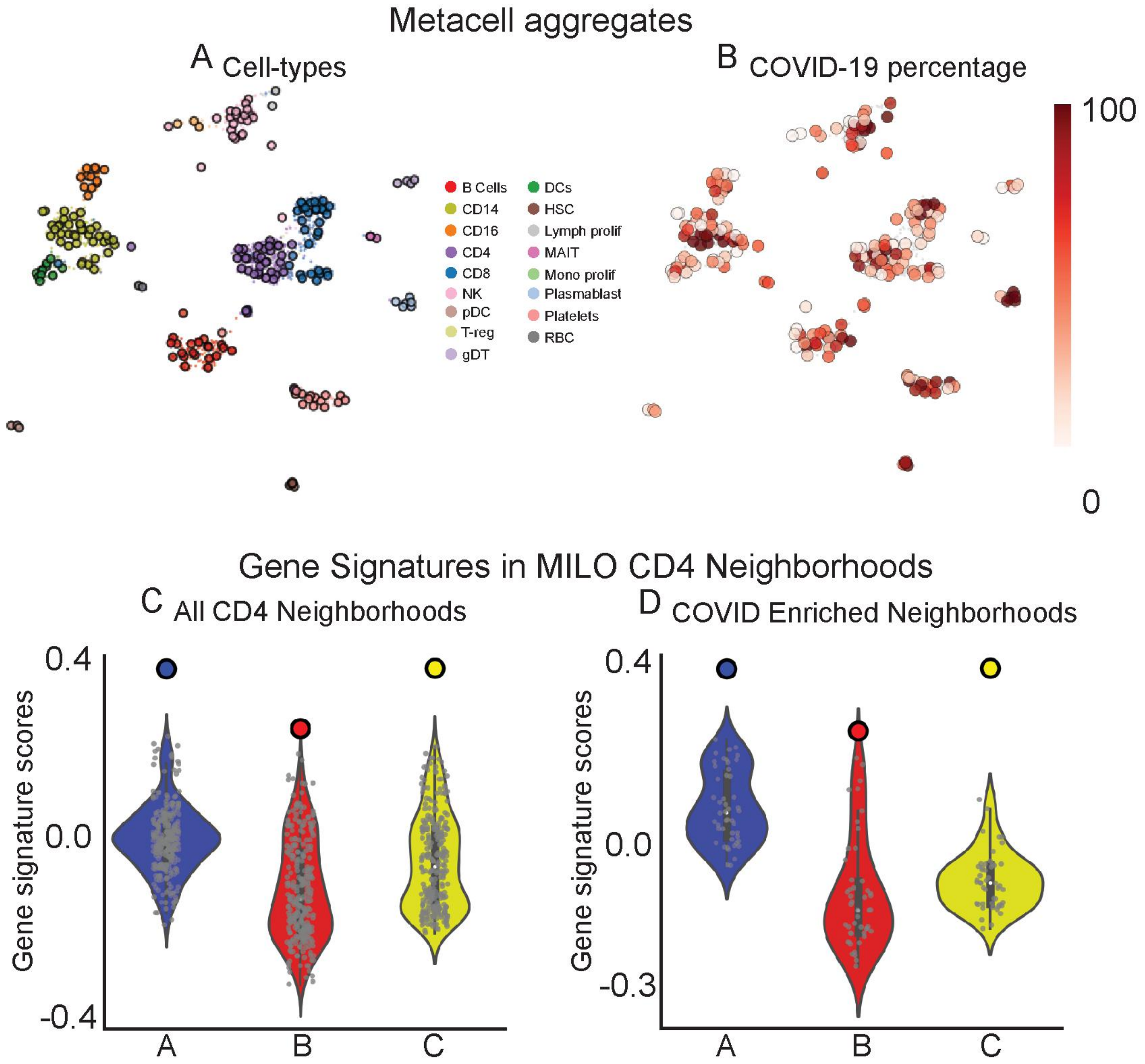


**Supplementary Fig. 22: Metacell aggregates enable comparison across conditions**

A,B. UMAPs showing Metacell aggregates (SEACells applied to metacells pooled from healthy and COVID-19 donors), colored by cell-type (A) and percentage of cells derived from COVID-19 patients (B). Metacell aggregates serve as input for differential abundance testing.

C,D. Gene signature scores for the CD4 metacells in Fig. 6D. Violin plots represent signature scores for MILO[4] neighborhoods, solid dots represent scores for the SEACells metacells. Scores are computed for all CD4 T-cell MILO neighborhoods (C) and for the subset of neighborhoods enriched in COVID-19 (D).

### References


1 Laughney, A. M. *et al.* Regenerative lineages and immune-mediated pruning in lung cancer metastasis. *Nat Med* **26**, 259-269, doi:10.1038/s41591-019-0750-6 (2020).

2 Baran, Y. *et al.* MetaCell: analysis of single-cell RNA-seq data using K-nn graph partitions. *Genome Biol* **20**, 206, doi:10.1186/s13059-019-1812-2 (2019).

3 Korsunsky, I. *et al.* Fast, sensitive and accurate integration of single-cell data with Harmony. *Nat Methods* **16**, 1289-1296, doi:10.1038/s41592-019-0619-0 (2019).

4 Dann, E., Henderson, N. C., Teichmann, S. A., Morgan, M. D. & Marioni, J. C. Differential abundance testing on single-cell data using k-nearest neighbor graphs. *Nat Biotechnol* **40**, 245-253, doi:10.1038/s41587-021-01033-z (2022).
